# Disentangling cellular heterogeneity into interpretable biological factors through structured latent representations

**DOI:** 10.1101/2024.11.06.622266

**Authors:** Amir Ali Moinfar, Fabian J. Theis

## Abstract

Deep generative models have become central to single-cell omics analysis, but their latent spaces remain difficult to interpret biologically. Linear factor models offer dimension-wise interpretability, but often lack the nonlinear flexibility, scalability, and integration quality required for large multi-batch atlases. We bridge this gap with Disentangled Representation Variational Inference (DRVI), an unsupervised deep generative model that learns dimension-wise interpretable representations for single-cell omics without supervised priors on cell types or biological processes. DRVI achieves this through additive decoder subnetworks combined with log-sum-exp pooling, enabling disentangled nonlinear gene programs. Across atlases, perturbation screens, and developmental datasets, DRVI separates cell identity, signaling pathways, stress responses, developmental trajectories, and perturbation effects into distinct interpretable factors. These factors identify rare migratory dendritic cells and recover coherent combinatorial perturbation programs in CRISPR screens. Systematic benchmarks show that this interpretability does not reduce integration performance. DRVI recovers biological factors more accurately while maintaining competitive integration quality. Altogether, DRVI provides a practical route to nonlinear single-cell modeling with factor-level interpretability.

## Introduction

Single-cell sequencing technologies profile transcriptomes and other omics levels at single-cell resolution in a high-throughput fashion [1, 2]. These readouts enable researchers to explore developmental processes, assess drug and perturbation effects, construct reference atlases, and compare organoids with primary tissues [3–10]. Among the many computational tools developed for single-cell analysis [11], deep generative models have proven effective for summarizing data within a latent space, learning low-dimensional representations that integrate data across batches and outperform classical approaches in large-scale benchmarks [12–14]. Several extensions now address diverse modalities and use cases [15–18].

Yet these latent spaces are typically entangled, with multiple unrelated biological processes intertwined within a single dimension [19], which limits their direct biological interpretability [20, 21]. In practice, the latent space is therefore used only as an intermediate for computing neighborhood graphs, on which visualization via UMAP [22, 23] and cell-type identification via Leiden clustering [24] are performed. While essential for single-cell analysis [25], clustering identifies the most dominant signals, typically cell types, and cannot capture continuous processes, shared programs across cell types, or secondary signals within the same cells. Linear factor models provide dimension-wise interpretable representations, but their linearity limits flexibility, scalability, and integration quality in complex multi-batch datasets [26]. Bridging the disentanglement capacity and interpretability of factor models with the flexibility and scalability of deep generative models remains an open challenge.

Disentangled representation learning offers a principled framework for this goal by assigning distinct sources of variation to separate latent dimensions, each representing a coherent biological process (Fig. 1a) [27]. In single-cell omics, supervised approaches leverage perturbation labels or predefined gene sets to guide latent assignments (e.g., sVAE [28], SAMS-VAE [29], expiMap [30], pmVAE [31], and Spectra [19]), but such auxiliary information is often unavailable for discovery tasks and, when available, can bias the analysis toward known priors. Theoretical work shows that learning such disentangled factors without supervision requires suitable inductive biases on both model and data [32]. For example, deep convolutional architectures (CNNs) enable unsupervised disentanglement in computer vision [33, 34]. In single-cell omics, factor models such as MOFA [35] and LIGER [36] offer dimension-wise interpretability through linearity, sparsity, and non-negativity as inductive biases, but lack the nonlinear flexibility and integration quality required for complex multi-batch data [26]. Deep generative models typically rely on unconstrained multilayer perceptrons lacking such structural biases. We address this limitation by combining additive decoders with a novel pooling as an inductive bias for single-cell omics. A comprehensive overview of dimension-wise and block-based (less relevant to our work) disentanglement methods, along with the methodological contributions of this work, is provided in Supplementary Note 1 and Supplementary Table 1.

**Figure 1:**
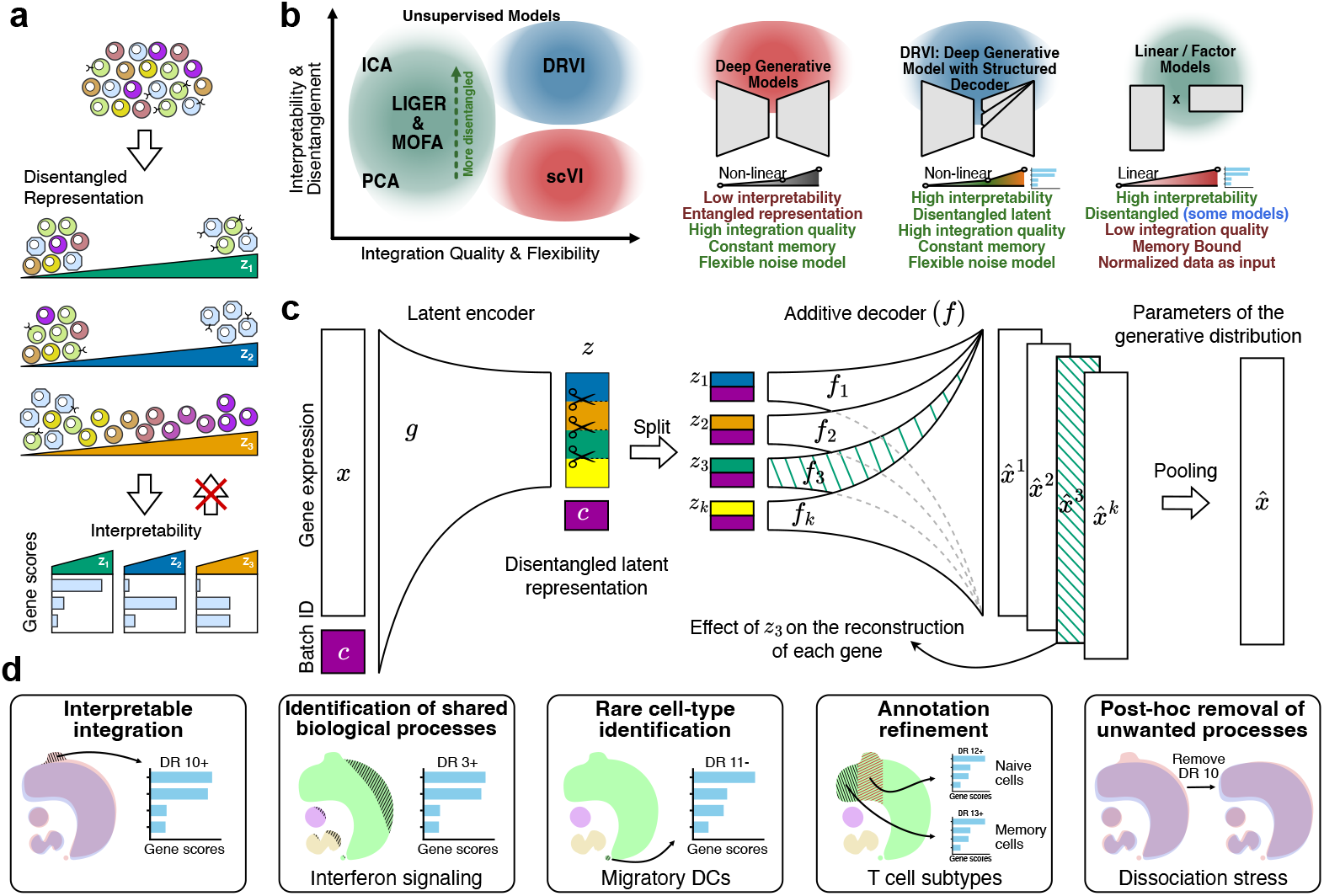
DRVI recovers interpretable axes of biological variation. **a**, Disentanglement concept. A disentangled latent model should encode concepts such as cell states, biological processes, and developmental stages in distinct latent factors that can be interpreted independently through their associated gene scores. Not every interpretable model solves disentanglement. **b**, Comparison of unsupervised representation-learning models based on integration quality, flexibility, interpretability, and disentanglement. Linear factor models (PCA, ICA, LIGER, MOFA) are interpretable but memory-bound and limited in integration quality and flexibility. Modeling constraints in ICA, LIGER, and MOFA encourage disentanglement. Deep generative models such as scVI integrate batches well but yield entangled, uninterpretable latent spaces. DRVI combines the flexibility and integration quality of deep generative models with the interpretability and disentan-glement of factor models. **c**, Schematic of the DRVI model. The model uses a nonlinear encoder to map gene expression and optional batch covariates to a latent representation, splits the latent space into factors, decodes each factor separately with additive decoder subnetworks, and pools the decoded outputs to form the parameters of the generative distribution. Since each factor is decoded separately, they can be interpreted independently.**d**, DRVI enables unbiased exploration of singlecell transcriptomics. It provides interpretable integration, identifies shared biological processes such as interferon signaling, isolates rare cell types such as migratory dendritic cells, supports annotation refinement, and transfers learned factors to unseen datasets. DR, disentangled representation.

We present Disentangled Representation Variational Inference (DRVI), an unsupervised deep generative model that learns interpretable nonlinear factors from single-cell count data. DRVI addresses the need for inductive bias in unsupervised disentanglement [32] and bridges the gap between factor models and deep generative models by combining additive decoders, which support latent-variable identification under theoretical conditions, with log-sum-exp (LSE) pooling function suited to singlecell omics. Unlike factorization models, whose log-space additivity implies multiplicative effects between overlapping programs in count space, LSE pooling represents overlapping programs additively in count space (Supplementary Notes 2 and 3).

Across atlas-scale datasets, perturbation screens, and developmental data, DRVI separates cell identity, signaling pathways, stress responses, developmental trajectories, and perturbation effects into distinct interpretable factors. In the immune dataset, it recovers a rare fibroblast population and refines dendritic-cell annotations, including a DC1 subset that baseline models do not identify. In the Human Lung Cell Atlas, DRVI disentangles interferon signaling, angiogenesis, proliferation, metal-ion stress responses, and rare populations beyond coarse cell types, including a chemokine-response program that remains entangled under baselines. In CRISPR screen data, it recovers coherent perturbation response programs for single and combinatorial perturbations. On a developmental pancreas dataset, it achieves better separation of cell-cycle stages and identification of terminal states than the baseline models.

The identified factors are consistent and transferable. Models trained independently on separate PBMC datasets recover matching gene signatures, and factors transfer to unseen cohorts while preserving disentanglement quality. Systematic benchmarks on six primary datasets and twelve additional multi-batch datasets from distinct organs confirm that DRVI consistently improves disentanglement while maintaining competitive integration quality.

## Results

### DRVI learns interpretable nonlinear latent factors

DRVI (Disentangled Representation Variational Inference) integrates datasets while learning unsupervised interpretable latent factors that can be read as biological processes from single-cell transcriptomics data. A representation is disentangled when distinct expression patterns, such as cell types and signaling pathways, occupy different axes, so each dimension can be read as its own coherent gene program rather than a mixture of several effects (Fig. 1a; see Methods and Supplementary Note 4 for formal definitions). The disentanglement concept is distinct from statistical independence of latent factors (Supplementary Note 5). Compared with factorization models and unconstrained autoencoders, DRVI is designed to combine nonlinear modeling, factor-level interpretability, and practical batch integration (Fig. 1b). While DRVI is built within the commonly used conditional variational autoencoders (CVAE) framework [12, 37], its decoder is fundamentally different from existing models. Specifically, it employs additive decoders followed by a pooling function to encourage the model toward representing distinct sources of variation in different latent factors.

The encoder *g* takes the (commonly) normalized gene expression of a cell *x* and optionally the covariate variable *c* — representing batch or other technical information to regress out, where both one-hot encoding [12] and embedding [38] are supported — as input and maps them to the latent space *z*. The latent representation *z* is split into factors that are decoded separately by additive subnetworks *f*_*k*_, and the resulting partial reconstructions are pooled to form the parameters of the generative distribution such as a Gaussian or Negative Binomial (Fig. 1c). Unlike the encoder, which employs a multi-layer perceptron (MLP) capable of approximating a wide range of functions, the additive decoder architecture [34] decodes latent dimensions separately, allowing for dimension-wise interpretability yet remaining more flexible than linear decoders. The decoder takes the latent space vector *z*, of dimension *K*, and splits it into *K/D* splits of dimension *D*, where we use *D* = 1 for the most powerful form of disentanglement by default. Each split, optionally together with the covariate variable *c*, is decoded separately using a nonlinear subnetwork *f*_*k*_ (where *k* ∈ {1, …, *K/D*}) back into the expression space 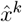 to later form the parameters of the generative distribution. Finally, the decoded vectors 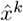 are aggregated using a potentially nonlinear pooling function, combining *K/D* vectors of the same size into one, to form the final generative distribution. The overall model is optimized based on the variational inference framework, with the encoder representing the approximate posterior and the decoder representing the generative model (see ‘The Generative Model’ in Methods).

DRVI aggregates decoded factors with log-sum-exp (LSE) pooling: 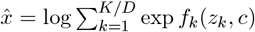. To assess the contribution of this pooling choice, we also consider DRVI with average pooling (DRVI-AP), which simply averages the decoded parameters. In practice, LSE pooling is essential for effective disentanglement in DRVI. Compared with linear models in log-transformed space (e.g., PCA, ICA, and MOFA), which implicitly assume that overlapping but distinct biological processes multiply in gene count space (Supplementary Note 2), LSE pooling has the mechanistic interpretation that distinct but overlapping biological processes combine additively in gene count space (justified in Supplementary Note 3).

Because each latent factor is decoded independently into expression space, its partial reconstruction can be interpreted on its own to define a gene program (Fig. 1c). To interpret these factors, we traverse each latent factor and inspect the affected features (e.g., genes in transcriptomics). Since the effect changes nonlinearly as we traverse the factor, we consider the maximum effect for each factor (see ‘Interpretation and ordering of learned factors’ in Methods). We refer to the genes identified by the interpretability pipeline as ‘top relevant genes’ for that factor. To interpret the factors beyond the genes used for model training, one can perform differential expression (DE) analysis.

Once we identify the top relevant genes, we can determine some programs through supervised information or external tools, such as existing annotations and gene-set enrichment analysis (GSEA). Programs can also be contextualized via literature search, marker atlases, automated tools, or expert review. So the identification procedure is not limited to GSEA analysis. Because such external information is not supplied during training, it cannot bias the fitted representations.

In the following, we show that DRVI combines atlas-scale integration with dimension-wise interpretability. Its factors reveal shared biological processes, specific or rare cell types, and finer subtypes that can be annotated through their top genes. Because these signals are separated into individual factors, they can be interpreted, compared, and transferred across related datasets (Fig. 1d).

### DRVI identifies biological signals beyond cell types

To introduce the types of biological concepts that can appear as DRVI’s interpretable latent factors, we first applied DRVI with 32 latent factors to an immune dataset [26] composed of 9 batches from four human peripheral blood and bone marrow studies, with 16 annotated cell types (Fig. 2a, Supplementary Fig. 1b).

**Figure 2:**
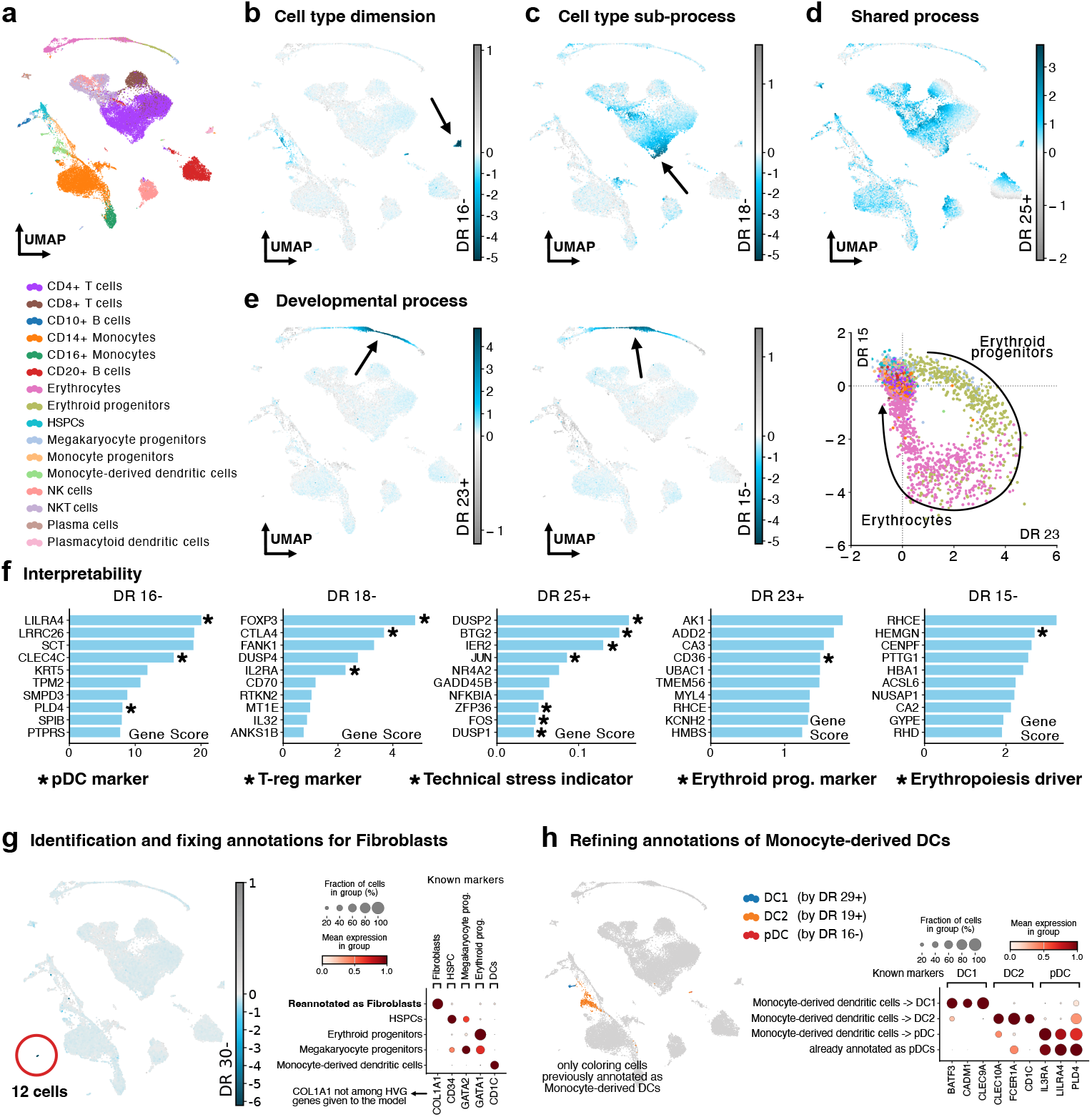
DRVI provides unbiased disentangled representation in Immune data. **a**, Uniform manifold approximation and projection (UMAP) of the immune dataset integrated by DRVI colored by cell types. **b**, Cell-type factor (DR 16-) identifying plasmacytoid dendritic cells (pDCs). **c**, Cell-type sub-process (DR 18-) corresponding to regulatory T-cells (T-regs). **d**, Shared process (DR 25+) active across cell types, representing technical stress. **e**, Developmental process captured by DR 23+ and DR 15-along the erythroid lineage, with a scatterplot of the two factors. The arrow is for visualization and is not derived from the model. **f**, Top identified genes for each example factor. Known marker genes are marked with stars. **g**, Identification of fibroblasts in the immune data. We identified and fixed the annotation of 12 fibroblast cells separated by DR 30-. We verified this subset using COL1A1, a fibroblast marker that was not present in the HVGs given to the model. **h**, For cells originally annotated as monocyte-derived dendritic cells, we used DR 29+ (DC1), DR 19+ (DC2), and DR 16-(pDC) to refine annotations. Cells originally marked as pDC are not updated. The dotplot of known marker genes indicates successful annotation refinement. DC, dendritic cells; pDC, plasmacytoid dendritic cells; T-reg, regulatory T-cells; prog., progenitors.

DRVI ranks the latent factors based on their impact on reconstruction, labeled as DR 1, DR 2, etc. (see ‘Interpretation and ordering of learned factors’ in Methods). As with many latent-factor models, such as ICA and PCA, positive and negative directions along an axis have no intrinsic meaning. Therefore, interpretation must consider both the dimension and direction (e.g., DR 1+ or DR 1-): values close to zero indicate weak factor activity, whereas larger magnitudes in either direction indicate stronger activity of the corresponding program.

We highlight example factors representing a cell type, a cell-type sub-process, a shared process, and two consecutive stages in a developmental process (Fig. 2b-e). The top genes related to each factor are illustrated in Fig. 2f and Supplementary Fig. 29, and the geometry formed by the developmental process is shown in a scatterplot (Fig. 2e).

DR 16-identifies plasmacytoid dendritic cells (pDCs), which are distinguishable by the expression of CLEC4C, LILRA4, and IL3RA [39, 40]. At a finer level, the expression of FOXP3, CTLA4 (CD152), and IL2RA (CD25) in DR 18-cells (Fig. 2c,f) suggests that this factor targets regulatory T-cells (T-regs) [41, 42].

DRVI can assign shared variation to dedicated latent factors. DR 25+ is active across multiple cell types (Fig. 2d), and BTG2, IER2, DUSP1, FOS, JUN, and ZFP36, all within the top relevant genes, have been previously reported as dissociation- or heat-induced stress-response genes in single-cell transcriptomic data [43, 44]. Aligned with previous findings, this technical response is cell-type dependent [44, 45].

Factors DR 23+ and DR 15-represent two consecutive stages in the developmental lineage of erythrocytes (Fig. 2e). The scatter plot illustrates an increase in DR 23 followed by a delayed decrease (increase in magnitude) of DR 15 in erythrocyte progenitors. As erythrocyte progenitors transition to mature erythrocytes, DR 23 decreases to zero while DR 15 reaches its maximum magnitude. Finally, DR 15 increases back to zero as the cells leave this developmental stage. Looking at the relevant genes, CD36 for DR 23+ is a known marker of erythrocyte progenitors [46]. For DR 15-, HEMGN (EDAG) is a key driver of erythropoiesis [46, 47], and RHCE encodes the RhD antigen in the red blood cell (RBC) membrane [48] (Fig. 2f).

A practical application of DRVI in disentangling cell types is annotation refinement and rare cell type identification. For instance, DR 30-revealed an underrepresented population of 12 fibroblast cells (Fig. 2g and Supplementary Fig. 1g-j), which were previously misannotated as HSPCs, monocyte progenitors, or erythroid progenitors. The validity of this revised annotation was confirmed using COL1A1, a known fibroblast marker not included in the model’s HVG input (Supplementary Fig. 1k). DRVI also guides users to potential misannotations. We used DR 16-, DR 29+, and DR 19+ to reannotate pDC, DC1, and DC2 subtypes of dendritic cells, respectively (Fig. 2h and Supplementary Fig. 1c-f). We validated this refinement using curated markers of these subtypes (Supplementary Fig. 1l).

### DRVI allows atlas-level identification of biological processes

To evaluate DRVI in a complex atlas-scale setting, we applied it to the Human Lung Cell Atlas (HLCA) [7]. We used the curated core of this atlas, comprising 166 samples from 14 datasets and 585K cells with curated finest-level annotations across 61 distinct cell identities.

We trained DRVI with 64 latent factors (Fig. 3a). For interpretation, we first excluded 12 vanished factors with near-zero values across all cells; we later return to the phenomenon of vanishing dimensions. We then identified 31 latent factors corresponding to cell types in the finest-level annotations. These factors show a favorable one-to-one correspondence with many cell identities (Fig. 3b top), in contrast to the weaker correspondences observed for benchmarked methods below. Disentanglement metrics confirm this observation quantitatively. The remaining 21 factors represent candidate axes for biological processes beyond cell identity (Fig. 3b bottom). Because the positive and negative directions of some factors capture distinct processes, including 4 cell-type factors that encode a biological process in the other direction, we analyze the two directions separately.

**Figure 3:**
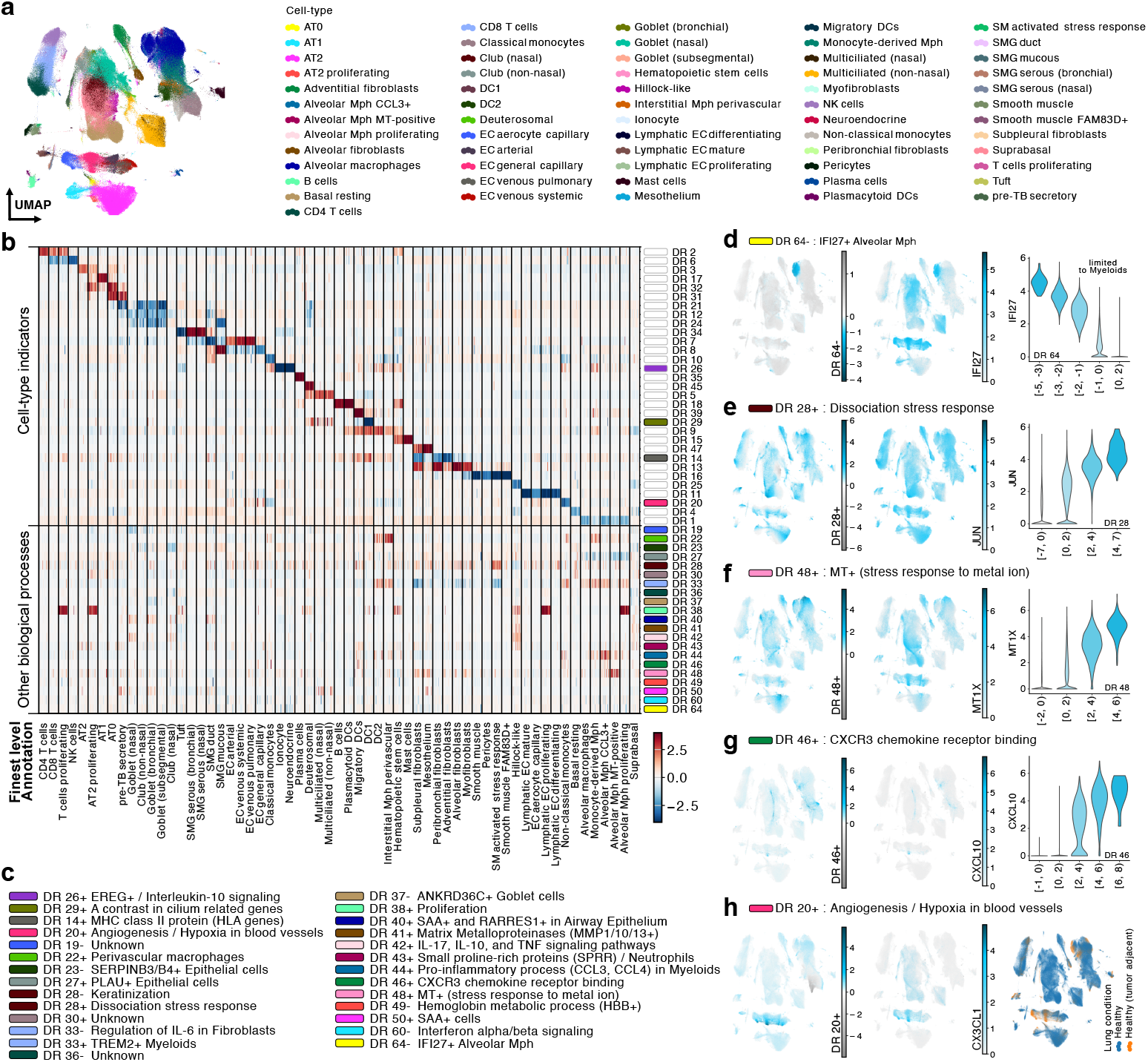
DRVI uncovers the variations in HLCA beyond cell types. **a**, UMAPs of the HLCA integrated by benchmarked methods. Cells are colored by finest-level annotations. **b**, DRVI latent factors. The heatmap illustrates each factor’s activity in cells vertically grouped by the finest-level annotations. Cells are subsampled to have an equal number of cells in each cell type. The factors are horizontally grouped into two categories: factors indicating cell-type (top) and factors indicating other biological processes (bottom). Factors representing non-cell-type processes are color-coded on the right-hand side of the plot. Vanished factors are omitted. For better illustration, some tail values are clipped. **c**, For each factor indicating a non-cell-type biological process, a short description is provided. **d-g**, Demonstration of five example factors. Left: Activity of the example factors on the UMAP. Middle: The expression of a relevant gene on the UMAP. Right: Violin plots indicating the expression levels of relevant genes versus the activity of factors. **h**, The activity of DR 20+ and one of the identified genes in the nonlinear gene program. This factor indicates angiogenesis and hypoxia in endothelial cells and is highly active in some tumor-adjacent cells. Mph, Macrophage.

Among the cell-type indicator factors, several are selectively activated in specific identities, including AT1 cells (DR 17+), plasma cells (DR 35+), DC1 cells (DR 39-), and migratory DCs (DR 39+). Other factors capture broader categories, such as macrophages (DR 1-), ciliated cells spanning deuterosomal and multiciliated cells (DR 5+), and transitional club-AT2 cells spanning AT0 and pre-TB secretory cells (DR 31+) [7] (Fig. 3b). Because disentanglement metrics already quantify cell-type identification, we do not evaluate this aspect further here. Notably, DRVI identifies most challenging cell types highlighted by the HLCA study, including migratory DCs (DR 39+), AT0 cells (DR 32+), hillock-like epithelial cells (DR 25-), and pre-TB secretory cells through the transitional club-AT2 factor (DR 31+). The pre-TB result is consistent with the HLCA study, where these cells were described through marker genes corresponding to transitional club-AT2 cells (not their own specific marker genes) [7]. Other methods often fall short in identifying these cell types, particularly migratory DCs, which only DRVI can capture along its latent factors (Fig. 5b). Identification of these cell types otherwise requires multiple clustering iterations [7] or very high-resolution clustering with hundreds of clusters; for example, identifying migratory DCs with scVI requires a resolution yielding 161 clusters (Supplementary Fig. 2). DRVI may therefore facilitate systematic discovery and seed annotation of novel or rare cell types in future cell atlases.

Although high-quality annotations facilitate interpretation, cell-type labels are not required to learn or identify cell-type factors. Users can instead annotate these factors post hoc by matching their interpreted signatures to marker atlases or literature, as in the immune example. Because this information is not provided to the model, label quality cannot bias the learned latent space.

For non-cell-type factors, we applied the interpretability pipeline to identify top relevant genes (Supplementary Figs. 31 and 30) and annotated the corresponding processes using existing literature and available tools. This yielded biological descriptions for 22 of the 25 factors expected to represent biological processes (Fig. 3c). Below, we detail five example factors (Fig. 3d-h); the remaining biological factors are discussed in Supplementary Note 6. To support these annotations, supporting evidence from the published literature is provided.

The five example factors illustrate how DRVI separates cell-state variation, shared stress programs, and overlapping biological processes in the HLCA (Fig. 3d-h). DR 64-captures an IFI27-associated alveolar macrophage (AM) program, consistent with reported IFI27+ AM subclusters [49–51]. Despite the expression of IFI27 in other cell types, such as blood vessels, DRVI has nonlinearly localized its expression to AMs due to the distinct patterns of co-regulated genes. DR 28+ represents a broad dissociation-stress response marked by ATF3, EGR1, JUN, FOSB, and FOS [43, 44]. DR 48+ reflects metallothionein (MT) gene family upregulation and metal-ion response; its overlap with DR 64-in AMs shows that DRVI can represent parallel, non-disjoint processes rather than forcing them into mutually exclusive clusters or statistically independent components. DR 46+ marks a CXCL9/CXCL10/CXCL11 chemokine-response program shared across cell types and associated with CXCR3 receptor binding. This factor is particularly relevant in the context of lung diseases [52, 53]. Finally, DR 20+ is primarily active in endothelial cells and is associated with angiogenesis and hypoxia-related genes. The upregulation of top relevant genes, including CX3CL1, SERPINE1, AKAP12, GRP4, and FSTL3, under hypoxic conditions [54–58], a common phenomenon observed in tumor-adjacent environments [59], indicates a potential role in tumor growth. The increased abundance of tumor-adjacent endothelial cells in the higher end of DR 20+ further supports this interpretation (Fig. 3h and Supplementary Fig. 3).

Altogether, DRVI disentangles interpretable sources of variation within the HLCA, a large-scale cell atlas, facilitating the identification of rare cell types and underlying biological programs. These factors capture inflammatory responses, signaling pathways, and variation that can be shared across related cell types or localized to specific populations, depending on their gene expression signatures. This allows scientists to study variations at a significantly finer resolution.

### DRVI’s learned factors saturate when overparameterizing the latent space

The number of biological processes, and therefore the required latent dimensionality, is usually unknown. We therefore examined overcomplete DRVI models, in which the number of latent dimensions *K* exceeds the number of processes needed to explain the data (Supplementary Fig. 4a). We define a factor as vanished when its maximum absolute value is less than 1. Consistent with other KL-regularized models [60], unused dimensions tended to vanish (Supplementary Fig. 4b). After saturation, increasing *K* left disentanglement, measured by LMS-SMI (see ‘Metrics’ in Methods), and integration quality, measured by total scIB score, largely unchanged (Supplementary Fig. 4c,d). The number of non-vanished factors depended on data complexity, but saturation of the number of non-vanished factors, the disentanglement metric, and integration quality was consistent across datasets (Supplementary Figs. 4e-g and 5). This behavior provides a practical strategy for choosing latent dimensionality when the true number of biological processes is unknown: starting from an expected latent dimensionality and increasing it until most additional dimensions vanish.

### The identified programs are consistent and transfer across datasets

To demonstrate the consistency and generalizability of DRVI, we individually trained four DRVI models, each with 128 latent factors, on four Human PBMC datasets in CellHint Blood data (Supplementary Fig. 6a). These datasets comprise a total of 336k cells with expert-curated annotations [61]. We observed that the number of non-vanished factors ranged from 31 to 43 (Supplementary Fig. 6b,c), with the variability likely stemming from minor biological and technical differences between the datasets.

We assessed the consistency of the identified programs across datasets using the Rank-Biased Overlap (RBO) similarity score [62] (see ‘Metrics’ in Methods). For comparison, we also trained PCA, ICA, LIGER, MOFA, and scETM on the same four datasets. DRVI and LIGER showed high RBO among same-cell-type factors and low RBO among different-cell-type factors, indicating consistent and specific gene signatures across datasets (Supplementary Fig. 7a,b). We next evaluated transferability by applying each model trained on one Blood dataset to the remaining datasets. DRVI preserved the highest disentanglement quality under this transfer setting (Supplementary Figs. 7c,d).

### DRVI disentangles cell-cycle information and identifies developmental stages

We applied DRVI and other benchmarked methods to single-cell mouse samples of the developmental pancreas at E15.5, comprising 3,696 cells across 15 finest-level annotated states (Fig. 4a and Supplementary Fig. 8a). We first considered the cell cycle as a well-studied biological process. Although cell-cycle variation can confound some integration tasks, unsupervised dimensionality reduction methods should retain this signal, and disentanglement methods should isolate it into identifiable factors. In practice, isolating such factors allows researchers to analyze the remaining dimensions with reduced interference from confounding variation.

**Figure 4:**
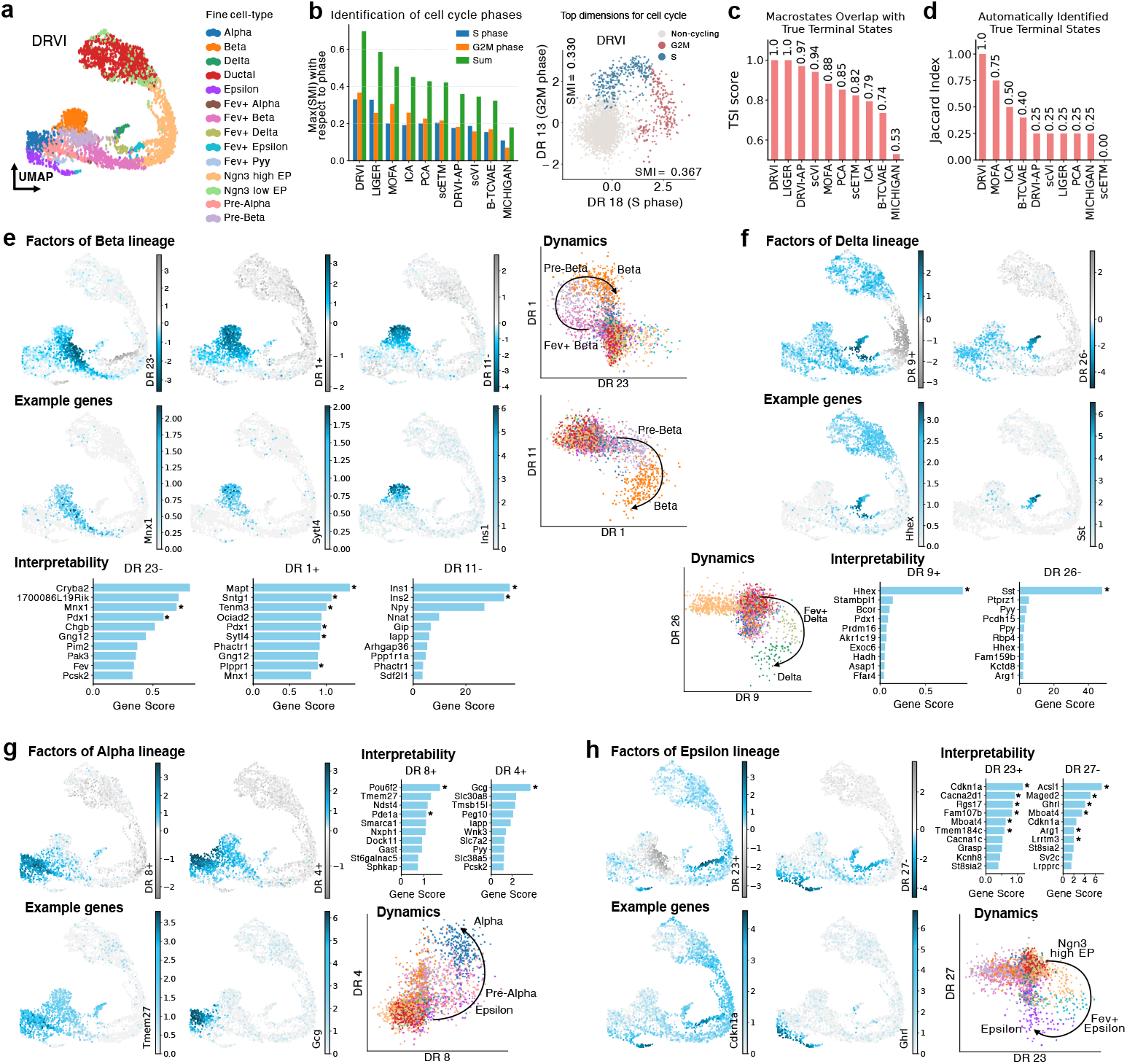
Identification of cell cycle and developmental stages by DRVI. **a**, UMAP of DRVI latent space for the developmental pancreas data. Cells are colored by finest-level annotations. **b**, The barplot shows the SMI score for the factors maximizing the SMI with respect to the S (blue) and G2M (orange) cell cycle phases in each benchmarked model. The sum of the two scores is provided in green. The scatterplot on the right shows DRVI factors representing these two phases, with points colored by the dominant cell cycle phase. Cells are assigned to the non-cycling category if both S and G2M scores are below 0.2. Otherwise, they are assigned to the state with the higher score. **c**, Terminal State Identification (TSI) scores for DRVI and benchmarked methods. This metric quantifies the overlap between identified stationary states and ground-truth terminal states as the number of macrostates increases. **d**, Jaccard index between the true known terminal states and the automatically identified terminal states using CellRank based on the embeddings from different benchmarked methods. **e-h**, Relevant factors for beta (e), delta (f), alpha (g), and epsilon (h) differentiation. For each of these terminal states, a group of relevant factors is provided. For each factor, the top relevant genes in the identified gene program are illustrated as a barplot, and the activity of a relevant gene on the UMAP is plotted. In addition, for consecutive developmental stages, a scatterplot illustrates the differentiation process. The arrows are for visualization and are not derived from the model. SMI, Scaled Mutual Information.

**Figure 5:**
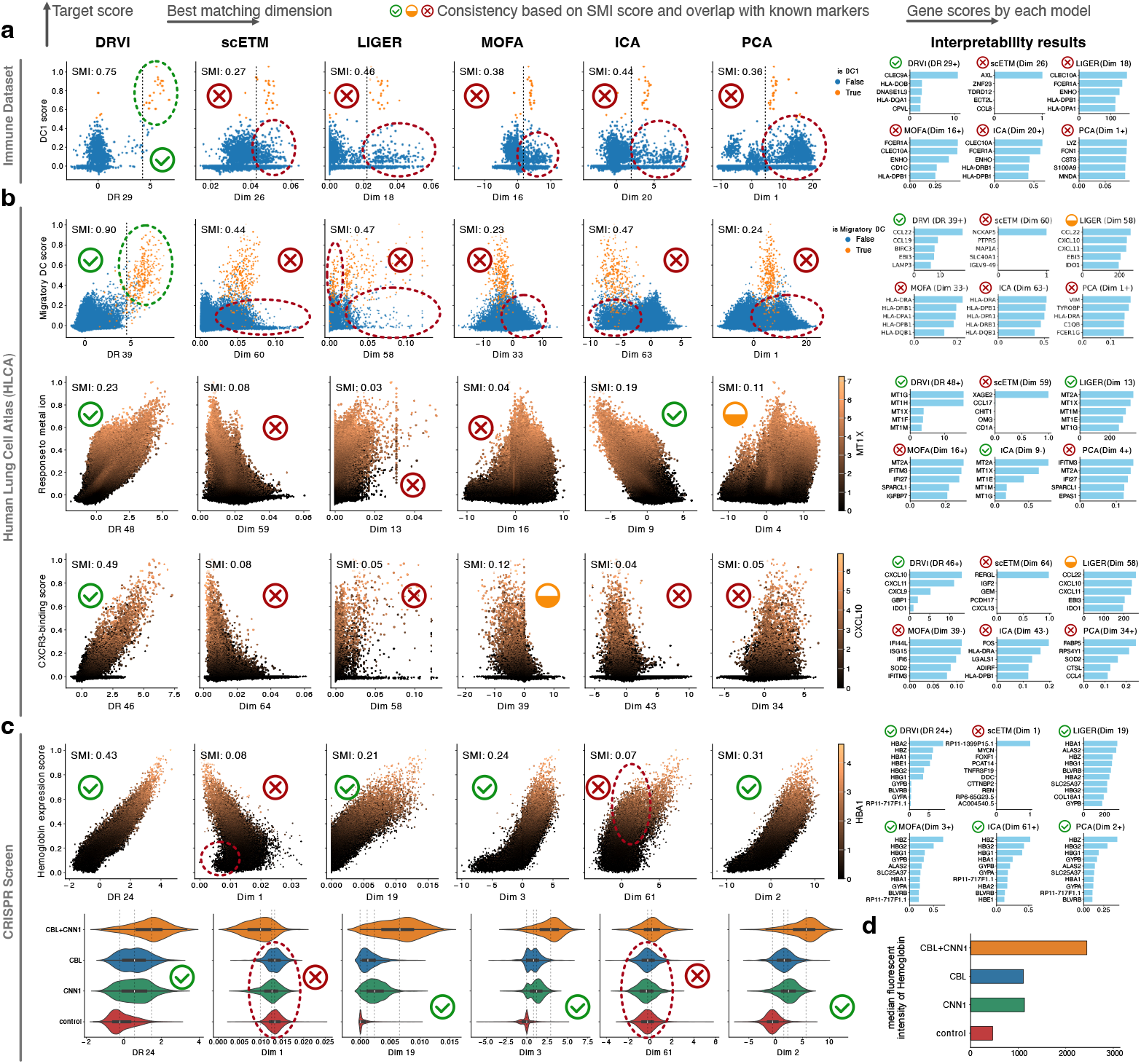
DRVI isolates biological programs that factor-based baselines mix. **a-c**, This comparison demonstrates examples where DRVI outperforms factor-based baselines in identifying biological programs. Green checks, red crosses, and partial marks indicate whether the selected factor separates the program of interest and yields an interpretable gene ranking consistent with known biology. Dashed green circles highlight the successfully disentangled target group, and dashed red circles highlight cell populations that remain mixed with the target group in the best-matching factor. **a**, Immune dataset example. DC1 cells were scored using marker genes, and cells with the highest scores were highlighted. DRVI isolates the DC1 program in a single factor, whereas the highest-scoring factors from scETM, ICA, MOFA, LIGER, and PCA mix DC1 cells with other cells, mostly DC2 cells. **b**, HLCA examples. Migratory dendritic cells, metal-ion response, and CXCR3-binding programs were scored, and the highest-scoring factor was selected for each method. DRVI separates each program into an interpretable factor, whereas baseline factors often show weaker alignment or mixtures with unrelated cells. **c**, CRISPR perturbation example. Hemoglobin activity was scored using hemoglobin genes in the CRISPR screen dataset. DRVI, MOFA, LIGER, and PCA identify a factor whose activity separates the CBL+CNN1 perturbation from CBL and CNN1, and separates both single perturbations from control cells, matching the expected hemoglobin response; ICA and scETM fail to distinguish the relevant perturbation groups (dashed red circles). **d**, Flow-cytometry measurements from the original CRISPR screen study [5] confirm increased hemoglobin expression under CBL+CNN1 perturbation, providing independent validation of the DRVI factor in panel c. Data extracted from [5].SMI, Scaled Mutual Information; DC, dendritic cells.

The cell cycle is a complex, multidimensional process encompassing G1, S, G2, and M phases, frequently modeled as a set of single-dimensional processes corresponding to each phase [63]. We therefore used the G2M and S cell-cycle scores as proxy biological processes and quantified each method’s ability to disentangle them. For each method, we calculated the scaled mutual information (SMI; see ‘Metrics’ in Methods) between each factor and the processes of interest. DRVI, MOFA, LIGER, and ICA were the only models that separated at least one of the S and G2M phases (Supplementary Fig. 8b-e). DRVI achieved a 19% higher sum of SMI scores than the next-best model and exhibited a smoother representation of cycling cells with clearer state boundaries (Fig. 4b and Supplementary Fig. 8f).

In addition to the cell cycle in the developmental pancreas dataset, cells undergo development, namely endocrinogenesis, ultimately reaching four terminal states: alpha, beta, delta, and epsilon. We used scVelo [64] and CellRank [65] to infer velocity fields, macrostates, cell fates, and terminal states. scVelo determines possible future cell states by considering (I) spliced and unspliced counts and (II) the latent space where cell similarities are calculated. Therefore, we supplied embeddings from DRVI and other benchmarked methods as input latent spaces. DRVI improved the quality of velocity vector fields (Supplementary Fig. 9a), reflected by the Terminal State Identification (TSI) score, which measures the overlap between identified macrostates and ground-truth terminal states as the number of macrostates increases. Moreover, we used cellRank to distinguish between initial, transitioning, and terminal macrostates. Only DRVI and LIGER achieved a full TSI score (Fig. 4c and Supplementary Fig. 9b-d). Moreover, only the DRVI embedding enabled CellRank to automatically identify all four terminal states and separate them from transitioning and initial states (Fig. 4d and Supplementary Fig. 9e). Within its interpretable disentangled latent space, DRVI identifies a set of factors leading to these four terminal states, which we selected and describe below.

DR 23+, DR 1+, and DR 11-describe beta-lineage formation, spanning Fev+ beta cells to insulin-producing beta cells (Fig. 4e). For DR 23+, top genes include Mnx1 and Pdx1, which promote endocrine progenitor differentiation into beta cells [66, 67]. Pdx1 also appears among the top genes for DR 1+, consistent with its role in maintaining the beta-cell state [68]. This factor further includes Sytl4, which regulates beta-cell maturation [69], and beta-specific markers reported by Klein et al., including Sntg1, Mapt, Tenm3, and Plppr1 [70]. Finally, DR 11-, marked by Ins1 and Ins2, indicates mature insulin-secreting beta cells. The corresponding scatterplot highlights the latent geometry from endocrine progenitors toward mature beta cells (Fig. 4e).

DR 9+ and DR 26-model development toward delta cells (Fig. 4f). DR 9+ is mainly identified by Hhex, which is required for delta-cell differentiation [71], whereas DR 26-is mainly identified by Sst, secreted by delta cells.

DR 8+ primarily captures alpha and pre-alpha cells, together with subsets of epsilon and pre-beta cells. Although Pou6f2 and Pde1a are computationally suggested alpha-cell markers [70], the remaining top genes are not alpha-specific, suggesting that this lineage is only partially disentangled, potentially due to limited latent dimensionality, insufficient signal, or modeling limitations. By contrast, DR 4+ is mainly identified by Gcg, indicating alpha cells (Fig. 4g).

The lineage from endocrine progenitors toward epsilon cells is captured by DR 23+ and DR 27-. Among the top genes of DR 23+, Cdkn1a, Cacna2d1, Rgs17, Mboat4, Fam107b, and Tmem184c were reported as computational drivers of epsilon progenitors [70]. For DR 27-, the presence of Ghrl indicates the epsilon cell type, and Ascl1, Maged2, Ghrl, Mboat4, Arg1, and Lrrtm3 were computationally derived as drivers of the epsilon population [70]. The scatterplot of DR 23 and DR 27 illustrates differentiation of endocrine progenitors toward epsilon cells (Fig. 4h).

In conclusion, DRVI captures the developmental stages leading to terminal pancreatic endocrine states. Its interpretability identifies relevant genes for each stage, and, without using additional time points, DRVI recovers developmental drivers that align with those previously identified using optimal transport on additional time points [70]. A quantitative evaluation based on CellRank and scVelo further showed that DRVI improved terminal state identification in this dataset.

### Disentanglement allows for the analysis of interventions in latent space

To demonstrate DRVI’s capabilities on perturbational data, we applied it to a CRISPR screen dataset consisting of 104,339 K562 cells perturbed by 106 single-gene and 131 combinatorial perturbations [5]. Program-level latent variables enable visualization, analysis, and differential testing of perturbations on interpretable latent factors rather than in the original high-dimensional, noisy gene-expression space.

To examine whether the latent space preserves perturbation geometry, we focused on IRF1 and SET perturbations and selected, for each method, the two factors with the highest mutual information with these perturbations. Only DRVI and DRVI-AP, and to some extent ICA, scVI-ICA, and LIGER, identified the perturbations in distinct latent factors and displayed additive structure in their latent spaces (Supplementary Fig. 10). DRVI outperformed ICA by 12% and 2% in SMI scores for IRF1 and SET, respectively. Despite SET being one of the perturbations with the highest factor relevance in scVI, this method fails to isolate these variations.

DRVI’s nonlinear factorization also models combinatorial perturbations. We illustrate this with three examples covering synergistic, epistatic, and mutual inhibitory effects. The CBL+CNN1 perturbation significantly increases DR 24+ (hemoglobin expression, Supplementary Fig. 11a) and DR 5+ (SLC25A37 iron importer, which facilitates heme synthesis, Supplementary Fig. 11b). This suggests hemoglobin overexpression, which was confirmed by flow cytometry [5]. In another example, the effect of ETS2 (DR 15-) is canceled when combined with DUSP9 (Supplementary Fig. 11c). Finally, the mutual inhibition between DUSP9 and MAPK1 can be inferred from the decreased response signatures of both genes (DR 15- and DR 8-) in the combined perturbation (Supplementary Fig. 11d).

Perturbation signatures can also serve as external reference variables to assess whether models isolate perturbation effects in individual latent dimensions. In the next section, we use the hemoglobin response as one example for comparing interpretable baseline models. In the quantitative benchmark, perturbation signatures are then used as proxy variables, showing that DRVI better isolates perturbation effects in individual latent factors without using guide RNA information as supervision.

### Interpretable baselines do not consistently isolate known biological programs

The examples above show that DRVI factors can be assigned direct biological interpretations. However, there are other methods that also present interpretable factors, raising the question of whether they can recover the same programs in practice. To test this, we selected programs that DRVI successfully disentangled in the preceding sections and asked whether interpretable baselines isolate them equally well. We compared DRVI, scETM, ICA, MOFA, LIGER, and PCA on a set of known cell identities, pathway activities, and perturbation response programs across three datasets (Fig. 5). For each program, the y-axis shows a ground-truth score derived from known markers, and the x-axis shows the best-matching latent dimension from each model.

In the immune dataset, we focused on DC1 cells. We scored cells using curated marker genes (CLEC9A, CADM1, BATF3, and THBD) and highlighted the cells with the highest scores. DRVI isolates this DC1-associated signal in a single interpretable latent factor whose activity aligns with the marker-positive cells and top relevant genes are consistent with dendritic-cell biology. In contrast, the strongest baseline factors either fail to concentrate DC1 cells or mix them with CLEC10A+ DC2 populations, indicating a failure to disentangle this rare subtype (Fig. 5a).

We next considered three HLCA programs that represent distinct types of biological variation. For migratory dendritic cells, we used HLCA annotations to mark the relevant cells and known markers (EBI3, CCR7, CCL19, LAMP3, and FSCN1) to score them. For metal-ion response and CXCR3-associated chemokine signaling, we scored cells using all genes participating in the corresponding processes: the metallothionein gene family for metal-ion response and CXCL9, CXCL10, and CXCL11 for CXCR3 signaling. Across these examples, DRVI produces factors whose values, SMI scores, and top relevant genes are concordant with the program of interest. For metal-ion response, only DRVI, ICA, PCA, and to some extent LIGER were able to disentangle the process. However, only DRVI successfully disentangles migratory dendritic cells and CXCR3 signaling. In the baseline embeddings, the relevant populations or response programs remain entangled with other variation (Fig. 5b).

Finally, we examined the CRISPR screen dataset, where the CBL, CNN1, and CBL+CNN1 perturbations were previously assayed by flow cytometry for hemoglobin expression (Fig. 5d) [5]. We scored hemoglobin activity using hemoglobin genes and compared each method’s best-matching factor. This program is highly variable and therefore easy to detect, as reflected by its recovery in PC2. We therefore used it to test whether each method can recover the expected perturbation ordering without mixing it with unrelated variations. DRVI separates the CBL+CNN1 perturbation from CBL and CNN1, and separates both single perturbations from control cells, matching the flow-cytometry result that the combined perturbation induces the strongest hemoglobin expression. MOFA, LIGER, and PCA recover this structure to some extent, whereas ICA and scETM fail to distinguish control cells from the single perturbations (Fig. 5c).

### DRVI outperforms previous disentanglement approaches while maintaining integration quality

Disentangled representation learning aims to place distinct sources of biological variation along separate, monosemantic axes of the latent space. For evaluation, we therefore ask whether each source of variation is identified by a distinct latent factor. This is quantified by finding the best one-to-one matching of biological signals and latent factors, known in the disentanglement literature as the latent matching score (LMS). Perfect disentanglement evaluation is fundamentally impossible in biological data because the true set of latent biological processes is unknown. We therefore evaluate disentanglement using available proxy variables, such as annotated cell types, applied perturbations, and known processes such as cell cycle (Fig. 6a). Although this supervised information is used for evaluation, none of the benchmarked models depends on or has access to it during training. By default, we measure similarity of biological signals and latent factors using Scaled Mutual Information (SMI; see ‘Metrics’ in Methods).

**Figure 6:**
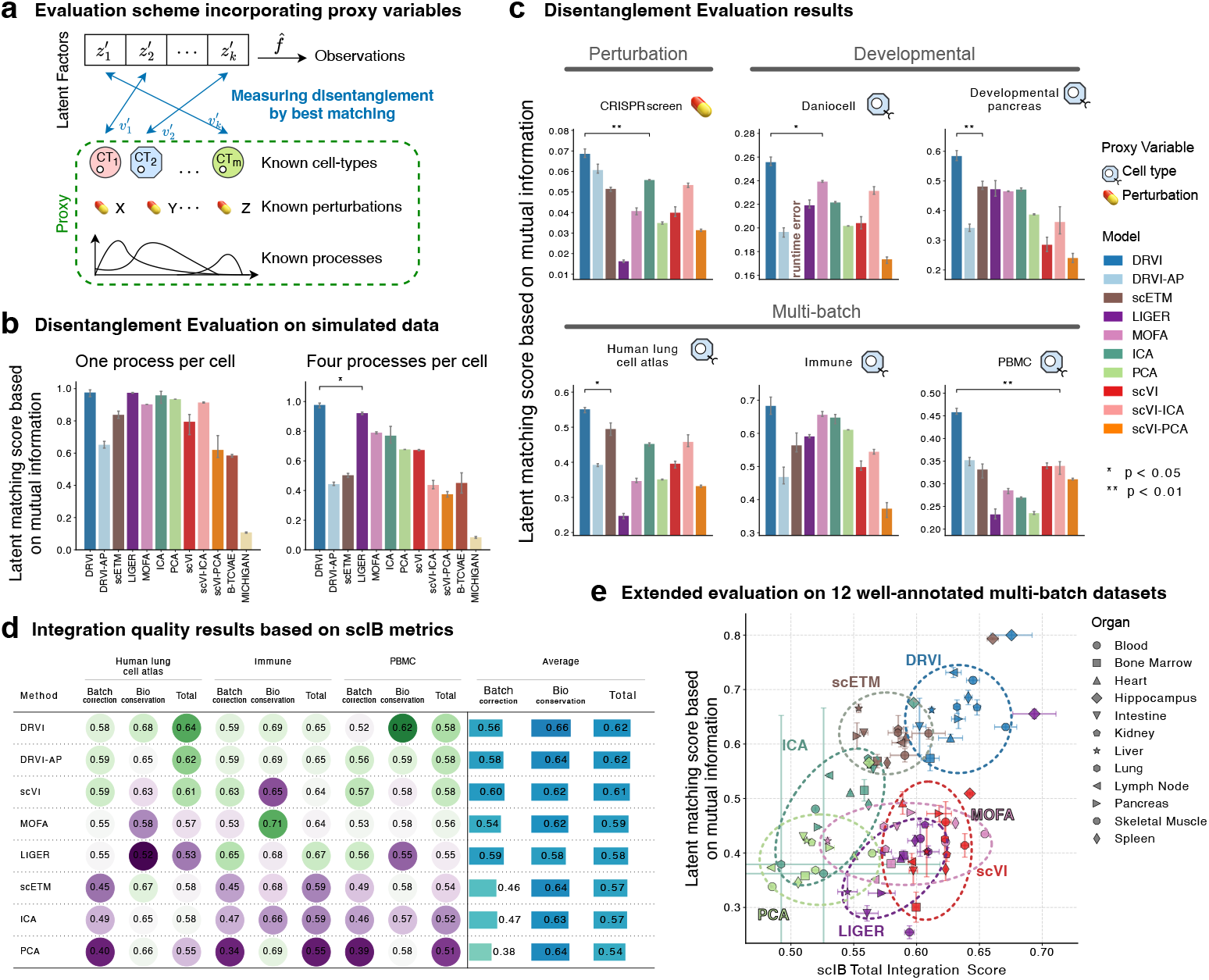
Benchmarking DRVI in terms of disentanglement and integration quality. **a**, Evaluation of disentanglement requires access to ground-truth biological processes, which are often unreachable. We use cell-type annotations, perturbations, and known processes as proxies when available. **b**, Disentanglement evaluation on simulated datasets with known ground truth. Latent matching scores are calculated by finding the best one-to-one alignment of latent factors and underlying processes of the simulated datasets. Disentanglement scores based on mutual information are shown for settings with one active process per cell and up to four active processes per cell. **c**, Disentanglement evaluation on the six primary benchmark datasets, covering perturbation, developmental, and multi-batch settings. Latent matching scores are calculated by finding the best one-to-one alignment of latent factors and proxy variables using mutual information. Asterisks indicate significance levels for the comparison with the next-best method. **d**, Integration quality evaluated using scIB metrics on the three primary multi-batch datasets. Batch correction, biological conservation, and total integration scores are shown for each dataset and averaged across datasets. **e**, Joint evaluation of disentanglement and integration quality on twelve well-annotated CellHint multi-batch datasets from distinct organs. Each point summarizes three training runs per method and organ dataset, with LMS-SMI and total scIB integration score on the two axes. Shaded regions show 95% confidence intervals.

To systematically assess disentangled representation learning, we benchmarked DRVI against unsupervised baselines spanning linear models (PCA, ICA, MOFA, and LIGER), topic models (scETM), nonlinear variational models (scVI, *β*-TCVAE, and MICHIGAN), and post-hoc rotations of the scVI latent space (scVI-PCA and scVI-ICA). We also included DRVI-AP, which replaces LSE aggregation with average pooling, to emphasize the contribution of LSE aggregation.

We first validated the metrics under controlled ground truth using two simulated datasets with known active processes (Fig. 6b and Supplementary Fig. 12). When processes were mutually exclusive across cells, DRVI and linear factor models recovered the ground-truth structure. When cells could contain up to four overlapping processes, only DRVI and LIGER accurately recovered the active programs (Fig. 6b and Supplementary Fig. 13).

We then evaluated the models on six primary benchmark datasets: two atlas-scale multi-batch datasets (HLCA [7] and PBMC [72]), a smaller multi-batch immune dataset [26], two developmental datasets at different scales (developmental pancreas [4] and Daniocell [73]), and one genetic perturbation dataset (CRISPR screen [5]), summarized in Supplementary Table 2. We further evaluated the same methods on twelve multi-batch datasets from distinct organs, each with curated and harmonized annotations from the CellHint project [61]. We used finest-level cell-type annotations as proxy variables for all datasets except the perturbation screen, for which we used perturbation signatures (see ‘Datasets’ in Methods).

Across the six primary datasets, DRVI achieved the highest LMS-SMI in each setting, with statistically significant gains over the next-best method in most comparisons (Fig. 6c). The next-best method varied across settings, and no baseline consistently matched DRVI. The average LMS-SMI across the six primary datasets was 0.43 for DRVI, 0.38 for scETM, and 0.35 for ICA, the next-best models. The significant gaps to scVI and DRVI-AP highlight the essential contributions of the structured decoder and LSE aggregation, respectively.

To further evaluate the benchmarked methods, we explored the same-process neighbors (SPN) similarity function (see ‘Metrics’ in Methods). Consistent with the mutual information metrics, DRVI demonstrated superior performance, improving on average by 11% in LMS-SPN compared to LIGER, the closest competitor (Supplementary Figs. 14 and 15). We also present two additional disentanglement metric families (Supplementary Fig. 16): Most Similar Averaging Score (MSAS) and Most Similar Gap Score (MSGS). MSAS measures coverage by calculating the maximum similarity between each proxy variable and latent factors, while MSGS additionally penalizes similarity to other factors (see ‘Metrics’ in Methods). When using SMI as the similarity function, MSGS reduces to the Mutual Information Gap (MIG). DRVI consistently achieved the highest average MSAS metrics, whereas ICA, scVI-ICA, DRVI, and LIGER each performed best for specific datasets or sub-metrics of the MSGS family. We find that the decline in DRVI’s MSGS performance on the Immune dataset stems from a limitation in the ground-truth annotations rather than the model. DRVI identifies finer, more biologically precise cell subtypes (e.g., Naive vs. Th2 cells) that are grouped under single ground-truth labels. The MSGS metric incorrectly penalizes this identification of more granular subtypes (Supplementary Fig. 17). Considering this, the LMS metric family is favored due to its alignment with the formal definition of disentanglement, less sensitivity to imperfect proxy variables, and its unique ability to penalize overlapping processes within a single factor, a key obstacle to interpretability.

Some unsupervised methods, such as ICA and MOFA, enforce disentanglement through strongly limiting model constraints like linearity and sparsity, restricting their ability to represent rich, multi-batch data. To assess the applicability of disentangled models on complex data, we assess integration quality alongside disentanglement. For datasets containing multiple batches, we assessed biological signal conservation and batch correction using scIB metrics [26]. Across the three primary multi-batch datasets, DRVI achieved the highest biological conservation score and the highest total integration score, while scVI-based embeddings achieved stronger batch correction (Fig. 6d).

The expanded benchmark across twelve organ datasets with curated annotations from CellHint showed the same pattern. In this broader multi-batch setting, DRVI consistently combined high cell-type disentanglement with strong integration, successfully bridging the gap between linear disentangled models and flexible generative methods (Fig. 6e). Details of per-organ results are provided in Supplementary Figs. 18, 19, and 20.

### DRVI extends to further data modalities

We primarily demonstrate the utility of DRVI in single-cell transcriptomics data. However, DRVI easily extends to other modalities, such as single-cell DNA accessibility (scATAC), to obtain interpretable latent factors. For this purpose, DRVI supports Negative Binomial, Poisson, and Gaussian (with MSE loss) noise models. As an example, we applied DRVI to the single-cell ATAC modality of the NeurIPS 2021 multimodal data integration challenge [74]. Specifically, we employed the Poisson noise model on ATAC fragments, which has recently been shown to be the proper assumption for this modality [17]. To assess disentanglement and integration, we compared DRVI against DRVI-AP, PoissonVI [17] (a recent best-performing method), and PeakVI [75] (a widely used method in scATAC-seq integration). As in scRNA-seq, we observe that DRVI combines disentanglement with robust integration performance: DRVI improves disentanglement with a 97% improvement in LMS-SMI and a 167% improvement in LMS-SPN over the next best-performing model (Supplementary Fig. 21c). Additionally, DRVI achieves comparable total scIB integration quality to PoissonVI and PeakVI, demonstrating its applicability to the ATAC modality (Supplementary Fig. 21d).

## Discussion

We introduced DRVI, an unsupervised deep generative model that learns interpretable nonlinear factors for single-cell transcriptomics by replacing the usual fully connected decoder with additive decoder subnetworks and LSE pooling. While single-cell generative models have become central to integration and representation learning, their latent spaces are typically used for visualization and clustering and are difficult to interpret mechanistically. This limits their use for discovery tasks in which cell states, pathways, technical responses, and perturbation effects can overlap within the same cell. Linear factor models provide interpretable axes, but their linearity limits flexibility in complex multi-batch data. By providing biologically navigable latent spaces, DRVI bridges these two model classes and combines the scalability and integration quality of nonlinear generative models with the factor-level interpretability of linear approaches. Unlike the log-space additivity of linear factor models, which implies multiplicative effects between overlapping processes in count space, DRVI’s LSE pooling more naturally represents overlapping programs as additive count-space contributions.

The architectural contribution of DRVI is not restricted to the specific model implementation studied here. Additive decoder subnetworks and LSE pooling define a decoder inductive bias that can, in principle, be adopted by other generative models regardless of their noise model, prior distribution, approximate posterior, or training paradigm. This modularity may allow future single-cell models to incorporate the DRVI decoder while retaining domain-specific likelihoods or covariate structures, thereby gaining factor-level interpretability.

Across atlas-scale datasets, developmental data, and perturbation screens, DRVI outperformed benchmarked methods in recovering interpretable latent factors while maintaining competitive integration quality. In the immune dataset, HLCA, developmental pancreas, and CRISPR screen, it isolated rare cell states, pathway programs, and combinatorial responses that factor-based baselines failed to separate. These programs are consistent across related datasets and transferable to new cohorts, supporting comparison and reuse across experiments.

Despite promising results, our work, as a step towards unsupervised disentangled representations in single-cell omics, has limitations. While highly effective for current individual atlases, DRVI is more resource-intensive than entangled autoencoder frameworks such as scVI or DCA, and scaling the model to hundreds of millions of cells, all genes, and thousands of latent variables remains a challenge for future optimization. As with PCA, ICA, and MOFA, DRVI may encode two distinct concepts along opposite directions of a single axis, requiring both directions to be inspected; this can be mitigated by applying an activation function during training or by clipping factors post hoc. The LMS metric already penalizes this behavior through its one-to-one matching, yet DRVI still achieves the best results. Perfect disentanglement evaluation is also fundamentally impossible in biological data because the true set of latent biological processes is unknown. We therefore evaluate models using imperfect proxies such as cell-type annotations and perturbation labels. These proxies are often incomplete, coarse, or combinatorial, which limits metric precision, yet they remain the most principled evaluation strategy available.

Future work can extend DRVI’s inductive bias to multiomic, velocity-aware, and spatial generative models. Its count-space additivity is especially relevant for settings with compositional measurements, imperfect segmentation, background contamination, doublets, and combinatorial perturbations. In spot- and segment-based assays, counts reflect compositional mixtures rather than single cells, so count-space additivity may support deconvolution and robustness to imperfect segmentation, although extensions that incorporate spatial coordinates remain necessary. DRVI’s ability to generate sparse latent variables suggests its potential for scaling to larger, more complex datasets. A future direction may be to expand the model to multiple atlases and create a comprehensive, unified dictionary across organs.

## Data and code availability

All datasets analyzed in this manuscript are public and have been published in other papers. We have referenced them in the manuscript and provided the pre-processing steps at http://github.com/theislab/drvi_reproducibility.

DRVI is available as a package at http://github.com/theislab/drvi. The code to reproduce the results and figures is available at http://github.com/theislab/drvi_reproducibility.

## Author contributions

A.A.M. and F.J.T. conceived the project. A.A.M. designed and implemented the algorithm. A.A.M. performed experiments and analyses. A.A.M. and F.J.T. wrote the manuscript. F.J.T. supervised the work.

## Acknowledgments

This work was supported by the Deutsche Forschungsgemeinschaft (DFG) through the Leibniz Prize and project 458958943 (grant number 5010338). A.A.M. is a member of the ELLIS PhD Program of the European Laboratory for Learning and Intelligent Systems (ELLIS) Society. F.J.T. acknowledges support from the Helmholtz Association’s Initiative and Networking Fund through Helmholtz AI (grant number ZT-I-PF-5-01) and from the Leibniz Prize from DFG.

We thank Malte D. Luecken, Lisa Sikkema, Sara Jimenes, Dominik Klein, Leander Dony, Alessandro Palma, Mariia Minaeva, Mojtaba Bahrami, Anastasia Litinetskaya, Mohammad Lotfollahi, Philipp Weiler, Daniel Strobl, and Michaela Müller for valuable discussions and feedback on the manuscript and the framework. We also thank all members of the Theislab for their support.

## Declaration of generative AI and AI-assisted technologies in the writing process

During the preparation of this work, the authors used ChatGPT by OpenAI, Gemini by Google, and Grammarly by Grammarly Inc. to improve the readability and language of the text. After using these tools or services, the authors reviewed and edited the content as needed and take full responsibility for the content of the publication.

## Methods

### DRVI: Disentangled Representation Variational Inference

A representation is disentangled when distinct sources of variation occupy separate latent dimensions, so each factor reflects one coherent process rather than a mixture of several [27]. Formally, a learned decoder is disentangled with respect to a ground-truth set of factors when a one-to-one correspondence (permutation of factors) exists between its latent factors and the ground-truth factors, up to per-factor transformations [27, 34] (see Supplementary Note 4 for formal mathematical definitions). In single-cell omics this includes (but is not limited to) recovery of cell types, signaling pathways, inflammation processes, and cell cycle stages as latent factors. While some disentangled representation learning frameworks assume statistical independence of underlying factors, aligned with recent advancements [27, 34] and the causal behavior of biological processes, we do not make such an assumption (Supplementary Note 5 and Supplementary Fig. 22).

DRVI is an unsupervised deep generative model that employs log-sum-exp (LSE) pooling on top of additive decoders to disentangle latent factors. The additive decoder architecture splits the latent space into individual slots, decodes them into the output space, and sums them up [34]. This architecture has been shown to provide provable disentanglement under specific assumptions. However, we practically observe that, without proper modifications, combining current variational autoen-coders in single-cell omics with additive decoders does not result in disentangled representations (DRVI-AP baseline). Thus, we replace the sum aggregation in additive decoders with LSE pooling, which results in higher-order interactions within slots and first-order interactions across slots. The interaction asymmetry framework explains why this modification works: disentanglement can be achieved when each slot has higher order interactions within itself and lower order interactions across slots [76].

To increase the usability and highlight the utility of our architecture, we built our model based on the scvi-tools framework and minimized the deviation from the scVI model [12, 15, 77].

In the following sections, we present the generative model of DRVI, explain its decoder architecture and pooling function, and discuss additional details, including interpretability and the evaluation framework.

### The Generative Model

We use capital letters for random variables (e.g., *X*), lowercase for realizations (e.g., *x*), and upper-case bold for observation matrices (e.g., ***X***, with IID rows ***X***_1_, …, ***X***_*n*_). Capital letters may also denote constants (e.g., *N*).

A generative process is defined by a latent space Ƶ ⊂ ℝ^*K*^, a distribution *p*_*z*_(*z*) over Ƶ, and a decoder *f*: Ƶ → ℝ^*M*^ mapping latent variables to the observational space.

A(pooling function defined by *ψ, σ*: ℝ → ℝ aggregates a set of real-valued numbers *y*_1_, …, *y*_*T*_ by *σ*(∑_1≤*t*≤*T*_ *ψ*(*y*_*t*_)). Pooling functions can also be applied element-wise to a set of vectors or tensors.

Assume that we have an observed count matrix ***X*** ∈ ℕ^*N*×*M*^, where *N* is the number of cells and *M* is the number of genes. Optionally, we can consider a covariate vector ***C*** ∈ ℝ^*N*×*R*^, where *R* is the dimensionality of the batch covariate for each cell. For example, in batch correction frameworks, ***C***_*i*_ could be a one-hot vector representing the batch identity if modeled as in scVI [12], or an embedding vector if modeled as in scPoli [38] (DRVI accepts both conventions). Then, the goal is to discover latent representation ***Z*** ∈ ℝ^*N*×*K*^ where *K* is the latent space dimension for the observed data. The generative process is defined by the pooling of multiple decoders that map latent space blocks to parameters of the noise distribution. Specifically:

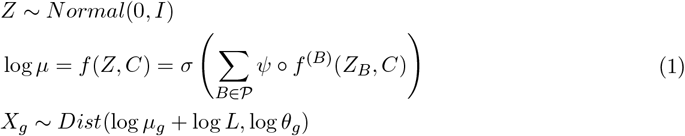

Here, *f* is 𝒫 the decoder, 𝒫 is a partition of the latent space dimensions, *f* ^(*B*)^ denotes the additive decoder subunits defined per block (referred to as decoder subnetworks), *σ* and *ψ* form a pooling function, *Dist* models the noise distribution, *θ*_*g*_ is a parameter indicating gene-wise dispersion or variance, *L* corresponds to the random variable indicating library size of the cell, and *Z*_*k*_ denotes the *k*-th dimension of *Z*. DRVI users have an option to alternatively define *θ* based on batches or optimize it like *µ* using variational inference. We set *L* to the observed values of the library size rather than incorporating it into the generative model, which is aligned with standard practices in the field.

DRVI supports Normal, Poisson, and Negative Binomial noise models. Overall, *Dist*(log *µ*, log *θ*) is defined as:

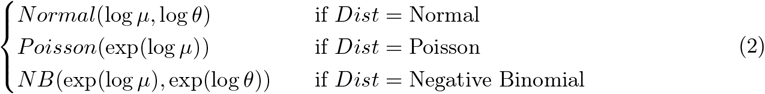

We parametrize the distributions in log space to ensure a clear formulation within our framework and to emphasize that all other methods also calculate their parameters in log space and transform them back to count space in the last step. For the Normal noise model, input data should be log-transformed (e.g., log(1 + *x*), optionally with library-size normalization). The Poisson noise model discards the dispersion parameter.

Fitting the generative process given observed data involves maximizing the log-likelihood function log ℙ_*θ*_(***X***|***C***) =∑_*n*_ log ℙ_*θ*_(***X***_*n*_|***C***_*n*_) where ***X*** is the matrix of observations, ***C*** indicates the matrix of batch covariates, and *θ* represents all parameters in the model. Since the marginal log-likelihood log ℙ_*θ*_(***X***|***C***) is intractable, we optimize the evidence lower bound (ELBO) via variational inference [12, 78]. An encoder network parameterizes an approximate posterior *q*_*ϕ*_(*Z*|*X, C*) as a diagonal Gaussian, and the decoder specifies ℙ_*θ*_(*X Z, C*); together they form a variational autoencoder optimized with stochastic gradient descent and the reparameterization trick [78, 79]. We use the Adam optimizer (learning rate 10^−3^) with KL annealing that linearly increases the KL weight from 0 to 1 during training. Early stopping is not used for DRVI or scVI to allow learning a more structured latent space.

### Additive Decoders, Pooling, and Disentanglement

The additive decoder and interaction asymmetry frameworks establish theoretical conditions under which disentanglement is guaranteed [34, 76]. These conditions are typically proven on very small toy examples [34] or heuristically optimized [76]. Similarly, we show that DRVI with LSE pooling heuristically satisfies these conditions by encouraging higher-order interactions within splits and first-order interactions across splits.

#### LSE Pooling

With *σ* = log and *ψ* = exp:

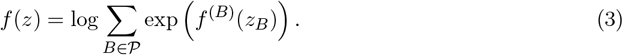

The *σ* = log term can be theoretically disregarded through a suitable transformation of the outputs (Remark 1 of additive decoders [34]). Among the conditions that can guarantee disentanglement, interaction asymmetry requires within-split interactions to be of higher order than cross-split interactions [76]. Considering *ψ f* ^(*B*)^ = exp *f* ^(*B*)^ as decoder functions and ignoring *σ*, our generative process can be considered as an additive decoder where a shared activation function (*ψ* = exp) is used at the end of each decoder subnetwork. The shared activation creates second-order interactions within each split, while the additive architecture keeps cross-split interactions to first order (detailed in Supplementary Note 7). Clearly, *ψ* = exp is effective and cannot be ignored.

#### Average Pooling

With *σ*(*x*) = *x/K* and *ψ* is the identity function, where *K* = |𝒫|:

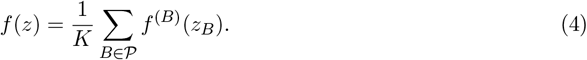

With the shallow MLPs typical in single-cell representation learning, this produces insufficient interactions within splits. In the extreme case of linear subnetworks, the decoder reduces to an affine transformation, making disentanglement impossible due to the well-known unidentifiability of linear ICA under independent Gaussian factors. Empirically, average-pooled additive decoders (DRVI-AP) improve only marginally over non-additive baselines (see Results).

### Mechanistic interpretation

LSE pooling provides a suitable mechanistic interpretation of the generative process. While the additivity of factors in log-transformed space of linear factor models implies multiplication in count-space (Supplementary Note 2), in LSE pooling, factor effects add up in gene count space. This aligns better with the inherent additivity of transcribed genes and is consistent with additive technical effects such as doublets and background expression commonly observed in single-cell omics (Supplementary Note 3 and Supplementary Fig. 23).

The decoder models mean parameters in log space. With average pooling (DRVI-AP, similar to factor models), decoder outputs sum in log space, which corresponds to multiplication in count space. Formally, let log 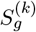 denote the output of the *k*th decoder for gene *g*:

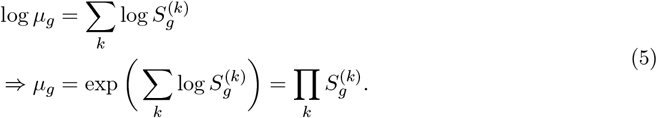

The same multiplicative behavior arises in matrix factorization and, to some extent, standard autoencoder models (see Supplementary Note 2 for details). This is unproblematic when factors affect disjoint gene sets, but biologically implausible when a gene is regulated by multiple factors.

With LSE pooling (DRVI), decoder outputs aggregate via the log-sum-exp function, which corresponds to addition in count space:

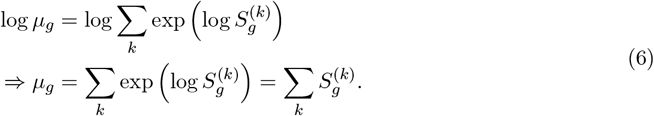

LSE pooling therefore implicitly encodes the assumption that “distinct but overlapping biological processes combine additively in gene count space,” which is more plausible than the assumption of linear factor models and average pooling that “distinct but overlapping biological processes multiply in gene count space.” (Supplementary Fig. 23 and Supplementary Note 3).

### Modeling Details

The encoder is an MLP that accepts log-transformed count data and outputs mean and variance vectors of a diagonal Gaussian posterior. Each MLP layer consists of an affine transformation, ELU activation, and stateless layer normalization, with dropout applied to inputs and hidden layers. Decoder subnetworks *f* ^(*k*)^ use the same architecture.

Decoder subnetwork parameters can be shared at different levels (none, last layer, or all layers); sharing all parameters is the default and works well in practice. Decoder outputs are aggregated via LSE pooling (default) or average pooling to form the output distribution parameters.

For multi-batch datasets, batch covariates are concatenated with each latent split and decoded by the subnetworks *f*_*k*_. DRVI supports one-hot encoding and vector embedding strategies for batch covariates [12, 38]; we use one-hot encoding in all analyses for fair comparison.

The choice of noise model depends on the data modality. The Negative Binomial distribution is standard for scRNA-seq [12, 37, 80, 81], while the Poisson distribution may be more suitable for scATAC-seq [17]. A Normal noise model is also supported for pre-normalized data [13, 60].

To enhance numerical precision, we introduce a log-space parametrization of the Negative Binomial likelihood, computing the likelihood directly from log *µ* instead of *µ* (Supplementary Note 8). The predicted mean is adjusted to match the observed library size after aggregation.

DRVI users can introduce multidimensional splits by adjusting a model parameter. This defaults to dimension-wise splits (one latent dimension per split), the most powerful disentanglement scenario.

### Interpretation and ordering of learned factors

Interpreting latent factors in terms of gene involvement helps identify both known and novel biological programs. In factorization models, loadings directly quantify gene-factor associations. In nonlinear models such as scVI, interpretability is challenging because each factor’s effect depends on all other factors. DRVI’s additive architecture removes cross-factor interactions: each subnetwork *f* ^(*i*)^(*z*_*i*_) decodes its factor independently into count space, making factors individually interpretable. We traverse each latent factor separately to infer top associated genes (Supplementary Note 9).

Unlike PCA, where principal components are ordered by explained variance, autoencoder latent factors are inherently unordered. We exploit the additive architecture to sort factors by their contribution to reconstruction (Supplementary Note 10).

### Hyperparameters

We explored various choices of normalization, activation functions, noise models, parameter sharing, and aggregation functions on the immune dataset. We then selected stateless layer normalization, ELU activations, log-space parametrized Negative Binomial noise (for RNA), full parameter sharing across decoder subnetworks, and LSE pooling. These settings were applied uniformly across all datasets without further tuning. Architecture parameters (latent dimensions, hidden layers, and depth) were matched between DRVI, DRVI-AP, and scVI for consistency. Detailed parameters for each dataset are provided in Supplementary Table 3. We tracked experiments using Weights & Biases [82].

### Metrics

Since DRVI aims to bridge the gap between disentangled linear models and flexible non-disentangled methods, we evaluate its performance from two perspectives: disentanglement and integration.

We assess the integration quality of multi-batch datasets based on the scIB metrics framework. Details on these metrics can be found at [26]. As in the original study, the total score is derived by averaging biological conservation and batch integration scores with weights of 0.6 and 0.4, respectively.

We measure disentanglement by finding the optimal one-to-one matching between underlying biological processes and latent factors that maximizes pairwise similarities. This defines the Latent Matching Score (LMS) [34]:

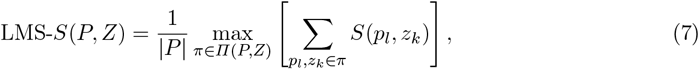

where *S* is a similarity function, *P* the set of underlying biological processes, *Z* the set of latent factors, and *Π* the set of all injective correspondences from *P* to *Z* (permutations when |*P*| = |*Z*|). We define two similarity functions:

#### Scaled Mutual Information (SMI)

We normalize mutual information by the entropy of the biological process so that a maximum of +1 is achievable:

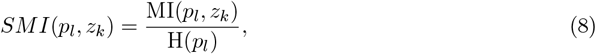

where MI is the mutual information and *H* the entropy. We discretize both vectors into 10 uniform bins before computing MI and H. SMI is also known as the uncertainty coefficient [83].

#### Same-Process Neighbors (SPN)

For binary on/off processes, SPN measures whether cells in the “on” state reside together along a latent factor. It computes the excess probability that two neighboring cells (sorted by *z*_*k*_) are both active, adjusted for random chance:

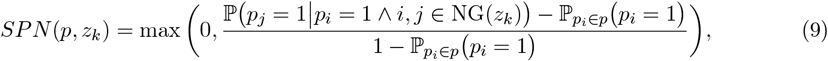

where *NG*(*z*_*k*_) denotes the set of adjacent cell pairs when sorted by factor *k. LMS*-*SPN* can be computed in *O*(|*P*|*Kn* log(*n*)). This can be interpreted as an approximation to the precision of a 1NN classifier predicting the activation of an underlying biological process from a given latent factor.

We use *LMS* − *SMI* as the default metric to evaluate the disentanglement of a trained model with respect to known biological processes. While *SMI* accepts both discrete and continuous processes, *SPN* is designed specifically for on/off processes.

We use two secondary metric families to complement LMS. The Most Similar Averaging Score (MSAS) averages similarity of best matching latent factors for each process, measuring coverage. The Most Similar Gap Score (MSGS) additionally penalizes a process being similar to other latent factors; with SMI, it reduces to the Mutual Information Gap (MIG) [84]. Formally:

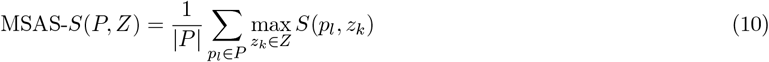

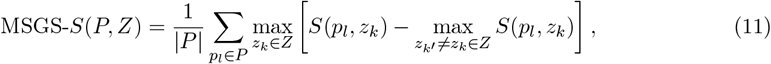

where *S* can be any similarity function, and *P* and *Z* are defined as before.

We prefer LMS because MSAS and MSGS do not penalize multiple irrelevant signals within a factor, the primary obstacle to interpretability (Supplementary Table 4). Moreover, MSGS heavily penalizes models that correctly disentangle finer subtypes when using coarse-grained proxy variables for evaluation (Supplementary Fig. 17), and it assumes statistical independence, which biological processes do not satisfy (e.g., cell cycle is not independent of cell type, as not all cell types proliferate).

Since underlying biological processes are typically unknown, we use cell-type annotations or perturbation labels as proxy variables. These proxies are imperfect: perturbations can be redundant or combinatorial, and coarse cell-type annotations penalize models that correctly resolve subtypes. Consequently, disentanglement score ranges vary across datasets. Despite these limitations, proxy-based evaluation remains the best available approach.

In addition to disentanglement metrics, we used Rank Biased Overlap (RBO) similarity score to demonstrate the consistency of interpreted signatures across datasets. For two sorted sets *S* and *T*, RBO is defined as:

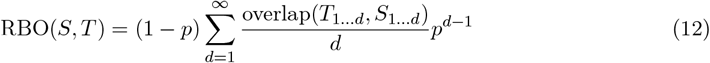

Where *p* ∈ (0, 1) is a parameter controlling the importance of top elements and overlap(*T*_1…*d*_, *S*_1…*d*_) is the number of shared items between the top *d* elements of *S* and *T*. We use *p* = 0.9 by default.

### Baseline models

All baseline models were trained with the same number of latent dimensions as used for DRVI. For non-count baselines, we used the normalized log-transformed gene expression provided by the dataset, or normalized total counts per cell and log-transformed the data using Scanpy [85, 86].

- **PCA** from scikit-learn [87], was applied to the log-transformed and normalized data. Prior to the transformation, the data was centered without scaling.
- For **ICA**, we applied the FastICA [88] from scikit-learn [87]. Since the number of samples exceeded the number of features across all datasets, we chose the whiten_solver=“eigh” option. The data was centered before applying ICA, without any scaling.
- **scVI** was trained with the same epochs and architecture as DRVI, without early stopping. For **scVI-PCA** and **scVI-ICA**, scikit-learn PCA and FastICA were applied to the learned latent space of scVI.
- *β***-TCVAE** and **MICHIGAN** were run via the MICHIGAN package [89]. We selected `GradientPenalty_lambda=100` after grid-searching key scalar hyperparameters over {0.1, 1, 10, 100}.
- **MOFA** was run via mofapy2 [35] on log-normalized data with `convergence_mode=`“`fast`”. We specified batch labels as `groups` to account for batch effects. We enabled `spikeslab_weights` and `spikeslab_factors` for sparse factor loadings after grid-searching sparsity settings.
- **scETM** [90] was trained on raw counts with batch covariates and default hyperparameters; the topic mixture (*θ*) defined the latent space.
- **LIGER** was run via pyLIGER [36] in a batch-aware manner on count matrices; the normalized cell factors (*H*_norm_) defined the latent space.

### Benchmarking details

Quantitative benchmark results were computed from three random seeds. Statistical annotations in benchmark figures compare DRVI to the next-best method per dataset using t-tests. The complete per-seed metric values and summary statistics are provided as Supplementary File 1.

### Consistency and transferability of models

To evaluate factor consistency, we trained each model independently on each reference dataset and compared ranked factor-associated gene signatures across datasets using RBO. For each method, dataset, and cell type, we selected the latent factor with the highest SMI against the binary indicator of that cell type. For signed factors (PCA, ICA, MOFA, DRVI), we used the direction associated with the cell type and compared the corresponding ranked gene signatures. RBO values for same-cell-type comparisons were computed between factors selected for the same annotation across independently trained models, whereas background comparisons used factors selected for different annotations.

To evaluate factor transferability, we trained each model on a reference dataset and projected the remaining datasets into the learned latent space using method-specific interfaces. For DRVI and scETM, query profiles were passed directly through the trained encoders. For PCA and ICA, normalized query expression was projected onto the fitted coordinate bases. For MOFA and LIGER, which do not provide native transform APIs, query cells were projected using feature matrices. For MOFA, query cell loadings were obtained by linear least-squares projection with pseudo-inverse of the reference loading matrix, *Z* = *X*(*W* ^†^)^*T*^. For LIGER, query loadings (*H*) were estimated from expression profiles by optimizing *X* ≈ *HW*^*T*^ with 50 multiplicative-update iterations.

### Simulated data

In addition to real-world datasets, we benchmarked models on two simulated datasets (see ‘Datasets’ and Supplementary Fig. 12). One dataset featured unique cellular processes per cell, and the other included up to four overlapping processes per cell group.

### Runtime Analysis

We benchmarked DRVI against scVI (for scRNA-seq) and peakVI (for scATAC-seq). DRVI’s run-time scales linearly with the number of cells. Despite higher floating-point operations, its parallel implementation maintains runtime on the same order as scVI and peakVI (Supplementary Fig. 24). Post-training steps (interpretability pipeline and factor sorting) complete in under 80 seconds per dataset (Supplementary Fig. 25). DRVI requires the same CPU memory as scVI, as both are based on the scvi-tools framework. Experiments were performed on a server with two Intel Xeon Platinum 8480+ processors and eight NVIDIA H100 (80GB) GPUs, constraining each run to **a single GPU** and **20 CPU threads**.

### Datasets

In this section, we provide information on the datasets used in this work (summarized in Supplementary Table 2). The pre-processing steps for each dataset are described. UMAP embeddings shown in figures were computed with scanpy or rapids-singlecell [85, 91]. Additionally, the code to reproduce simulated data as well as the pre-processing steps are available at https://github.com/theislab/drvi_reproducibility.

### Simulated

We simulated two single-cell gene expression datasets to evaluate representation learning methods under controlled conditions. Each dataset consists of *G* = 1000 genes and *N* = 6000 cells, each assigned to either a specific condition or a control. For each condition *c* ∈ {1, …, *C*}, we sample a binary vector **y**_*c*_ ∈ {0, 1}^*K*^, where *K* = 20 is the number of latent biological processes and *y*_*c,k*_ = 1 indicates that process *k* is active in condition *c*. Each cell under condition *c* inherits the same binary process activity vector. Latent representations **z**_*i*_ ∈ ℝ^*K*^ are drawn independently for each cell *i*, where ∼ *z*_*i,k*_ Beta(10, 2) if *y*_*c,k*_ = 1, and *z*_*i,k*_ = 0 otherwise. Control cells are assigned zero latent activity (**z**_*i*_ = **0**). We ensure that all processes are represented across the dataset and control the degree of overlap across conditions by varying *D* = max ∥**y**_*c*_∥_0_, the maximum number of active processes per condition. In one dataset, each condition is linked to a single active biological process (*D* = 1). In the other, four distinct processes (*D* = 4) may simultaneously be activated per condition, creating controlled overlap and compositional complexity.

Gene expression values are generated additively by mapping each active process to a sparse (and potentially overlapping) subset of genes using nonlinear monotonic cubic spline functions [92]. Specifically, the expression of each relevant target gene is computed individually using a smooth monotonic transformation of the latent factor, simulating process-specific gene regulation. To account for biological variability consistent with the negative binomial characteristics of single-cell data, we model each process-specific contribution with a Gamma distribution, Gamma(*α* = *s, θ* = *µ/s*), where *µ* is the derived process-wise mean and *s* is a shape parameter per gene uniformly sampled from (1, 5). In addition to ground-truth-related variation, we introduce background noise by injecting gene-wise Gamma-distributed noise with shape and scale parameters uniformly sampled from (0.05, 0.1) and (0.1, 10), respectively. The final expression profile for each cell is obtained by summing the noise vector and process contributions, followed by Poisson sampling. The code to generate simulated data is available in the reproducibility repository.

### Immune Dataset

The immune dataset consists of 32,484 cells collected from 4 human PBMC studies, including 9 batches and 16 distinct cell types after pre-processing. The data is obtained from Luecken et al. (2022) [26, 93]. The “Villani” study was excluded due to its non-integer values. The dataset was subset to 2,000 highly-variable genes (HVGs) in a batch-aware manner [26].

### The HLCA

The HLCA consists of a core dataset and an extended version [7]. We obtained the curated core part from cellxgene (https://cellxgene.cziscience.com/collections/6f6d381a-7701-4781-935c-db10d30de293), which includes 584,944 cells across 166 samples from 14 datasets. This dataset contains annotations at different levels. We used samples as the batch covariate and the finest-level annotations comprising 61 cell types to benchmark the methods. The dataset was subset to 1,996 HVGs originally used to construct the HLCA reference model available on Zenodo (https://doi.org/10.5281/zenodo.7599104).

### CRISPR Screen Dataset

The CRISPR screen dataset, also known as the Norman perturb-seq dataset, comprises 104,339 single-cells of K562 cell line (including control cells) perturbed by 106 single-gene perturbations and 131 combinatorial perturbations [5]. The 238 unique perturbation signatures were used to benchmark methods. We downloaded the pre-processed data using the pertpy package [94]. The data was limited to the 5,000 HVGs originally provided in the pre-processed data.

### PBMC Dataset

The PBMC dataset comprises 647,366 cells across 143 samples from 130 healthy and COVID-19 patients [72]. The data was collected at three sites, and we used the site as the batch covariate to account for the highest technical variation. We used the finest-level annotation containing 51 unique cell types for benchmarking the methods. The data was obtained from https://www.covid19cellatlas.org/ and was subset to 2,000 HVGs in a batch-aware manner.

### Developmental Pancreas Dataset

The developmental pancreas dataset consists of 3,696 pancreatic mouse cells during endocrinogenesis at embryonic day 15.5 [4]. The data was obtained from the scVelo package [64]. We used scVelo to pre-process and subset the data to the top 2,000 HVGs. The CellRank package provides a subset of this data with finer annotations [65, 95]. We transferred these finer annotations to the complete dataset and used the finest annotation comprising 15 unique cell states for benchmarking.

### Daniocell Dataset

The Daniocell dataset is a comprehensive transcriptional atlas of early zebrafish development, comprising 489,686 cells from 62 stages during zebrafish embryogenesis [73]. The data covers 20 tissues and 156 identity clusters, which we used to benchmark the data. We obtained the h5ad object from https://zenodo.org/records/8133569 [96]. The pre-processing of this data involved marking data with the annotations provided in the original publication, removing author-identified doublets, and subsetting data to 2,000 HVGs.

### NeurIPS 2021 Dataset

The NeurIPS 2021 Dataset contains nearly 120K single-cell paired RNA-ATAC and RNA-ADT (antibody-derived tags) measurements from the human bone marrow of 10 donors [74]. We used the ATAC modality of this dataset consisting of 69,247 cells from 13 batches covering 22 cell types. The dataset is retrieved from GSE194122 and subset to 14,865 highly variable peaks.

### Extended Organ-Specific Benchmarks

For the extended benchmark, we used twelve multi-batch datasets from blood, bone marrow, heart, hippocampus, intestine, kidney, liver, lung, lymph node, pancreas, skeletal muscle, and spleen, with expert-curated annotations from the CellHint project [61]. The datasets can be downloaded from the organ atlas page of CellTypist (https://www.celltypist.org/organs). For each organ, we used 4,000 HVGs, sample as the batch covariate, and curated annotations as the cell-type proxy variable for disentanglement metrics. For bone marrow, intestine, and skeletal muscle, we trained scVI and DRVI with gene-batch dispersion modeling due to higher batch-dependent dispersion variability.

## Supplemental Material

## Supplemental Note 1: Related works and paper contributions

To position DRVI within the field, we begin by discussing its relation to the broader disentangled representation learning literature. Then, we review existing disentanglement methods applied to single-cell omics and highlight the specific contributions of this work.

### Disentangled representation learning

Disentangled representation learning methods can be categorized according to their architecture, representation structure, and supervision signal [27]. Common disentangled representation learning frameworks utilize generative models such as variational autoencoders (VAEs), generative adversarial networks (GANs), or diffusion models [27]. These models are primarily applied in computer vision, employing various regularization losses to encourage disentaglement. Indeed, unsupervised disentanglement is theoretically impossible without inductive bias on the model or data [32]. The empirical success of disentanglement in computer vision architectures can therefore be attributed to inherent inductive biases, such as the spatial priors of convolutional neural networks (CNNs). Because generative models used for single-cell analysis typically employ unconstrained multilayer perceptrons and lack spatial or structural inductive biases, directly transferring computer vision paradigms to single-cell omics is challenging. To address this challenge, DRVI introduces a specific architectural inductive bias: an additive decoder architecture [34] integrated with log-sum-exp pooling to enforce dimension-wise disentanglement. Although implemented here within a variational autoencoder framework, this architectural inductive bias is generalizable and can be incorporated into other generative models, such as those reviewed in [27].

Disentangled latent space models can be categorized by the granularity of their representations. Block-based models partition the latent space into independent sub-spaces or blocks, each representing a high-level process, though the individual dimensions within each block remain entangled. In contrast, dimension-wise models disentangle each latent dimension individually to represent a single underlying process. DRVI falls into this second category, while also offering users the flexibility to perform block-based disentanglement by adjusting default parameters.

The theoretical impossibility of unsupervised disentanglement has motivated many frameworks to incorporate supervised or weakly supervised information [27, 97]. Consequently, existing single-cell disentanglement methods often require auxiliary annotations or pathway information as supervision. By contrast, DRVI relies on a structural inductive bias to learn fully disentangled, unsupervised representations, eliminating the need for labels or prior knowledge of the underlying biological processes.

### Disentanglement in single-cell omics

We review existing single-cell disentanglement methods, focusing on their architectural assumptions and latent space structures. Within block-based frameworks (less-relevant to our work), methods like MIDAS [98] and inVAE [99] separate biological signals from technical batch effects into distinct latent blocks. ContrastiveVI [100] and SC-VAE [101, 102] partition the latent space to isolate perturbation-specific responses from background variations. Furthermore, scDisInFact [103] and BIOLORD [104] construct multiple latent blocks corresponding to specific biological or technical covariates. Nevertheless, all these methods require the same information they seek to isolate in latent blocks as supervision.

To achieve dimension-wise disentanglement under the constraints of Locatello’s impossibility theorem [32], single-cell models typically constrain the search space or incorporate auxiliary biological supervision [34, 105–108]. For example, sVAE [28], SAMS-VAE [29], and sVAE-ligr [109] leverage experimental perturbations as supervision to guide dimensional assignments. Similarly, pmVAE [31] and expiMap [30] restrict decoder connections using predefined gene sets or pathways. Spectra [19] combines cell-type annotations and gene-gene interaction networks to guide a sparse, non-negative matrix factorization framework. Other models enforce structural constraints directly; BAE [110] and MOFA+ [35] enforce sparsity on linear components, LIGER [36] utilizes non-negative matrix factorization, and scETM [90] implements a linear topic model decoder. Unconstrained unsupervised methods, such as *β*-TCVAE [111] and MICHIGAN [89], attempt to achieve disentanglement solely through modified VAE objectives without architectural constraints. However, consistent with theoretical limits [32], our benchmarks confirm that these unconstrained models fail to achieve meaningful disentanglement in single-cell settings. We summarize these methods and their representational characteristics in Supplementary Table 1.

Dimension-wise disentanglement yields interpretable latent factors, a goal shared with topic modeling and matrix factorization [36, 90, 112]. Topic modeling methods often assume that cells are sampled from disjoint topics with probabilities that sum to one (usually modeled by the Dirichlet distribution) [90]. Consequently, while topic modeling approaches can identify dominant competing signals (interpretable as soft clustering), they may be unable to identifying secondary or shared signals. In contrast, disentangled representation learning allows multiple parallel processes to coexist within a cell. Factorization models represent the observational matrix as the product of two low-dimensional matrices [35, 36, 90, 112]. While the linearity constraint alone isn’t enough to mitigate the well-known unidentifiability issue [97], additional assumptions such as sparsity [35] or non-negativity [36] of loadings can potentially enforce disentanglement and facilitate interpretation.

### Methodological contributions

Beyond the results presented in the main text, we contribute the following methodological advancements:

- **Integrating Additive Decoders with Deep Generative Models**: We successfully integrate Deep Generative Models for single-cell data (specifically, scVI [12]) with additive decoders [34]. We demonstrate that naive integration fails to disentangle single-cell latent spaces, highlighting the necessity of our proposed structural modifications.
- **Proposing a Novel Pooling Function**: We introduce neural network architecture where each decoder subnetwork was followed by a shared activation function *ψ* = exp, and we formulated a more practical implementation as LSE pooling function. This modification (that still remains within the additive decoders framework) is novel, effective, and not discussed in [34] (Supplementary Fig. 26). We justify this choice biologically and technically, showing it models independent biological processes additively in count space.
- **Developing a Numerically Stable and Efficient Architecture**: We present practical modifications to ensure stable, large-scale VAE training. These include a log-space parametrization of the Negative Binomial likelihood (Supplementary Note 8) and efficient deep learning module to enable parallel processing of MLPs.
- **Efficient Interpretability Pipeline**: We introduce an efficient interpretability pipeline that allows for the interpretation of latent factors in terms of gene scores. The runtime of this algorithm is independent of the number of cells.

## Supplemental Note 2: On multiplicative behavior of current factorization models and autoencoders

Let ***Z*** ∈ ℝ^*N*×*K*^ and ***W*** ∈ ℝ^*K*×*G*^ represent the activity and weight matrices in a factorization model that represents log-transformed expressions. Typically, the log-transformed count matrix is modeled as the product of these matrices:

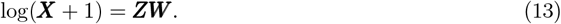

Here, ***Z***_*n,k*_ is interpreted as the activity of factor *k* in cell *n*, and ***W*** _*k,g*_ as the weight of gene *g* in factor *k*. The term ***Z***_*n,k*_***W*** _*k,g*_ represents the log-space contribution of process *k* to gene *g* in cell *n*. Mapping this representation back to the count space yields:

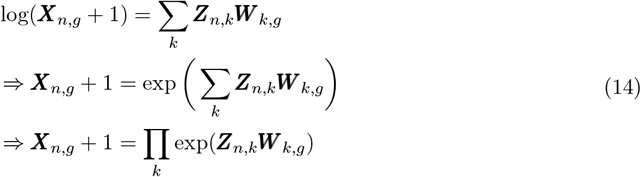

As we see, the values ***Z***_*n,k*_***W*** _*k,g*_ multiply in count space. This formulation shows that log-space additivity implies that when multiple independent processes affect the same gene, their joint effect is multiplicative in the original count space.

We can draw the same conclusion for autoencoder models. Let ***Z***_*n,k*_ be the latent factors in a variational autoencoder with a linear decoder. As in the matrix factorization case, let ***W*** be the weight matrix in the last layer of the decoder. Interpreting ***Z***_*n,k*_ as the activity of factor *k* in cell *n* and ***W*** _*k,g*_ as the signature weight, the same multiplicative aggregation in count space applies.

In the case of deeper decoders, one can define ***Z***_*n,k*_ as the value for cell *n* in node *k* in the last hidden layer of the decoder (instead of latent variables). Considering these values as “hypothetical” factors indicating activity of hypothetical factor *k* in cell *n*, we can conclude that autoencoders with MLP decoders also rely on the assumption that some activities (not latent activities but activities in the last layer) aggregate through multiplication in count space.

## Supplemental Note 3: On the additivity of underlying biological processes in count space

We propose count-space additivity as a physically motivated alternative to log-space additivity, which is standard in current factor models. Log-space additivity assumes that when two biological processes regulate a shared gene (probably at different timepoints), their joint effects accumulate multiplicatively in count space (Supplementary Note 2).

We offer a mechanistic justification for additivity in count space. Considering the decay of expressed RNAs within each process (as a modeling choice, RNA degradation is not modeled explicitly due to the requirement for more complex models and data limitations), if two biological processes affect the same gene in different time frames, the newly transcribed genes will result in an additive effect on the total RNA count and therefore add up in count space when sequenced (Supplementary Fig. 23 top). This aligns with biological scenarios such as developmental bifurcations, induced perturbations, and stress responses, where newly transcribed RNAs contribute additively.

Furthermore, this assumption also aligns with many technical processes and artifacts during the generation of scRNA-seq data. For instance, the combined expression of doublet cells, background contamination, hemoglobin (HBB) gene contamination, and even residual counts from neighboring cells due to imperfect segmentation in spatial omics all contribute additively to the observed count space (Supplementary Fig. 23 bottom).

Thus, in contrast to prior models, our additive assumption provides a more plausible framework for modeling single-cell count data.

Here we note that the task of interest in disentanglement is to identify the underlying biological processes, without inferring causal relationships between them. For instance, if process *P*_1_ activates *P*_2_, which subsequently feedbacks *P*_1_, each process is defined solely by its direct transcriptional targets. Accordingly, while we are interested in identifying P1 and P2, we are not interested in identifying their causal relationships, although disentanglement may help in this downstream task. In our framework, we define the output of a process solely based on its direct transcriptional effects, disregarding downstream or propagated causal effects. This assumption allows us to model the activity of individual processes additively. Inferring downstream causal links or networks among these factors remains a separate task for downstream analysis. Finally, in cases where the intersection of two programs induces a synergistic transcriptional state (e.g., via transcription factor heterodimerization), their joint effect can be modeled as a distinct, entirely new factor within our framework.

## Supplemental Note 4: Formal definitions and notations

To formalize the disentanglement, we present the definitions and notations below (adapted from [34]).

### Partition

Given a set 𝒜, a set 𝒫 is a partition of 𝒜 if the elements of 𝒫 are mutually exclusive (∀*B* ≠ *B*′ ∈ 𝒫, *B* ∩ *B*′ = ∅) and their union covers all elements of 𝒜 (u_∈𝒫_ *B* = 𝒜).

### Projection

The projection of a point *z* ∈ ℝ^*K*^ onto a block of dimensions *B* ⊆ {1, …, *K*} is defined as 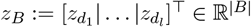, where *d*_*i*_ are the ordered elements of *B* = {*d*_1_ *<* · · · *< d*_*l*_}.

### Disentanglement

Let Ƶ ⊂ ℝ^*K*^ be the latent space, and let be 𝒫 a partition of the latent dimensions. Given a ground-truth generative process defined by a decoder *f* and a learned decoder 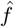 optimized on samples generated by *f*, we say 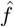 is disentangled with respect to *f* and partition 𝒫 if there exists a permutation of latent dimensions *π* and a function *v*: ℝ^*K*^ → ℝ^*K*^ such that for all *z* ∈ Ƶ:

- The function *v* does not mix different blocks of the partition 𝒫. Formally, for all distinct blocks *B*≠ *B*′ ∈ 𝒫 and all *i* ∈ *B, j* ∈ *B*′, the (*i, j*)-th entry of the Jacobian *J*_*v*_(*z*) is zero.
- The decoders satisfy 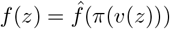, where *π*(*z*_1_, …, *z*_*K*_) = (*z*_*π*(1)_, …, *z*_*π*(*K*)_).

Under this definition, 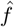 is equivalent to *f* up to blockwise invertible transformations and permutation of blocks. Disentanglement is dimension-wise when evaluated with respect to the discrete partition 𝒫 = {{1}, {2}, …, {*K*}}.

### Statistical Independence

Latent factors (or blocks of dimensions) representing a latent distribution *p*_*z*_(*z*) are statistically independent if, for any two blocks *B*≠ *B*′:

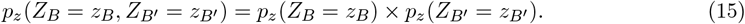

Unlike disentanglement, which evaluates the functional mapping of the decoder, statistical independence is a property of the latent probability distribution *p*_*z*_(*z*).

## Supplemental Note 5: Differences between disentanglement and independence

In the disentangled representation learning literature, two perspectives exist: the classical view assumes statistical independence among latent factors, whereas the modern view accommodates causal relationships between them [27]. In this work, we define disentanglement as a one-to-one mapping between the underlying biological processes and the learned latent factors. This formulation permits correlations and dependencies between factors, inherent in biological systems. Here, we highlight the distinctions between disentanglement and statistical independence using illustrative examples. In these examples, the data consists of images containing one red ball and one blue ball. We can then model this data using a generative process with two factors: one representing the height of the red ball (*A* ∈ [0, 1]) and the other representing the height of the blue ball (*B* ∈ [0, 1]).

### Example 1: Alignment of Independence and Disentanglement

When the underlying generative factors are physically independent and the training manifold is fully and uniformly sampled (*A, B* ∼ Uniform(0, 1)), disentanglement and statistical independence align. A model that successfully disentangles *A* and *B* will naturally yield statistically independent latent dimensions (Supplementary Fig. 22a). In this setting, both dependency-penalizing metrics (such as the Mutual Information Gap, MIG) and disentanglement-measuring metrics (such as the Latent Matching Score, LMS) are maximized.

### Example 2: Disentanglement without Independence (Selection Bias)

Consider a scenario where the physical mechanisms governing *A* and *B* are independent, but the training data is subject to selection bias. A model that correctly disentangles *A* and *B* will yield correlated latent factors because of the biased support of the training data (Supplementary Fig. 22b). This arises because statistical independence is a property of the observational distribution, whereas disentanglement is a property of the generative mapping. In single-cell biology, an example of selection bias is when two independent perturbations are observed, but their combination is absent from the experimental design or the coexistence of two perturbations results in cell apoptosis.

### Example 3: Disentanglement without Independence (Causal Dependency)

Assume the red ball is physically constrained to remain below the blue ball. The generative process is then: *A* ∼ Uniform(0, 1) and *B*|*A* ∼ Uniform(*A*, 1). A disentangled model will map *A* and *B* to distinct latent dimensions, but these dimensions will be correlated (Supplementary Fig. 22c).

Under such causal dependencies, dependency-penalizing metrics (like MIG) penalize correct models, whereas disentanglement-measuring metrics (like LMS) remain maximized.

Biological systems typically exhibit such causal structure: one program may induce another, or competing programs may restrict lineage choices. For example, proliferation depends on cell type or cytokine signaling.

### Example 4: Independence without Disentanglement (Over-compression)

Finally, we consider an entangled model that achieves statistical independence. Suppose we introduce a third object, represented by a binary process *C* ∈ {0, 1} (e.g., representing shape: square vs. triangle). A disentangled model should assign *A, B*, and *C* to three distinct latent dimensions. However, an over-compressed model might assign *A* to one dimension and a mixture of *B* and *C* to another. Such a model is entangled, yet the two latent dimensions remain statistically independent (Supplementary Fig. 22d). This demonstrates that over-compressed models can satisfy statistical independence at the cost of disentanglement. In this case, dependency-penalizing metrics (like MIG) are maximized, whereas disentanglement-measuring metrics (like LMS) correctly penalize the entanglement of multiple processes in a single dimension.

## Supplemental Note 6: Identification of the remaining non-cell-type factors for the HLCA

In this section, we annotate and characterize the remaining non-cell-type latent factors of the HLCA study not detailed in the main text. This characterization is based on the cell types these factors are active in (Supplementary Fig. 31) and their associated gene attribution scores (Supplementary Fig. 30). The factors are presented in the order of their appearance in Fig. 3b. To support these annotations, supporting evidence from the published literature together with enrichment results (Supplementary File 2) are provided.

### DR 26+

This factor captures an inflammatory program active in myeloid cells. The primary gene associated with this factor is EREG, co-expressed with key inflammatory mediators including IL1B, IL1A, CXCL8, and CXCL3. Gene Set Enrichment Analysis (GSEA) suggests an association with interleukin-10 signaling pathways.

### DR 29+

This factor captures a transcriptional gradient within the ciliated cells. DNAH11, DNAAF1, DNAH12, and DNAH6 are among the top identifiers of this factor, expressed across ciliated cells, with significantly higher expression observed in the upper gradient of DR 29+. GSEA confirms a strong association with cilium assembly and structure.

### DR 14+

The gene program for DR 14+ is dominated by human leukocyte antigen (HLA) class II genes, including HLA-DRA, HLA-DRB1, and HLA-DPA. GSEA associates this program with the MHC class II protein complex.

### DR 19-

This factor is active within epithelial cells, particularly nasal goblet cells. The associated gene program is characterized by KLK7, TMPRSS11B, SPINK7, ECM1, SCEL, and KLK6. GSEA suggests an association with esophagus and oral mucosa. A definitive biological interpretation remains to be determined.

### DR 22+

FOLR2, F13A1, STAB1, CCL13, and SELENOP are the main identifiers of the gene program associated with DR 22+. These genes indicate a perivascular population of macrophages [51, 113]. High activity of this factor in cells annotated as Interstitial Mph perivascular confirms this relevance.

### DR 23-

SERPINB3 and SERPINB4 are the top identified gene for DR 23-, active in epithelial cells. Although the coexpression of these two genes is biologically relevant [114], we could not find a specific biological explanation for the subpopulation of cells active in this factor.

### DR 27+

The activity of DR 27+ indicates the expression of PLAU, its receptor PLAUR, and an elevated expression of CLDN4 (Claudin-4) in airway epithelium. PLAU and CLDN4 are separately suggested as markers of lung injury, infection, and barrier repair [115–117]. GSEA suggests an association with dissolution of fibrin clot and regulation of wound healing.

### DR 28-

This factor is active in a subset of airway epithelial cells, including Hillock-like cells. GSEA on the associated gene signature including KRT6B, KRT13, KRT16, and KRT6A, suggests a keratinization program. Previous studies ([118, 119]) have employed a diverse set of KRT genes to characterize epithelial subpopulations. In particular, KRT16 and KRT6A are among the markers of certain Hillock-like cell populations.

### DR 30+

EDN1, ERRFI1, DST, and TNFRSF12A are the main indicators of DR 30+. This factor is more active in basal cells, with a significant upregulation of cells from the Nawijn_2021 dataset. A clear technical or biological explanation for this factor remains open.

### DR 33-

This factor indicates a subset of fibroblasts and smooth muscle cells identified by a gene program composed of genes such as CRISPLD2, IL6, and ADAMTS4. IL-6 is a pro-inflammatory cytokine, reported to induce expression of ADAMTS4 in rheumatoid arthritis samples [120]. Conversely, CRISPLD2 is a known inhibitory regulator of IL-6 [121, 122]. The coexpression of these genes suggests a dynamic of IL-6 regulation. The exact role of this process needs further experiments.

### DR 33+

C1QA, C1QB, C1QC, and TREM2 are the top indicators of the DR 33+, Suggesting that this factor represents TREM2 and the C1Q complex in myeloid cells, including TREM2+ macrophages [123–125].

### DR 36-

This factor is characterized by the expression of LYPD3, CALML3, AQP3, TACSTD2, and LY6D. The precise biological function of this program remains uncharacterized.

### DR 37-

The single-driver gene for this factor is ANKRD36C, which is expressed in a subset of goblet cells. This factor is active primarily in bronchial goblet cells, but its specificity to particular datasets suggests a potential batch-specific or localized artifact.

### DR 38+

DR 38+, characterized by MKI67, TYMS, and PCLAF expression, is associated with cell-cycle processes as identified by GSEA. This aligns with the cell-type annotations (Fig. 3b).

### DR 40+

This factor highlights a subset of airway epithelial cells, with RARRES1, CXCL6, SER-PINB4, and IL17C as the key drivers of its gene expression program. While GSEA suggested an IL-17 signaling pathway, we found the relationship between these genes and the pathway to be weak. As a result, we were unable to identify a meaningful explanation for this factor.

### DR 41+

This factor identifies a subset of epithelial cells, with MMP1, STC1, MMP13, and MMP10 as their top relevant genes. Accordingly, this factor highlights the activity of certain proteins in the Matrix Metalloproteinase family, suggesting an ongoing destructive process [126].

### DR 42+

This factor is active across two distinct cell lineages (Supplementary Fig. 31). In the myeloid compartment, it associates with OLFM4, S100A7, and DEFA3, suggesting a neutrophil signature [127, 128]. In the epithelial compartment, it is characterized by S100A7 and small proline-rich protein genes (SPRR1B, SPRR2D, SPRR2E, SPRR2F, SPRR3). GSEA suggests keratinization and formation of the cornified envelope for this factor. The co-existence of these two distinct programs in a single factor suggests a partial disentanglement failure, likely due to an insufficient number of latent dimensions to resolve all independent components.

### DR 43+

This factor captures a general inflammatory program characterized by CCL20, CXCL2, CXCL3, CSF2, CSF3, and ICAM1. GSEA indicates activation of the TNF, IL-17, and IL-10 signaling cascades, representing an inflammatory response [129].

### DR 44+

This factor indicates a subset of myeloid cells characterized by but not limited to CCL3, CCL4, CCL3L1, CCL4L2, CXCL3, IL1A, and IL1B genes. CCL3 (MIP-1*α*) and CCL4 (MIP-1*β*) and their paralogs CCL3L1 and CCL4L2, are proinflammatory chemokines that attract immune cells [130]. IL-1*β* is an inflammatory cytokine that activates cells to produce more CCL3 and CCL4 chemokines [131]. GSEA describes this factor by Interleukin-10 signaling, inflammatory response, and cytokine activity. This subpopulation of myeloid cells is of biological and clinical interest [132]. It is worth mentioning that DR 26+ complements this by highlighting another population of myeloid cells undergoing interleukin-10 signaling pathway.

### DR 49-

With HBA2, HBA1, and HBB as its top identifiers, GSEA analysis suggests that DR 49-represents the hemoglobin metabolic process. While this process is expected to be most active in erythroid cells that are not present in the HLCA, we observed non-negligible expression of hemoglobin genes in other cell types. The expression of hemoglobin genes in the HLCA could be attributed to technical or biological factors [133, 134].

### Dim DR 50+

SAA1 and SAA2 are the main identifiers of the dim DR 50+. So, we can conclude that this factor presents the activity of SAA as an inflammatory marker.

### DR 60-

The nonlinear gene program associated with DR 60-includes multiple genes such as ISG15, IFIT1, IFI44L, and IFI6. GSEA describes this factor by interferon alpha/beta signaling and response to virus pathways.

## Supplemental Note 7: Considerations for encouraging disentanglement

Here, we discuss our considerations for encouraging disentanglement in DRVI. Theorem A.22 of the interaction asymmetry framework [76] provides criteria under which disentanglement is guaranteed:

### (I) Number of processes

Disentanglement proofs typically assume that the number of latent dimensions matches the number of ground-truth (GT) processes. However, this number is rarely known in biological applications. To address this, we show that when excess latent dimensions are provided, the model saturates and the extra dimensions vanish (see Results). This allows users to specify an over-parameterized number of latent dimensions during training. Consequently, we assume that vanished dimensions are pruned before assessing other disentanglement criteria.

### (II) 1^*st*^ Order Interaction Asymmetry

First-order interaction asymmetry requires simple interactions across splits (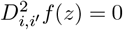 for dimensions *i, i*′ belonging to different splits) and higher-order interactions within splits (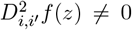 for dimensions within a split). Additive decoders have at most first-order interactions between splits (Definition 3.3 of [76]). Accordingly, we require at least second-order interactions within each split of our model. Average pooling does not necessarily enforce such nonlinearity within each split. However, because the exponential activation function (*ψ* = exp) has infinite differentiability, using this shared activation function in LSE pooling heuristically introduces at least second-order interactions within each split.

### (III) 1^*st*^ Order Sufficient Independence

This is the second main criterion for disentanglement, which depends on the choice of the activation function *ψ*. For average pooling, we cannot make any guarantees. For DRVI with LSE pooling, the criterion (Definition A.9 of [76]) takes the form:

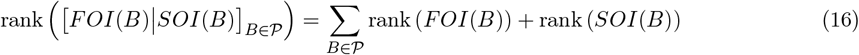

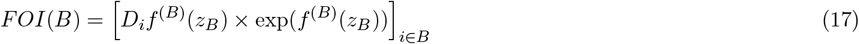

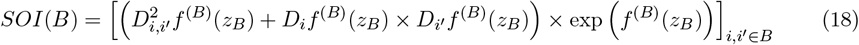

where *FOI*(*B*) and *SOI*(*B*) represent matrices of first-order and second-order interactions, and 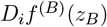 and 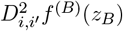 denote the first-order and second-order derivatives of *f* ^(*B*)^ at point *z*_*B*_.

We assume that each ground-truth biological process (block *B*) possesses a distinct set of marker genes. Without loss of generality, we can restrict the feature space to these markers. The term exp(*f* ^(*B*)^(*Z*_*B*_)) appears in all elements of both the first-order interaction (*FOI*(*B*)) and secondorder interaction (*SOI*(*B*)) matrices. Since the aggregation of these values (∑ *B*∈𝒫 exp(*f* ^(*B*)^(*Z*_*B*_))) will reconstruct the expression matrix and the outputs of each split of the generative process (exp(*f* ^(*B*)^(*Z*_*B*_))) are non-negative, they are nonzero if and only if the gene is a marker of the split. Thus, for each split *B*, only the rows corresponding to its marker genes contain nonzero values, meaning the matrices [*FOI*(*B*) *SOI*(*B*)] share no common nonzero rows across different splits. Accordingly:

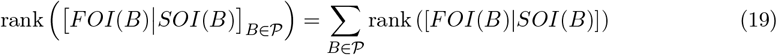

This significantly narrows the gap between the two sides of the sufficient independence criterion. In addition to this theoretical justification, we analyzed a DRVI model trained on the Immune dataset using 32 and 64 latent dimensions to construct the [*FOI*|*SOI*] matrix. We empirically observed that the non-vanished dimensions of DRVI form full-rank sufficient independence matrices (Supplementary Fig. 27).

### (IV) Diffeomorphism

The decoder is assumed to be a *C*^2^-diffeomorphism (*f* and its inverse being *C*^2^). We employ exponential linear units (ELU) [135] as the activation function in the subnetworks defining *f*, which offers a *C*^1^-smooth alternative to ReLU. However, *f* still does not satisfy the formal *C*^2^-diffeomorphism requirement. In practice, we are not aware of any deep generative model that strictly satisfies this mathematical criterion.

### (V) Reconstruction Loss

Since we estimate the parameters of the noise distribution, solving the reconstruction problem cannot be formally guaranteed. Nevertheless, achieving low reconstruction error is sufficient in practice, and this assumption is usually relaxed.

### (VI) Domain Assumptions

We assume that our domain is regularly closed. While path connectivity is typically required to derive global disentanglement from local disentanglement, this assumption often falls short in non-developmental data. However, latent space sampling during VAE training promotes connectivity and thus encourages global disentanglement. Additionally, the KL divergence term in the loss function actively penalizes redundant factors, encouraging the model to allocate distinct biological processes to individual latent dimensions when sufficient dimensions are available (Supplementary Fig. 28).

## Supplemental Note 8: Log-space Parametrization of the Negative Binomial log-likelihood

In generative models using the Negative Binomial distribution, the decoder typically outputs parameters in log space, which are then exponentiated to count space. The Negative Binomial log-likelihood is then evaluated as:

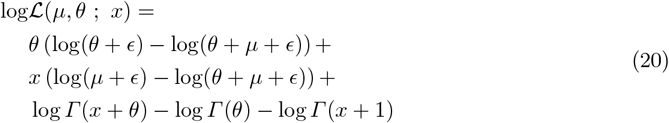

where *µ* and *θ* represent the mean and dispersion parameters of the distribution, *x* is the observed count, *Γ* is the Gamma function, and *ϵ* is a small constant for numerical stability.

This approach involves exponentiating the log-space outputs of the decoder only to apply logarithms during likelihood calculation, which can degrade numerical precision. To address this, we reformulate the Negative Binomial log-likelihood directly in terms of the log-space parameters. First, we use the Softplus algebraic identity to simplify the ratio term:

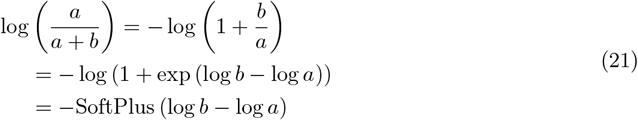

This allows us to rewrite the log-likelihood as:

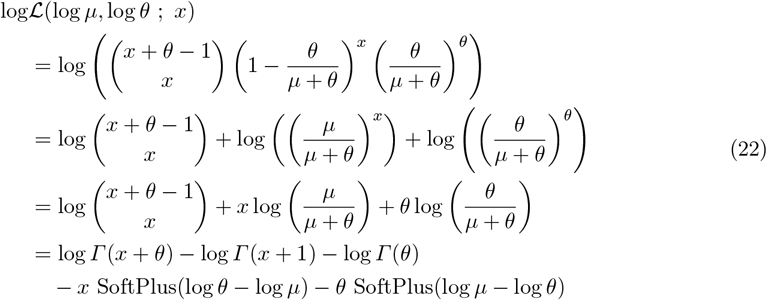

To ensure numerical stability, we add a small constant *ϵ >* 0 to the arguments:

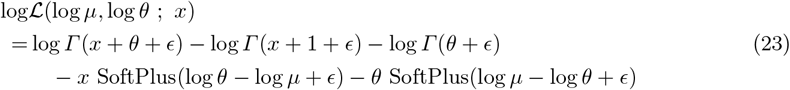

where log *µ* and log *θ* are the parameters output by the network in log space. Note that while both *θ* and log *θ* appear in the final expression, the mean parameter *µ* is parameterized solely via log *µ*. This increases numerical precision during training, as log *µ* aggregates the outputs of multiple decoder subnetworks and its gradients are central to optimization.

## Supplemental Note 9: Interpretation of Latent Factors via Additive Decoder Architecture

To interpret the latent factors, we leverage the additive decoder architecture to analyze each decoder subnetwork independently. We begin with average pooling as the baseline case. For DRVI-AP, the generative process is defined as:

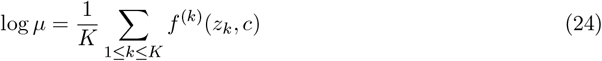

where *µ, z*_*k*_, *f* ^(*k*)^, and *c* indicate the mean parameter of the generative distribution, *k*th latent factor, *k*th decoder subnetwork in the additive decoder, and the covariate vector, respectively. In this case, for gene *g* and factor *i* of the latent space we have:

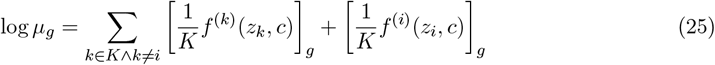

When perturbing *z*_*i*_, the LFC of *µ*_*g*_ solely depends on *f* ^(*i*)^(*z*_*i*_, *c*). We take the mean over covariates to consider the average effect over different batches. Since *f* ^(*i*)^ is a nonlinear function, we are interested in the maximum effect of this latent factor. Accordingly, we maximize the effect over different choices of *z*_*i*_. In practice, we observe factors encoding two different concepts in their negative and positive directions. So, we split these two directions for better interpretability. Consequently, the effect of dimension *Z*_*i*_ on gene *g* is defined as:

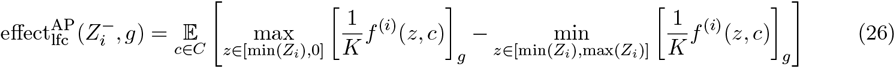

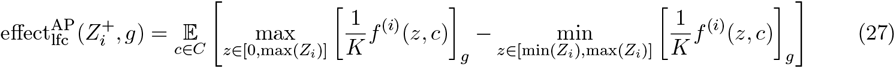

These effects can be efficiently approximated by sampling from covariates and traversing (one-dimensional) discrete values over each latent factor.

For LSE pooling function, the generative process is:

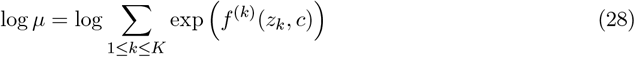

where *µ, z*_*k*_, *f* ^(*k*)^, and *c* are defined as in the average pooling case.

Accordingly, for each gene *g* and factor *i* of the latent space we have:

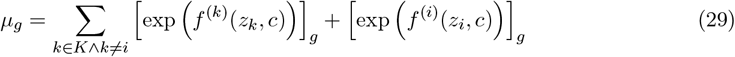

Here, the effect of perturbing *z*_*i*_ in count space solely depends on *f* ^(*i*)^(*z*_*i*_, *c*). Formally:

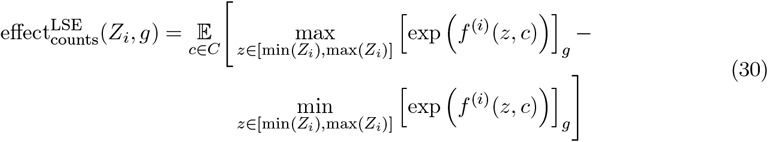

However, since we are usually interested in the LFC rather than the count space effect, we also have to consider the effect of other latent factors. The LFC will be more significant if other factors do not affect the target gene and will be smaller otherwise. Here, we make use of the additiveness property to write the minimum/maximum effect of a group of factors as the sum of the minimum/maximum effect of each individual factor. Accordingly, to calculate the maximum possible effect, we have:

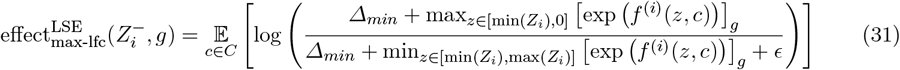

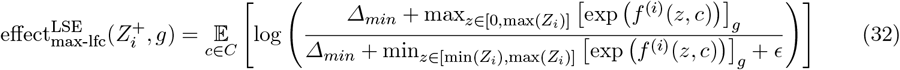

Where *Δ*_*min*_ is defined as 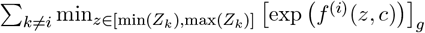 and *ϵ* is a small number defined for numerical stability.

Likewise, for the minimum effect, we define:

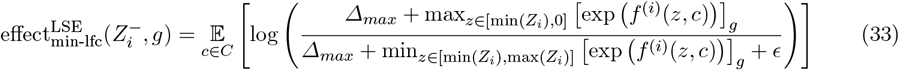

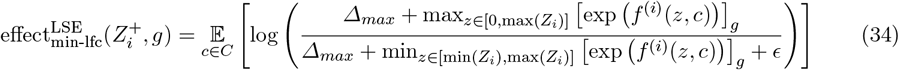

Where *Δ*_*max*_ is defined as 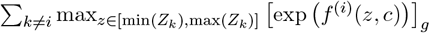.

All the minimum and maximum values over decoder subnetworks can be efficiently approximated by traversing discrete ranges over the latent factors. Averaging over categorical covariates can be done by sampling. The same approach applies to DRVI with any other pooling function.

The minimum and maximum effects measure two different aspects of the latent factors.

The effect_max-lfc_ measures the maximum effect of a latent factor regardless of the baseline expression. For example, for two genes with similar average expression, changing a log-mean parameter of a gene from 0.1 to 0.5 has the same score as changing the other from 1 to 5. The effect_min-lfc_ score is robust to such cases by taking the maximum expression of other factors into account as a baseline. Still, it drops significantly if a gene participates in multiple latent factors. Consequently, we combine the two scores by multiplying them. Formally: 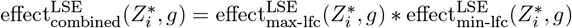 where 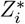 can be either 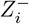 or 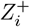. Additionally, we remove all the insignificant effects meeting any of the following criteria:

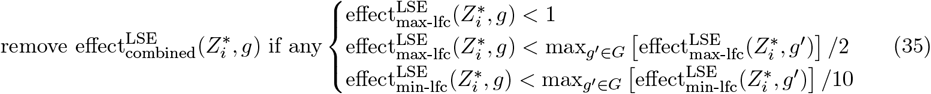

We found the mentioned interpretability pipeline accurate enough for our analysis. However, further tuning of the pipeline and the combining logic can be a follow-up, which is out of the scope of the current study.

Since the introduced approach directly reflects the model’s understanding of the generative process, it is limited to the set of genes included in the training data. Alternatively, one can perform differential expression (DE) analysis to identify the genes involved in a nonlinear gene program beyond the genes used for model training. To this end, cells can be categorized into high-valued and low-valued groups based on a latent factor, or latent factors can be incorporated into the design matrix of a regression model for DE testing. While DE analysis is a well-established approach, it may highlight genes that are not specific to a process but are simply expressed at higher-than-average levels in the target population. In addition, lowly expressed genes may be missed.

Besides looking up the literature and marker databases, we used the Python interface of the gProfiler service [136] for GSEA analysis. To account for the biological bias introduced by the selection of HVGs, they were given as the reference set to the gProfiler.

## Supplemental Note 10: Ordering of Latent Factors

Unlike Principal Component Analysis (PCA), where factors are ordered by explained variance, the latent dimensions of deep autoencoders (as well as many factor models such as ICA, MOFA, and LIGER) are inherently unordered. We exploit the additive architecture of DRVI to formulate a heuristic that orders latent factors by their contribution to the reconstruction (as in PCA). We evaluate the impact of removing each factor on the predicted mean parameters.

Let 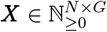 represent the observed count matrix, ***Z*** ∈ ℝ^*N*×*K*^ be the learned latent coordinates, and *f* be the decoder mapping. For the *k*-th latent factor, we construct the subnetwork output matrix ***S***^(*k*)^ ∈ ℝ^*N*×*G*^ where its *i*-th row is defined as:

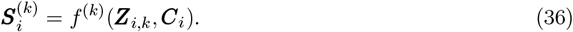

The reconstructed log-mean parameters for DRVI-AP are given by 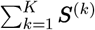. Consequently, removing the *i*-th factor alters the log-space parameter by ***S***^(*i*)^. For average pooling, we sort factors in descending order based on the Frobenius norm (or L1 norm) of their respective subnetwork matrices ***S***^(*k*)^.

For LSE pooling, the change in the reconstructed log-mean parameter upon removing the *i*-th factor is defined as:

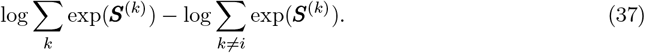

To ensure numerical stability, we add a regularizing constant *ϵ* = 1 within the log function. The reconstruction impact of factor *i* is then defined as:

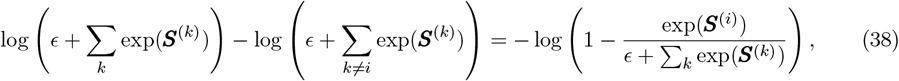

which can be efficiently computed across cells using the softmax operations.

## Supplemental Tables

**Supplemental Table 1.** Comparison of various methods for disentangling latent factors in single-cell omics.

| Method | Disentangled Entities | Supervised input | Representational constraints |
| --- | --- | --- | --- |
| DRVI (Ours) | Each latent factor | No | Additive decoders |
| β-TCVAE, MICHIGAN [89, 111] | | | Unconstrained |
| BAE [110] |  |  | Sparse **linear** encoder |
| scETM [90] |  |  | Topic model with **linear** decoder |
| MOFA / MOFA+ [35] |  |  | **Linear** decomposition and sparsity |
| LIGER [36] |  |  | **Linear** and non-negative |
| ICA |  |  | **Linear** decomposition |
| Spectra [19] | Each latent factor | Gene-gene knowledge graph and Cell-types | **Linear** decomposition and sparsity |
| expiMap [30] |  | Pathways | **Linear** decoder with pathway masks and sparsity |
| pmVAE [31] |  | Pathways | Nonlinear with pathway masks |
| sVAE-ligr [109] | Each latent factor | Perturbations / Variations of interest | No |
| SAMS-VAE [29] |  | Perturbations |  |
| sVAE / sVAE+ [28] |  | Perturbations |  |
| scDisInFact [103] | Multiple groups of latent factors | Variations of interest for each group / Perturbations | No |
| BIOLORD [104] |  | Variations of interest for each group |  |
| ContrastiveVI [100], SC-VAE [101, 102] | Two groups of latent factors | Perturbations |  |
| MIDAS [98] |  | Batch |  |
| inVAE [99] |  | Batch / [biological and technical factors] |  |

**Supplemental Table 2.**
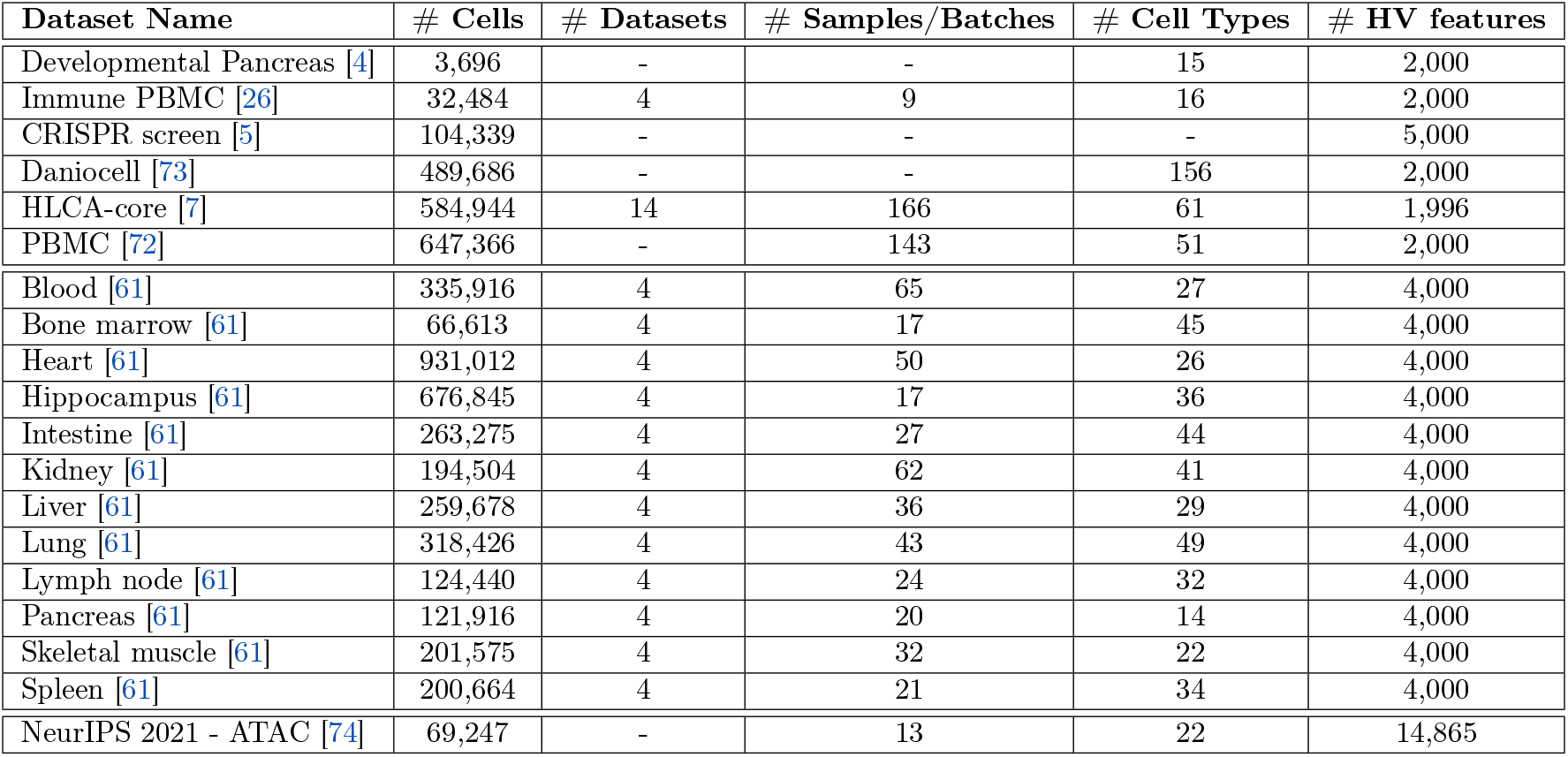
Summary of Datasets.

**Supplemental Table 3.** Summary of Model Parameters.

| Dataset Name | # Latent for all methods | Hidden Layers for DRVI / scVI | Sharing decoder subnetworks | Training epochs |
| --- | --- | --- | --- | --- |
| Developmental Pancreas | 32 | 128 | all layers | 1,000 |
| Immune PBMC | 32 | 128 | all layers | 400 |
| CRISPR screen | 64 | 512, 512, 512 | all layers | 400 |
| Daniocell | 64 | 256, 256 | all layers | 400 |
| HLCA-core | 64 | 256, 256 | all layers | 400 |
| PBMC | 64 | 256, 256 | all layers | 400 |
| 12-organ benchmarking datasets | 128 | 256, 256 | all layers | 400 |
| NeurIPS 2021 - ATAC | 64 | 256, 256 | all layers | 400 |

**Supplemental Table 4.**
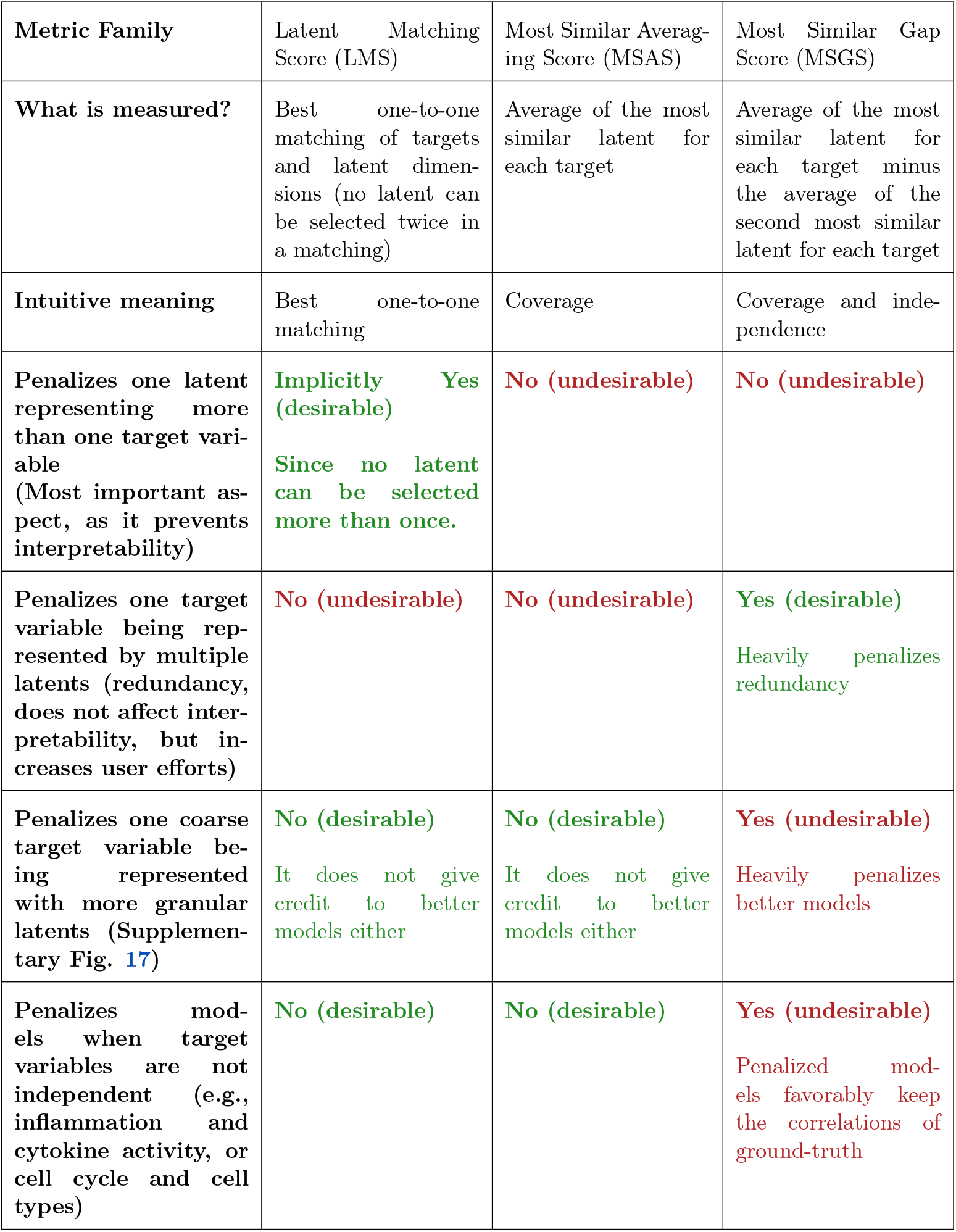
Comparison of Different Metric Families.

| Metric Family | Latent Matching Score (LMS) | Most Similar Averaging Score (MSAS) | Most Similar Gap Score (MSGS) |
| --- | --- | --- | --- |
| What is measured? | Best one-to-one matching of targets and latent dimensions (no latent can be selected twice in a matching) | Average of the most similar latent for each target | Average of the most similar latent for each target minus the average of the second most similar latent for each target |
| Intuitive meaning | Best one-to-one matching | Coverage | Coverage and independence |
| Penalizes one latent representing more than one target variable (Most important aspect, as it prevents interpretability) | **Implicitly Yes (desirable)**<br>Since no latent can be selected more than once. | **No (undesirable)** | **No (undesirable)** |
| Penalizes one target variable being represented by multiple latents (redundancy, does not affect interpretability, but increases user efforts) | **No (undesirable)** | **No (undesirable)** | **Yes (desirable)**<br>Heavily penalizes redundancy |
| Penalizes one coarse target variable being represented with more granular latents (Supplementary Fig. 17) | **No (desirable)**<br>It does not give credit to better models either | **No (desirable)**<br>It does not give credit to better models either | **Yes (undesirable)**<br>Heavily penalizes better models |
| Penalizes models when target variables are not independent (e.g., inflammation and cytokine activity, or cell cycle and cell types) | **No (desirable)** | **No (desirable)** | **Yes (undesirable)**<br>Penalized models favorably keep the correlations of ground-truth |

## Supplemental Figures

**Supplemental Figure 1:**
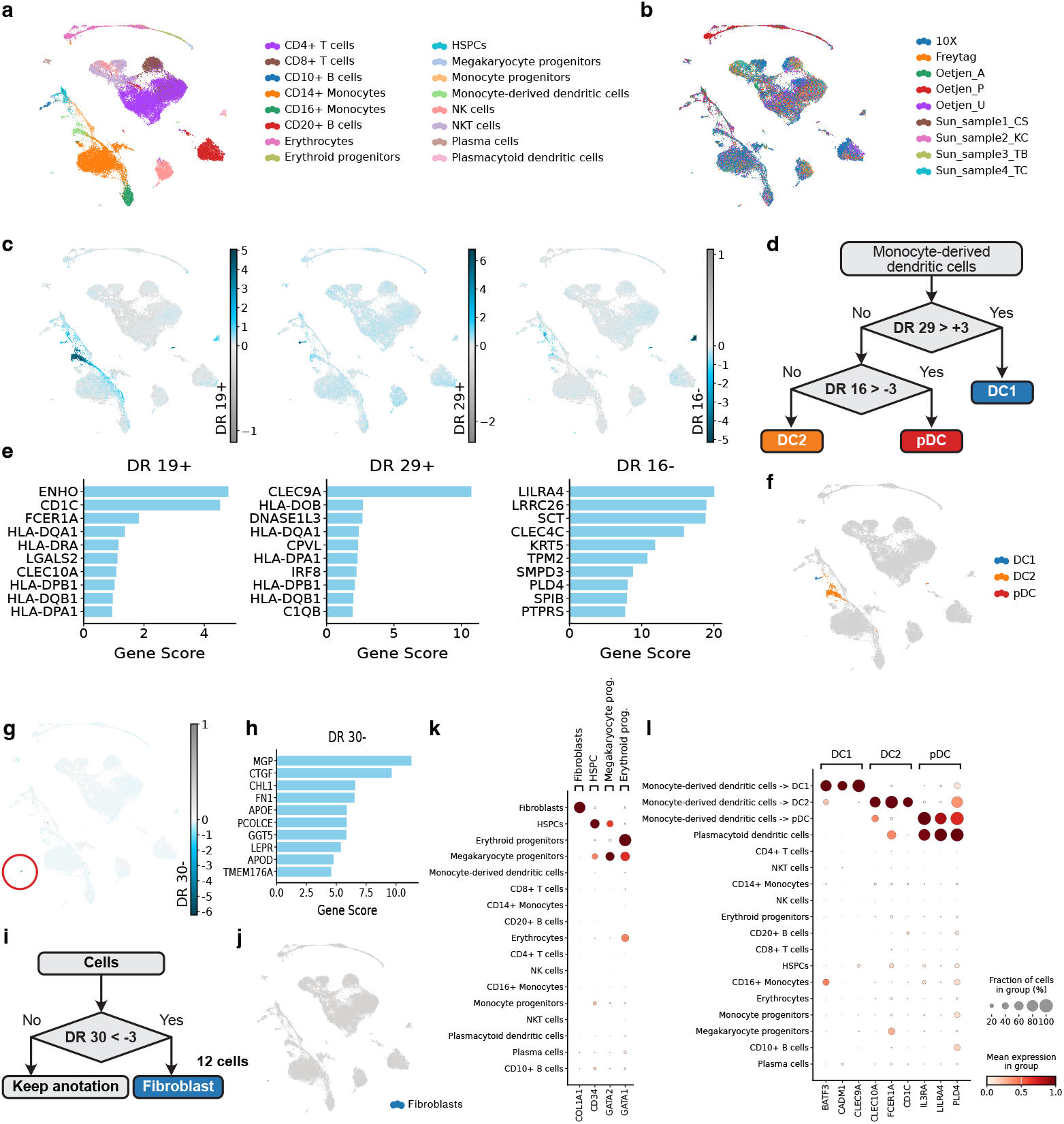
Annotation refinement and correction using DRVI in the immune dataset. **a**,**b**, UMAP representations of the integrated immune dataset, colored by batch (a) and cell type (b). **c**, UMAP visualization of latent factors identifying DC subtypes. **d**, Top genes associated with DC subtype factor. **e**, Flowchart for refining DC subtype annotations. **f**, UMAP visualization of refined DC subtype annotations. **g**, UMAP visualization highlighting the latent factor identifying Fibroblast cells. The population is encircled in red. **h**, Top genes associated with the Fibroblast factor. **i**, Flowchart for annotating Fibroblast cells. **j**, UMAP visualization of annotated Fibroblast cells. **k**, Dotplot validating new annotations for Fibroblasts, HSPCs, Megakaryocyte progenitors, and Erythroid progenitors using a representative curated marker gene for each. **l**, Dotplot of marker genes for DC subtypes for validation of DC subtype improvement. For validation, pDCs are intentionally labeled differently from the original ‘plasmacytoid dendritic cells’ annotations. DC, dendritic cells; HSPC, Hematopoietic Stem and Progenitor Cells; pDCs, plasmacytoid dendritic cells.

**Supplemental Figure 2:**
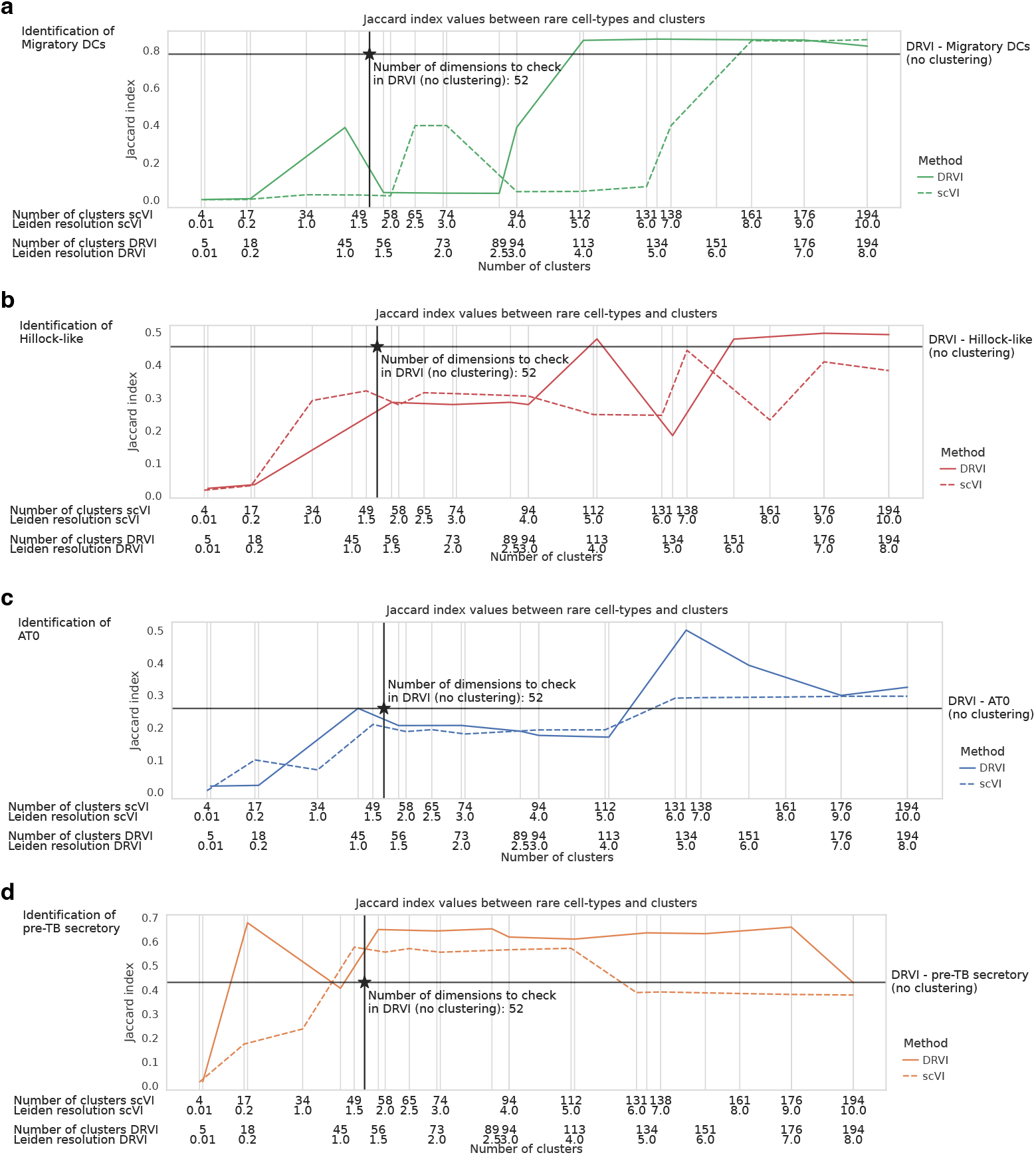
Rare cell-type identification performance of Leiden clustering in the HLCA. Leiden clustering was applied to DRVI and scVI embeddings at varying resolutions. For each cell type, we calculated the Jaccard index between the cell type and its most relevant cluster at different resolutions. Additionally, we evaluated the performance of DRVI by calculating the Jaccard index between the cell type and the separation achieved using a simple threshold on the most relevant DRVI factor (marked by a star). Migratory DCs were accurately captured by the Leiden algorithm only at the resolution of 8.0 with 161 clusters. AT0 cells were identified with comparable quality to DRVI starting from the resolution of 6.0 with 131 clusters. Hillock-like cells were captured as well as DRVI at the resolution of 7.0 with 138 clusters. Pre-TB secretory cells were captured well at a resolution of 1.5 with 49 clusters. These results demonstrate that identifying at least three rare cell types requires the creation of approximately 130 to 160 clusters using Leiden clustering. Given that DRVI can capture these cell types within its 52 non-vanished factors, we suggest DRVI as an efficient valuable tool for seed annotation and discovery of rare cell populations.

**Supplemental Figure 3:**
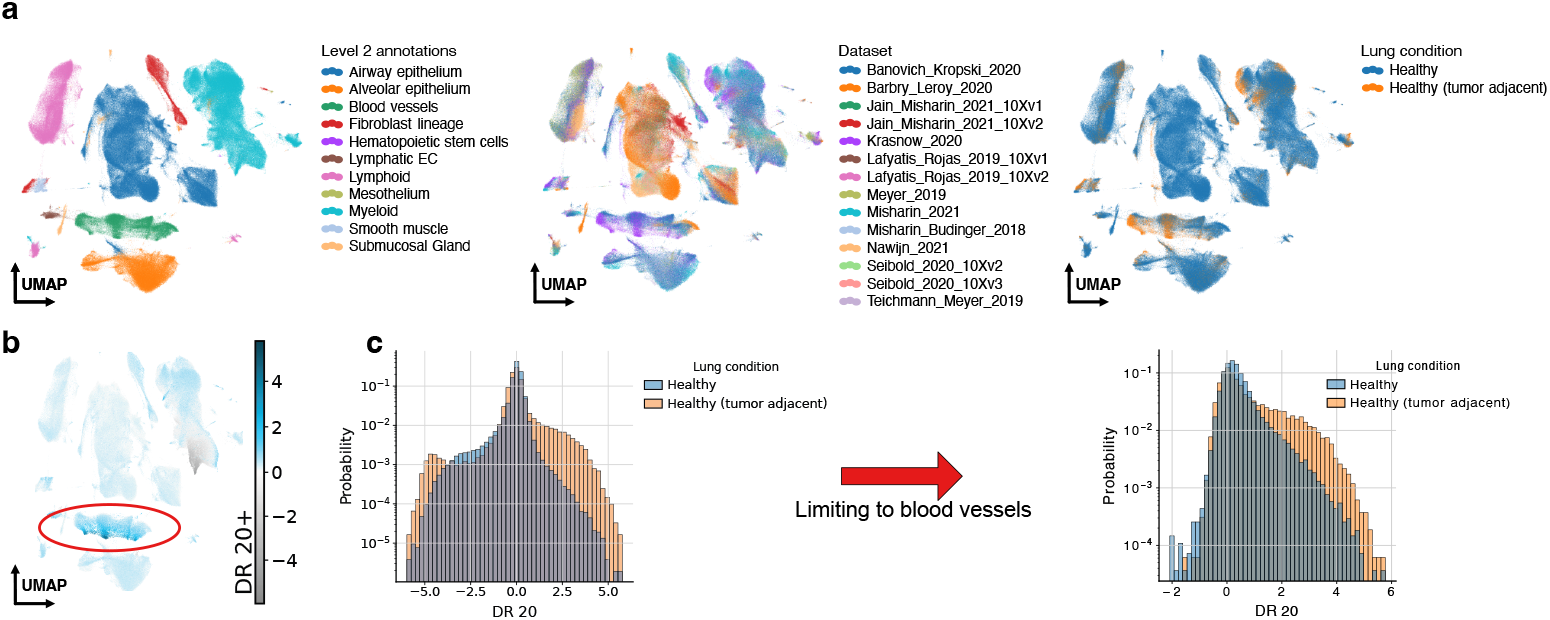
Indication of tumor-adjacent blood vessels in the HLCA. **a**, UMAPs of the HLCA integrated by DRVI, colored by second-level annotations, datasets, and tumor adjacency. Tumor-adjacent samples originate from the Krasnow_2020 dataset. By observing the blood vessels in the UMAPs, we find two groups of tumor-adjacent cells. Some integrate seamlessly with normal blood vessels, while others exhibit a degree of separation, suggesting the presence of underlying biological variation. **b**, Activity of DR 20+ on the UMAP. This factor highlights a subset of tumor-adjacent blood vessels that are distinctly separated in the UMAP, potentially representing a unique biological process or state within this cell population. **c**, Histogram of cells distributed across DR 20+, colored by tumor adjacency status before and after filtering for blood vessels. A significant proportion of cells active in the higher end of DR 20+ correspond to blood vessels located in tumor-adjacent regions, indicating a biological process or state specific to this subset of cells.

**Supplemental Figure 4:**
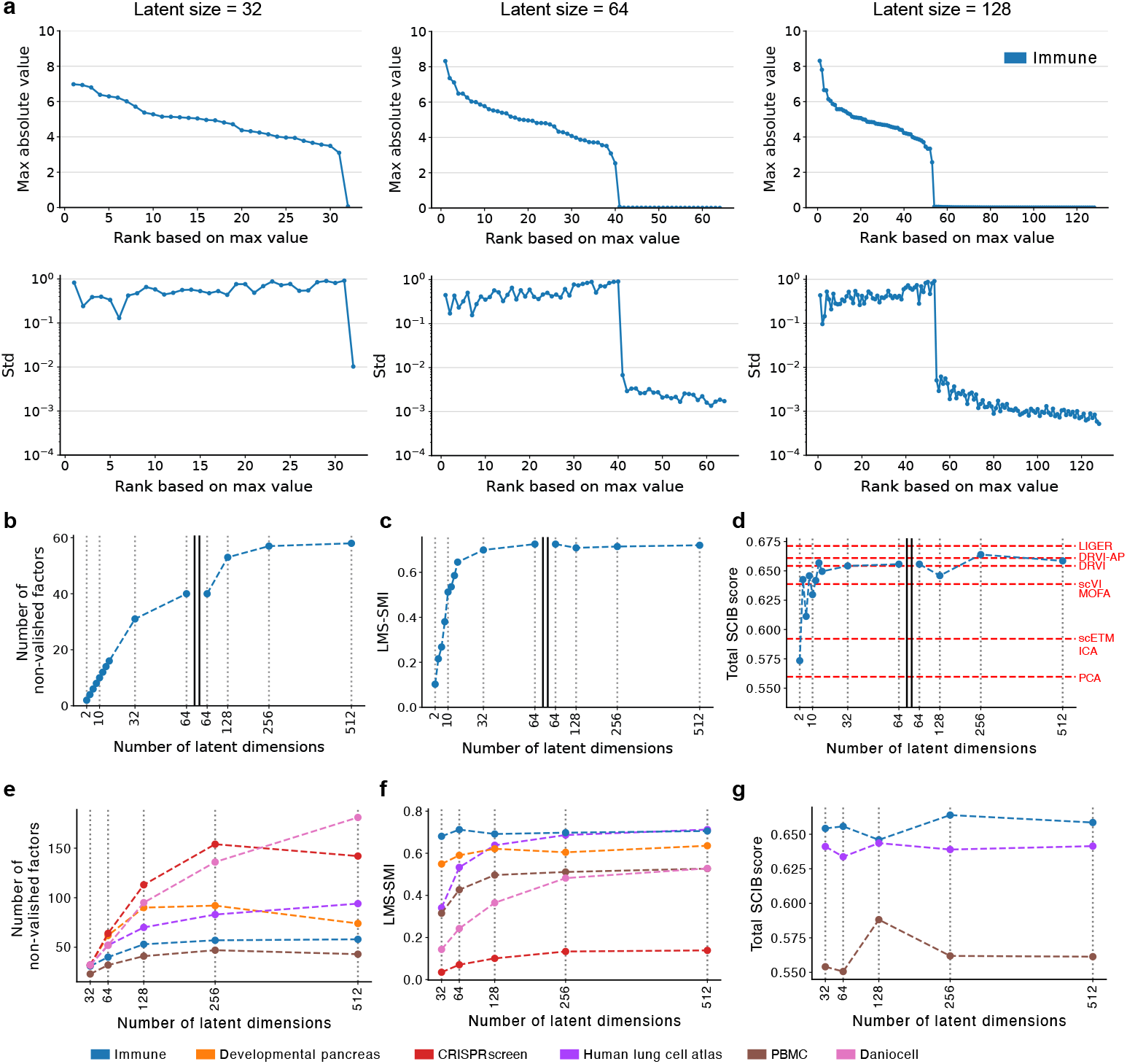
The saturation of the number of employed latent factors by DRVI. **a**, The vanishing dimensions phenomenon. DRVI is trained with 32, 64, and 128 latent factors on the immune dataset. Each point is a latent factor. The top row indicates max absolute values, and the bottom row indicates log-scale standard deviations of latent factors. latent factors are ranked based on max absolute value. Each column indicates a run with a different number of latent dimensions. **b-d**, Saturation of the number of non-vanished factors (b), disentanglement score (c), and integration score (d) when more latent factors are provided to DRVI trained on the immune dataset. Red lines annotate integration scores for benchmarked methods with default parameters. The x-axes on the right-hand side of the plots are shrunk by a factor of 5. **e-g**, Saturation of the number of non-vanished factors (e), disentanglement score (f), and integration score (f) when more latent factors are provided to DRVI trained on the benchmarking datasets. LMS, latent matching score; SMI, scaled mutual information.

**Supplemental Figure 5:**
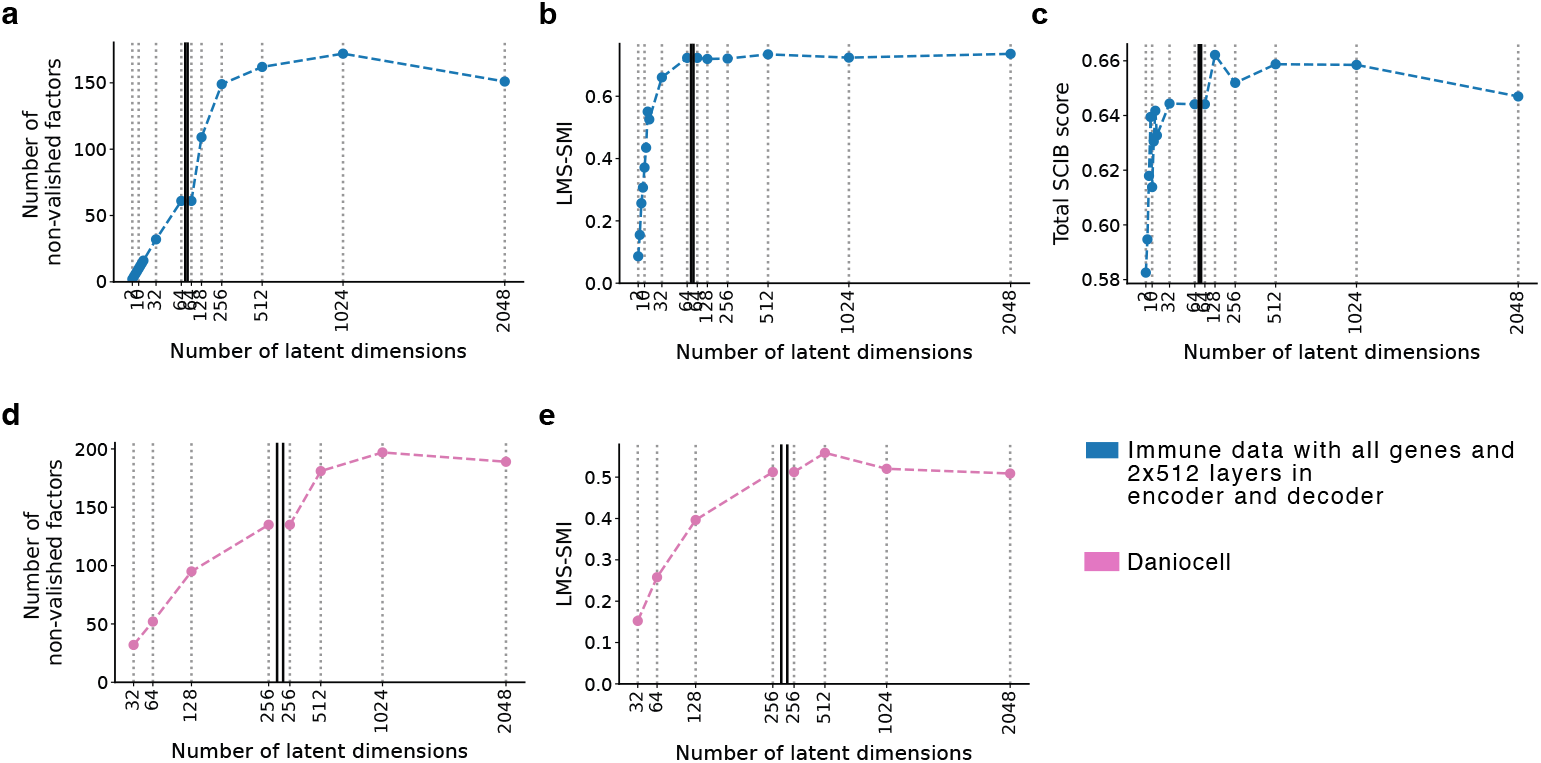
Further analysis on the saturation of employed latent factor. **a-c**, DVRI’s behavior trained on all genes of the immune dataset with different number of latent dimensions. The number of non-vanished dimensions (a), Disentanglement quality (b), and integration performance (c) are plotted when we change the number of latent dimensions. DRVI is trained on the full gene set of the immune dataset. Encoder and decoders contain 2 layers of 512 hidden neurons. As we observe, DRVI saturates at some specific point. At this point, disentanglement performance also saturates. Integration performance saturates and drops a bit. **d**,**e**, DVRI’s behavior trained on Daniocell data beyond 512 latent dimensions. The number of non-vanished dimensions (d) and disentanglement quality (e) are plotted when we change the number of latent dimensions. As we observe, even in Daniocell data where there are 156 cell types, DRVI saturates at some point. At this point, disentanglement performance also saturates. LMS, latent matching score; SMI, scaled mutual information.

**Supplemental Figure 6:**
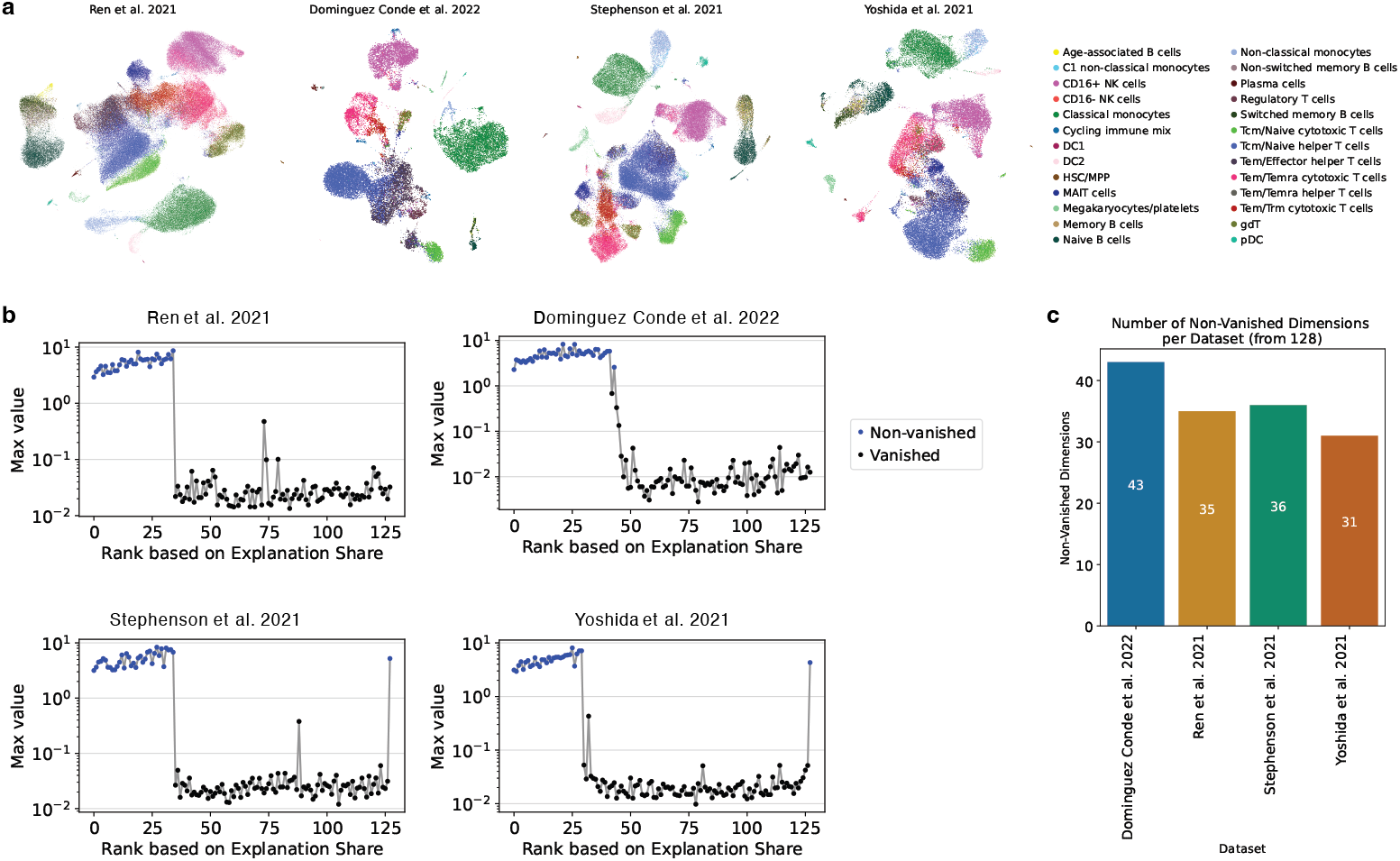
Training DRVI on four datasets of the Blood data. **a**, UMAP visualizations of individual Blood datasets, integrated by separate DRVI models. Cells are colored according to their curated annotations. **b**, Visualization of the vanished factors for each of the four DRVI models trained on the Blood datasets. **c**, The total number of non-vanished factors for each model trained on an individual Blood dataset.

**Supplemental Figure 7:**
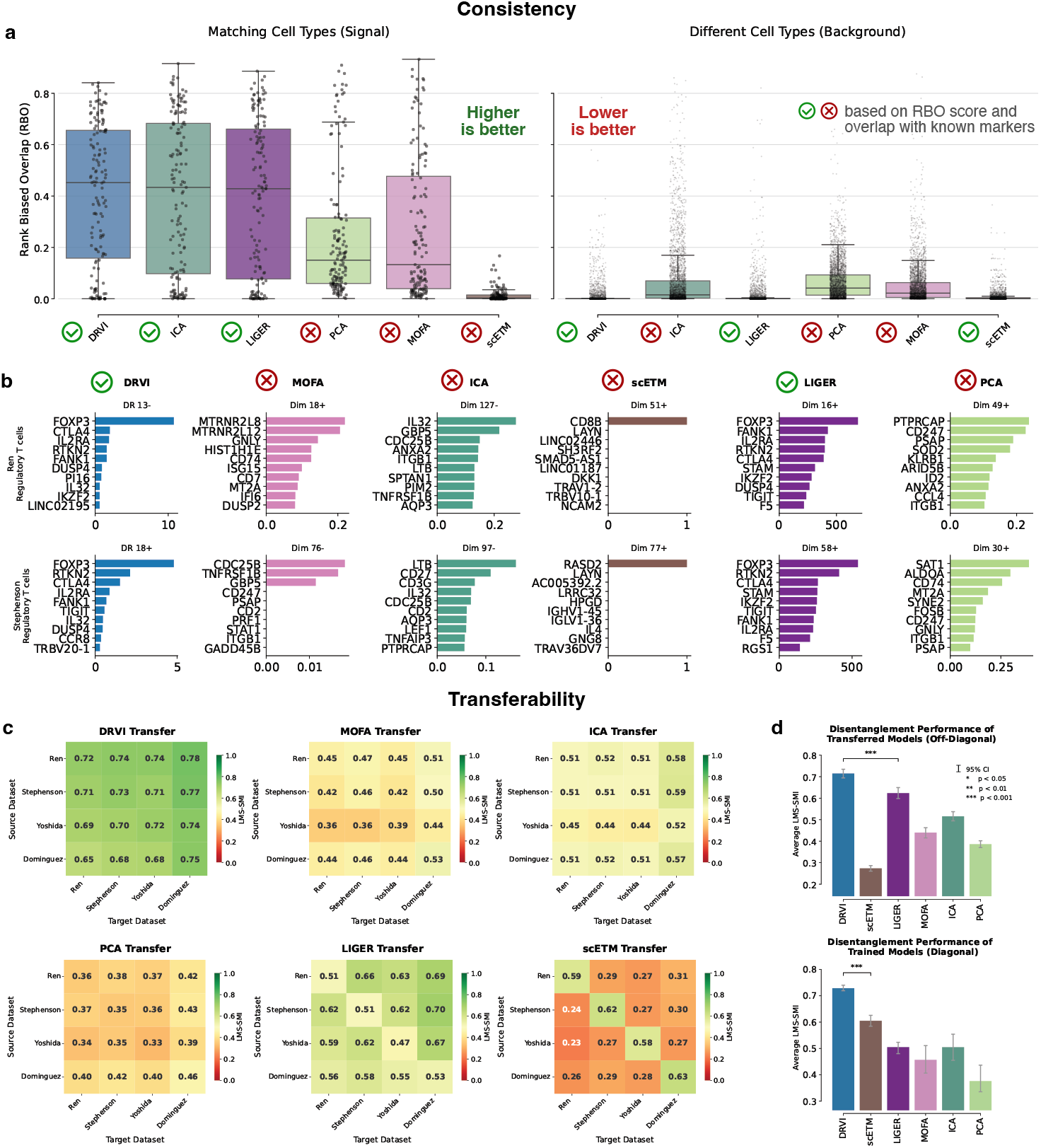
DRVI learned factors are consistent and transferable to unseen datasets. DRVI and the interpretable baselines are trained separately on four datasets of the Blood data. **a**, RBO similarity between ordered gene signatures of factors matched to the same cell type across datasets (signal; higher is better) or to different cell types (background; lower is better). Checks and crosses indicate whether the corresponding factors are consistent across datasets based on RBO. **b**, Example gene signatures in two datasets for factors best matched to regulatory T cells. Checks and crosses indicate whether the top genes overlap with known markers. **c**, Disentanglement matrices for benchmarked methods trained on one Blood dataset (source) and evaluated on another (target). Each trained model is applied to the remaining three datasets. Entries show LMS-SMI disentanglement scores. **d**, Average LMS-SMI scores for off-diagonal transfer settings and diagonal within-dataset evaluations. Error bars indicate 95% confidence intervals. LMS, latent matching score; RBO, Rank-Biased Overlap; SMI, scaled mutual information.

**Supplemental Figure 8:**
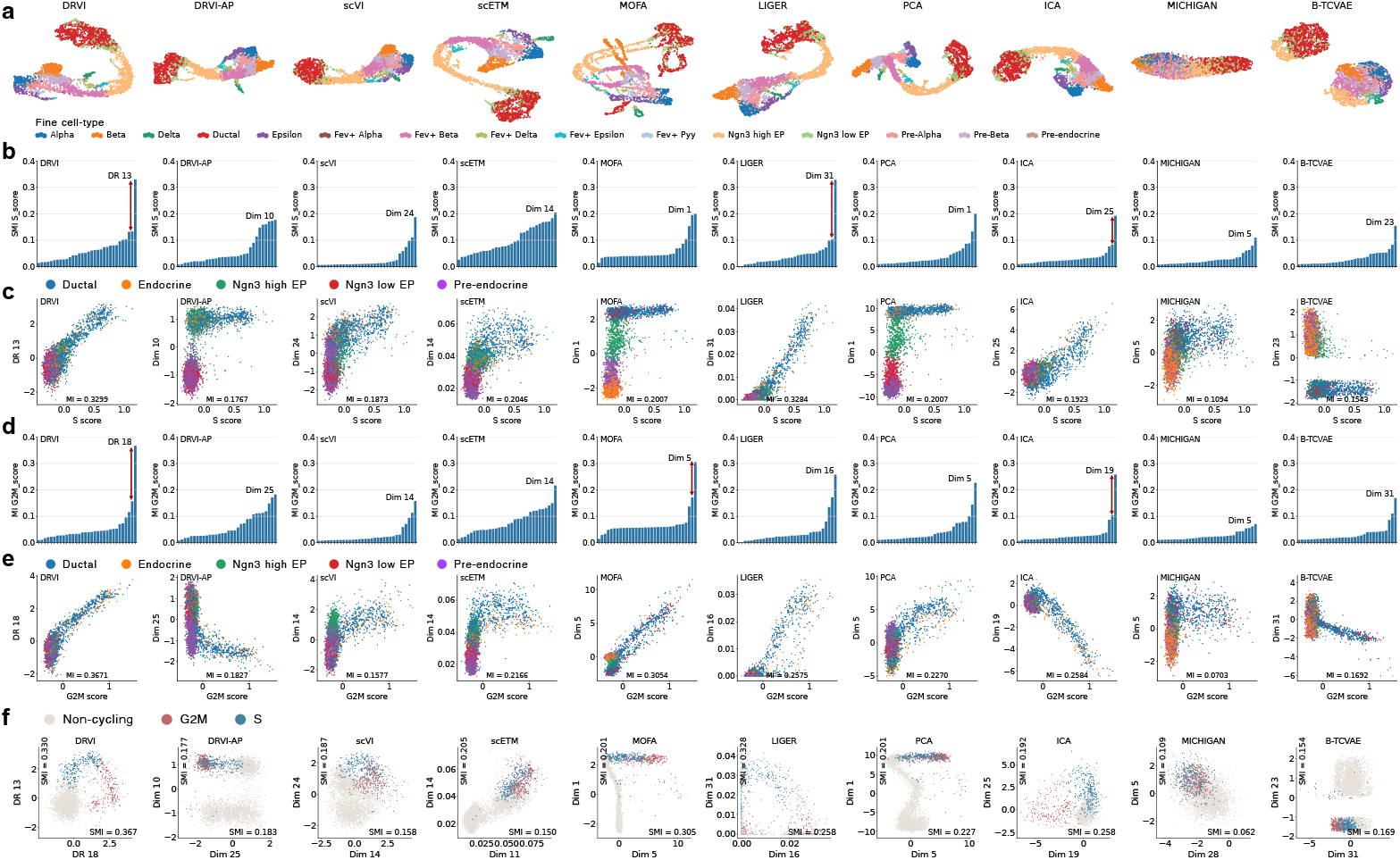
DRVI disentangles cell cycle stages. **a**, UMAPs of the developmental pancreas data applied to the latent spaces generated by benchmarked methods. Cells are colored by finest-level annotations. **b-e**, Barplots indicating Mutual information of each latent factor with cell-cycle S-score (b) and G2M-score (d). The factor maximizing the mutual information with the S-score(c) and G2M-score(d) are plotted versus the S-score (c) and G2M-score (e) for each method. Cells are colored by coarse annotations.**f**, The learned cell-cycle manifolds. The scatterplots of factors maximizing the mutual information with respect to the S-score and G2M-score are plotted. Cells are colored by the dominant cell cycle phase. MI, mutual information.

**Supplemental Figure 9:**
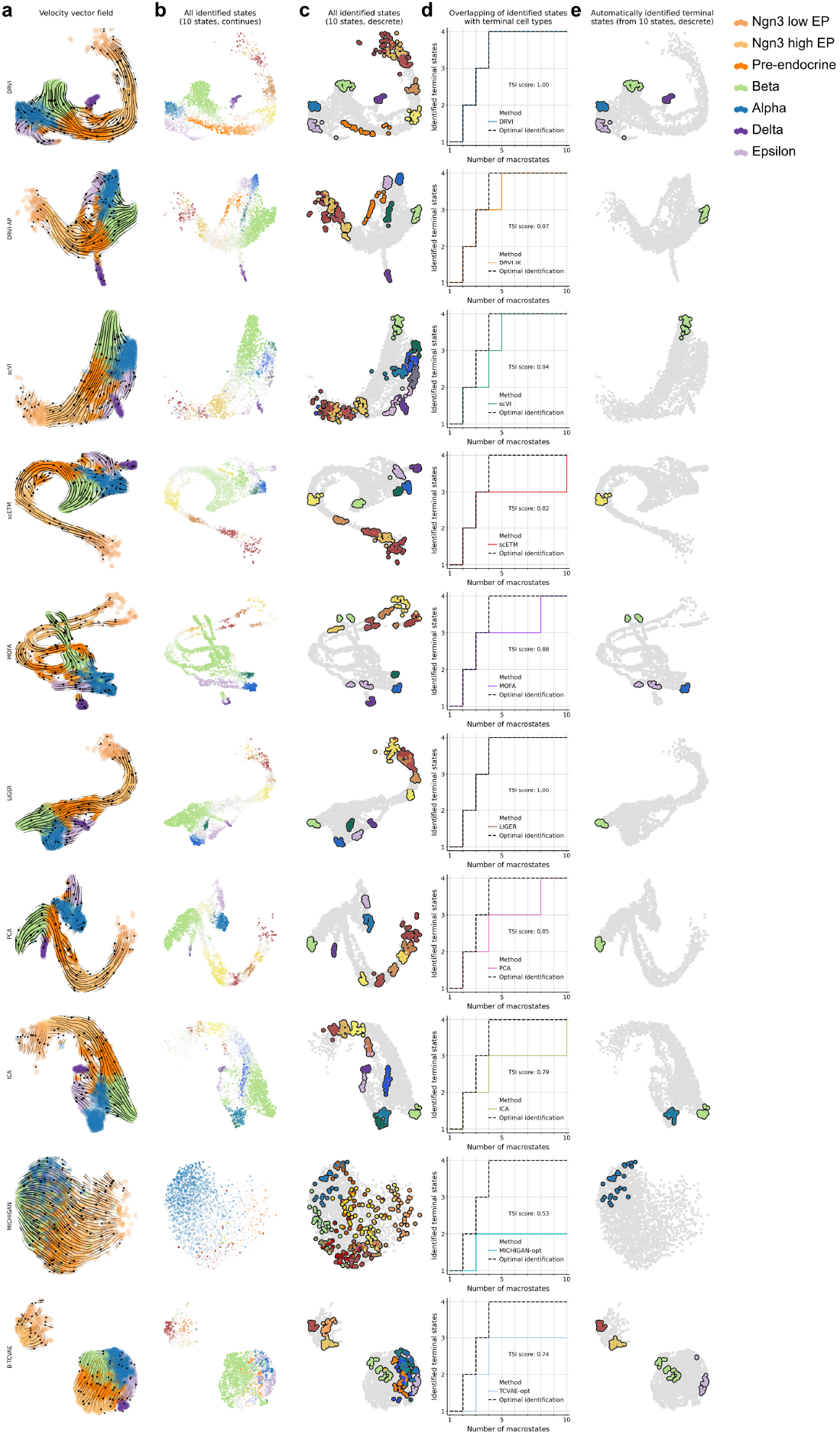
Developmental fate prediction based on model embeddings. **a**, UMAP visualizations of scVelo RNA velocity vector fields, generated based on embeddings derived from various benchmarked models. **b**,**c**, UMAP visualizations of stationary states identified by CellRank, based on the velocity vector fields shown in (a). **d**, Terminal State Identification (TSI) scores for different benchmarked methods. This metric quantifies the overlap between identified stationary states and ground-truth terminal states as the number of macrostates increases. **e**, UMAP visualization of automatically identified terminal states from ten macrostates using CellRank. TSI: Terminal State Identification.

**Supplemental Figure 10:**
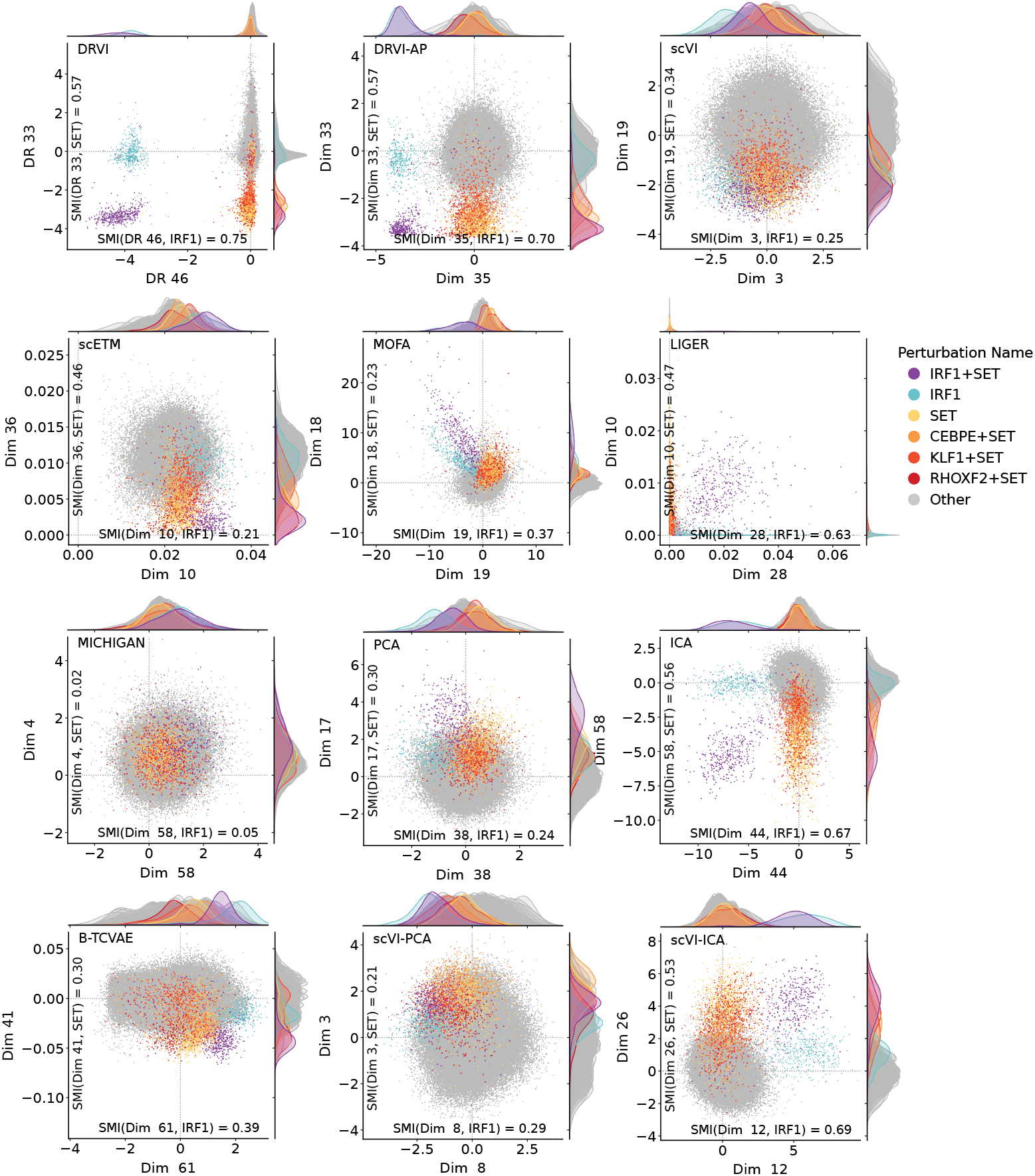
DRVI identifies and preserves the geometry of combinatorial perturbations. Each scatterplot illustrates the activity of the top two factors having the highest mutual information with respect to IRF1 and SET perturbations for a benchmarked method. The cells perturbed by IRF1 and SET and all combinatorial perturbations containing them are highlighted with distinct colors. Any other perturbation signature is colored grey. SMI, scaled mutual information.

**Supplemental Figure 11:**
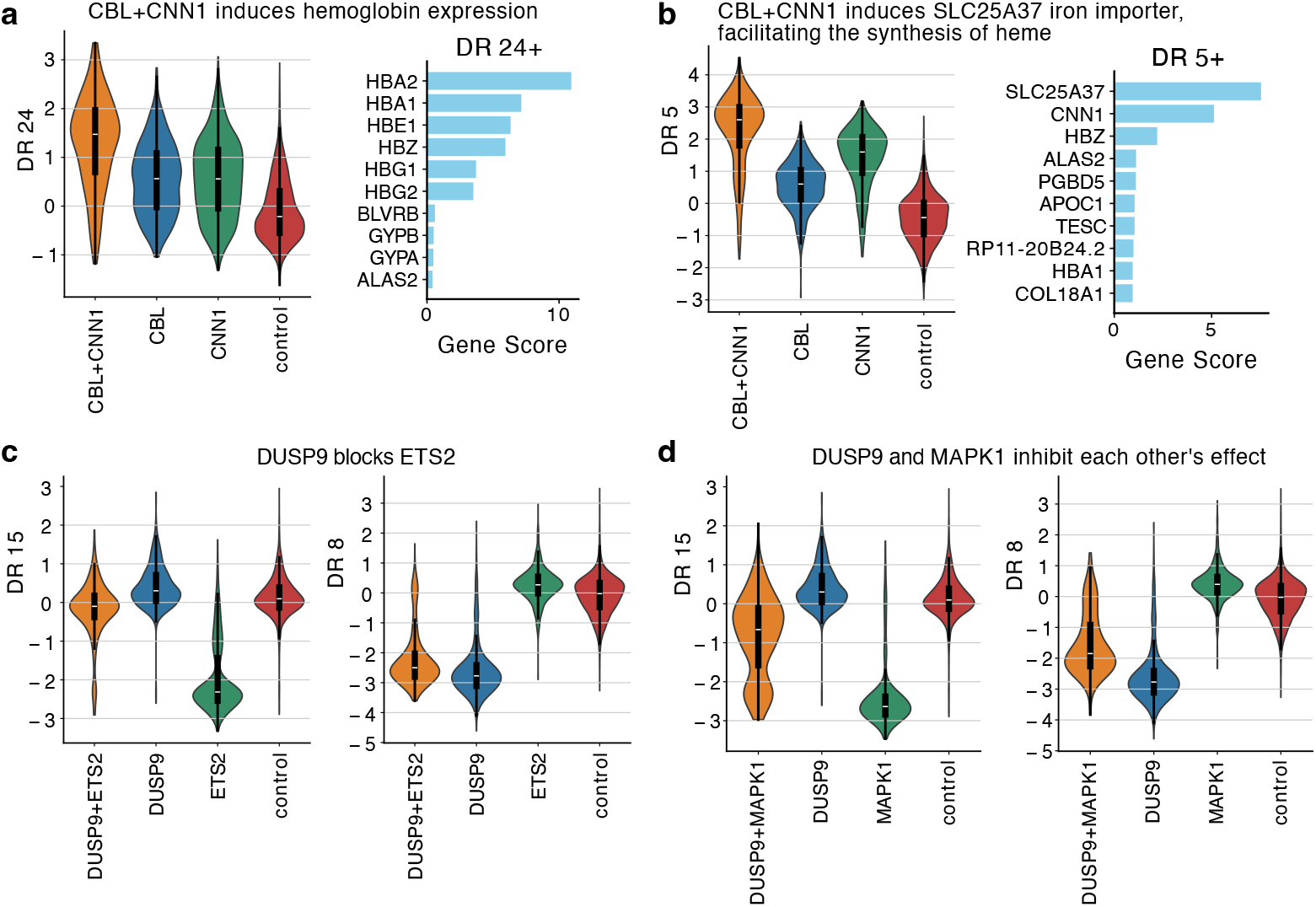
DRVI’s latent space effectively models interacting double perturbations. **a**, When the **CBL+CNN1** gene combination is perturbed, there’s a significant increase in the expression of **hemoglobin** (DR 24+). **b**, Perturbing **CBL+CNN1** leads to the overexpression of the iron importer **SLC25A37** (DR 5+), which supports the role of CBL+CNN1 in regulating hemoglobin expression. **c, DUSP9** acts as an inhibitor of **ETS2**, as its effect (DR 15-) is neutralized when DUSP9 is also perturbed. **d, DUSP9** and **MAPK1** exhibit a mutual inhibitory relationship, which is evident from the decreased response signatures of both genes (DR 15- and DR 8-) when they are simultaneously perturbed.

**Supplemental Figure 12:**
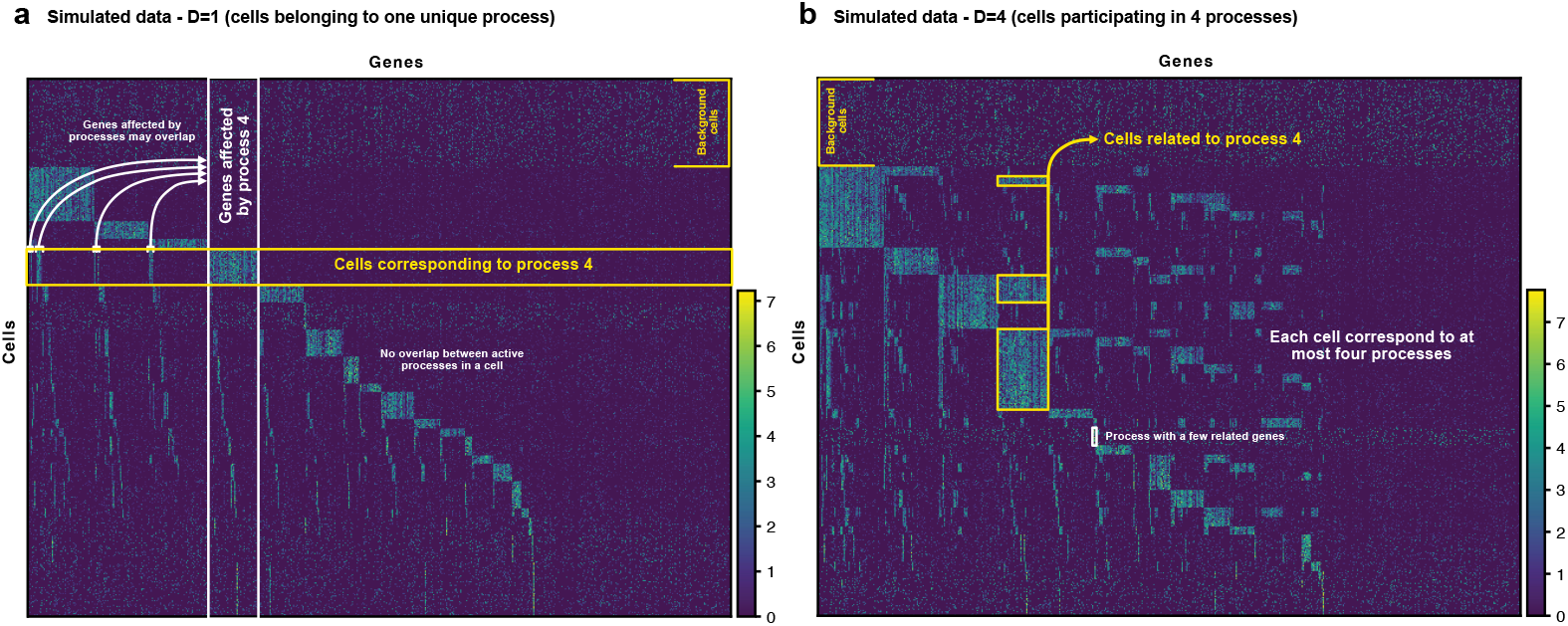
Heatmap of simulated datasets with known ground truth. The first simulated dataset (**a**) assigns non-background cells to a single process, while genes can be influenced by multiple processes. The second simulated dataset (**b**) allows cells to participate in up to four processes, introducing controlled overlap and compositional complexity. Cells with similar ground truth are grouped, as are genes affected by the same programs. Both cell and gene groups are then arranged to maximize their diagonal presentation.

**Supplemental Figure 13:**
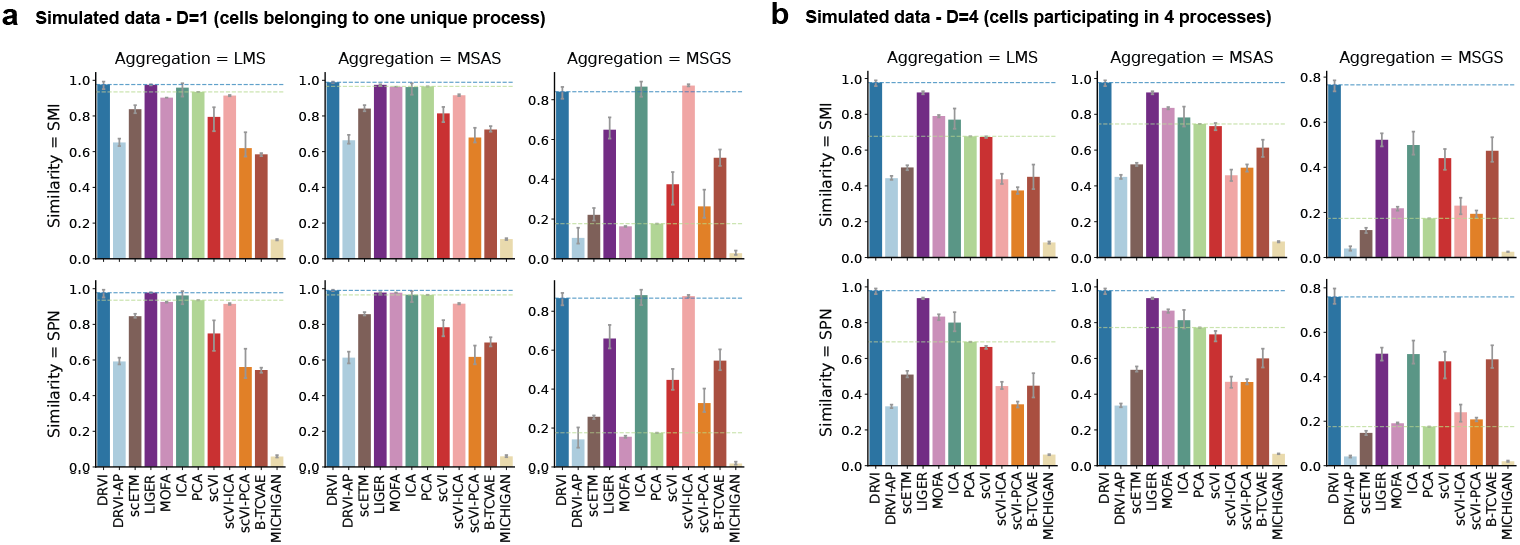
Evaluation results for simulated datasets with known ground truth. **a**,**b**, We calculated metric values for each benchmarked model across the simulated datasets. These metrics are subsequently presented for all possible combinations formed by the SMI and SPN similarity functions and the LMS, MSAS, and MSGS aggregation functions. In cases where cellular activities are mutually exclusive (a), DRVI, ICA, MOFA, PCA, LIGER, and scVI-ICA all perform effectively. However, when processes overlap (b), only DRVI and LIGER are capable of accurately identifying ground-truth processes. LMS, latent matching score; MSAS, Most Similar Averaging Score; MSGS, Most Similar Gap Score; SMI, scaled mutual information; SPN, same-process neighbors.

**Supplemental Figure 14:**
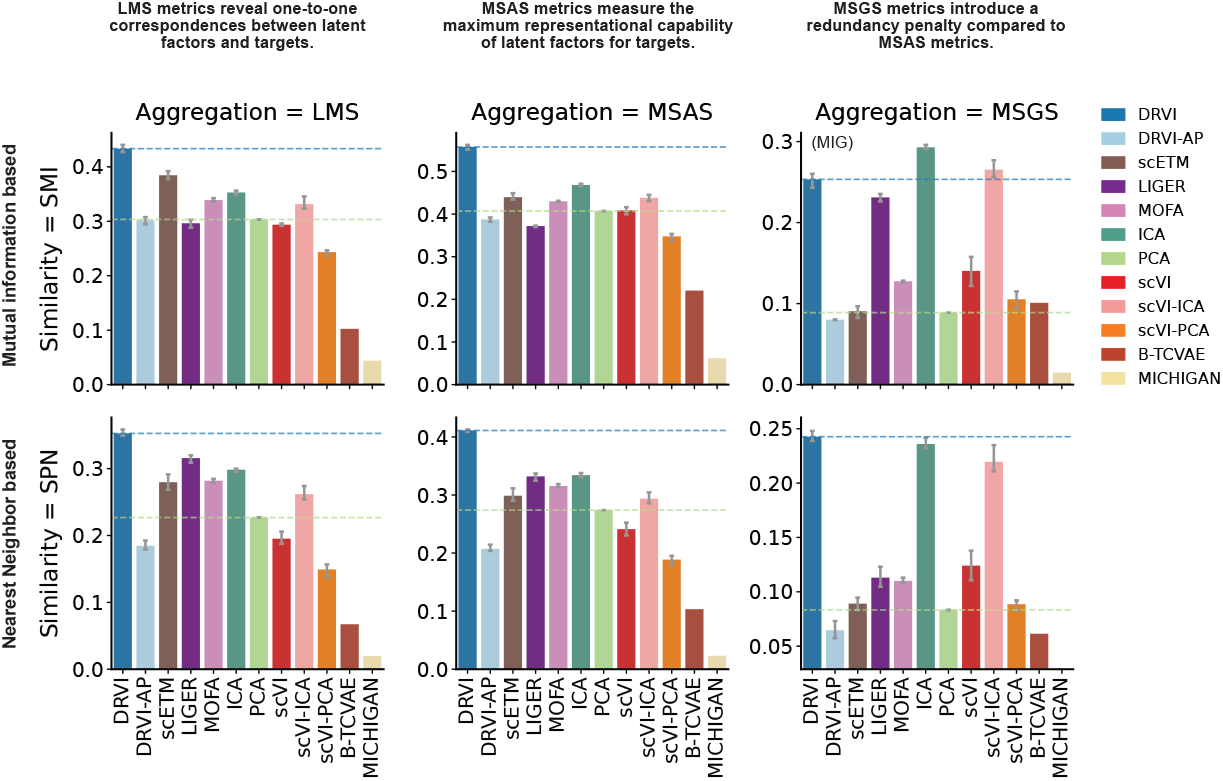
Summary of all metrics on the six primary benchmark datasets. This plot displays the average performance of the benchmarked methods across the six primary datasets. Means are shown for every combination of the SMI and SPN similarity functions with the LMS, MSAS, and MSGS aggregation functions. LMS, latent matching score; MSAS, Most Similar Averaging Score; MSGS, Most Similar Gap Score; SMI, scaled mutual information; SPN, sameprocess neighbors.

**Supplemental Figure 15:**
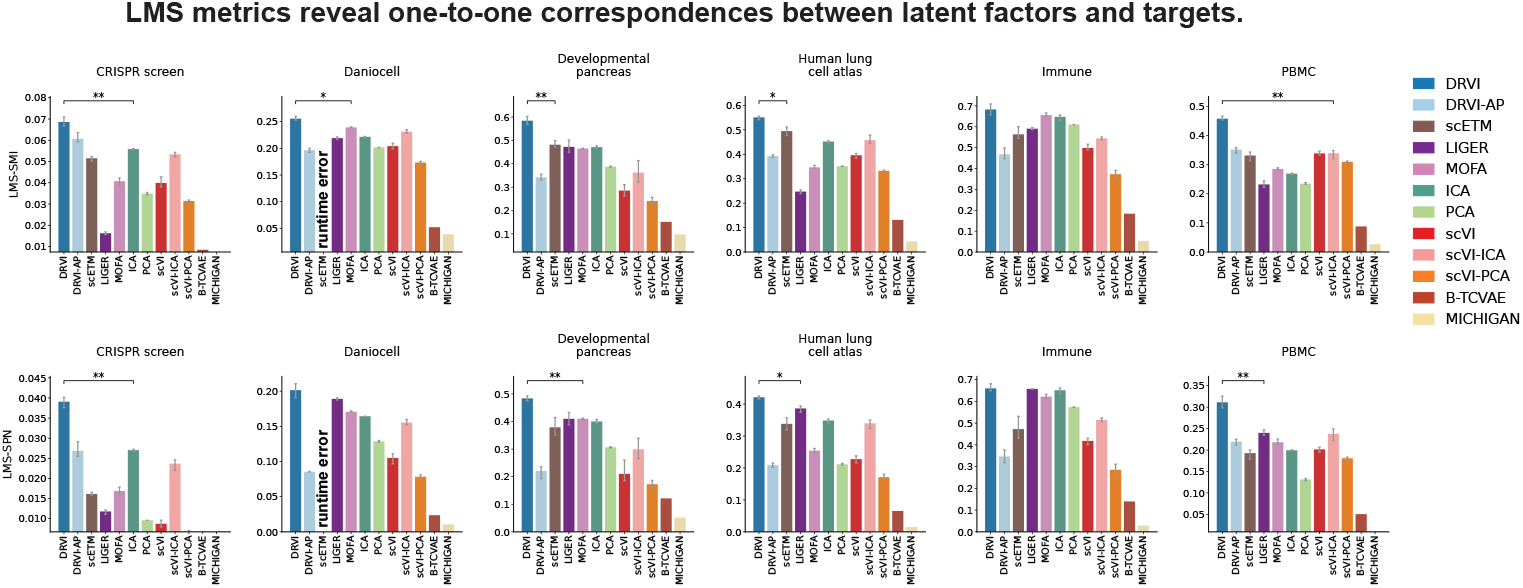
Results of the primary disentanglement metric family on the six primary benchmark datasets. The results of the LMS metrics, as the primary metric family, are shown for each primary dataset. LMS measures the ability of the model to capture biological processes in individual latent factors by matching latent factors and proxy variables. Asterisks indicate statistical significance of DRVI versus the next-best method per dataset. LMS, latent matching score; SMI, scaled mutual information; SPN, same-process neighbors.

**Supplemental Figure 16:**
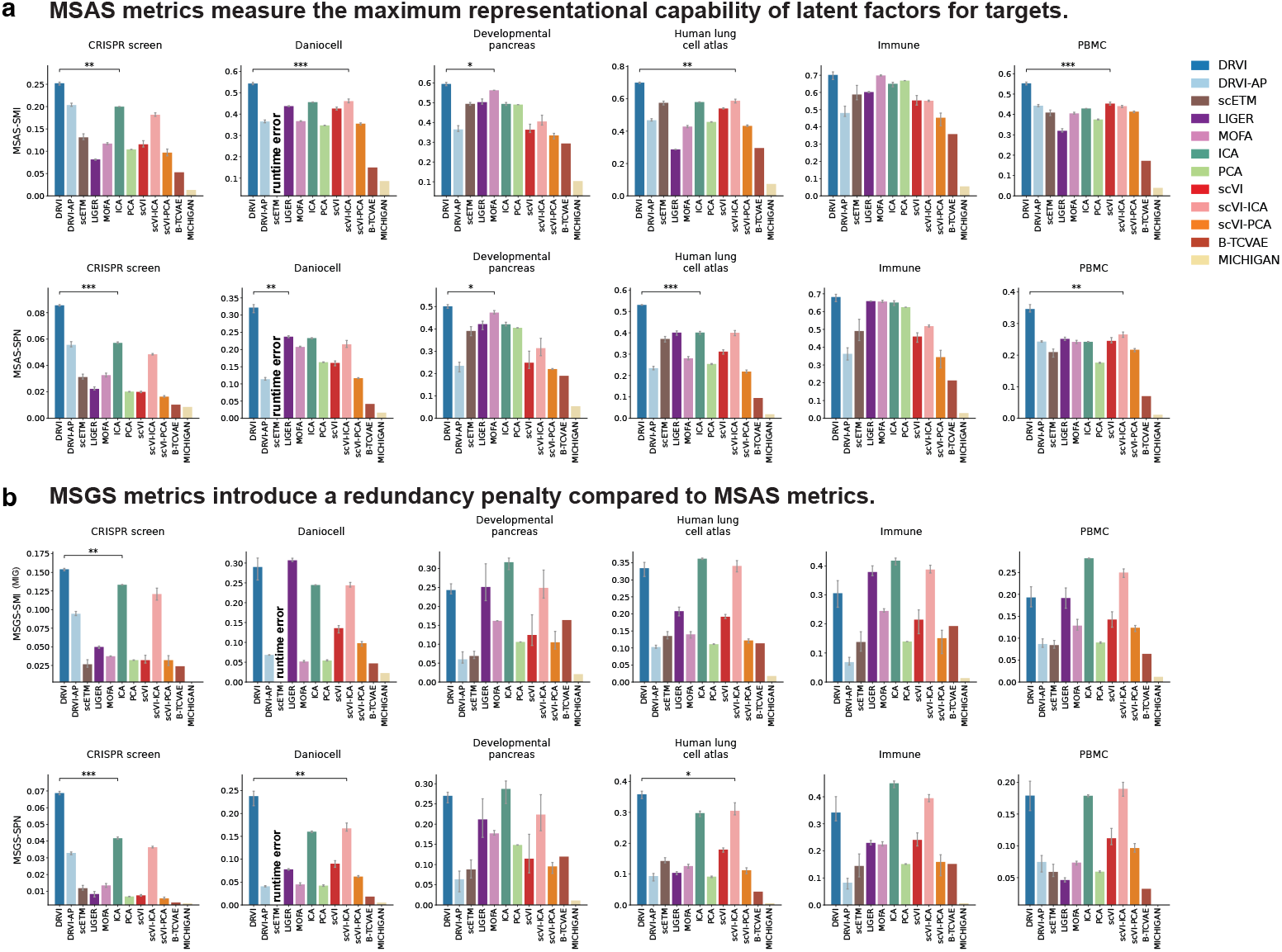
Results of the secondary disentanglement metrics on the six primary benchmark datasets. **a**,**b**, The results of the MSAS (a) and MSGS (b) as secondary metrics over the six primary datasets are provided. MSAS shows the capability of the model to capture biological processes in individual latent factors regardless of the redundancy. MSGS measures the amount of captured biological information that is specific to a single (probably overlapping) factor by penalizing redundancy. Unlike LMS, as the main metric, neither MSAS nor MSGS penalizes overlapping processes in a single factor. LMS, latent matching score; MSAS, Most Similar Averaging Score; MSGS, Most Similar Gap Score; SMI, scaled mutual information; SPN, same-process neighbors; MIG, mutual information gap.

**Supplemental Figure 17:**
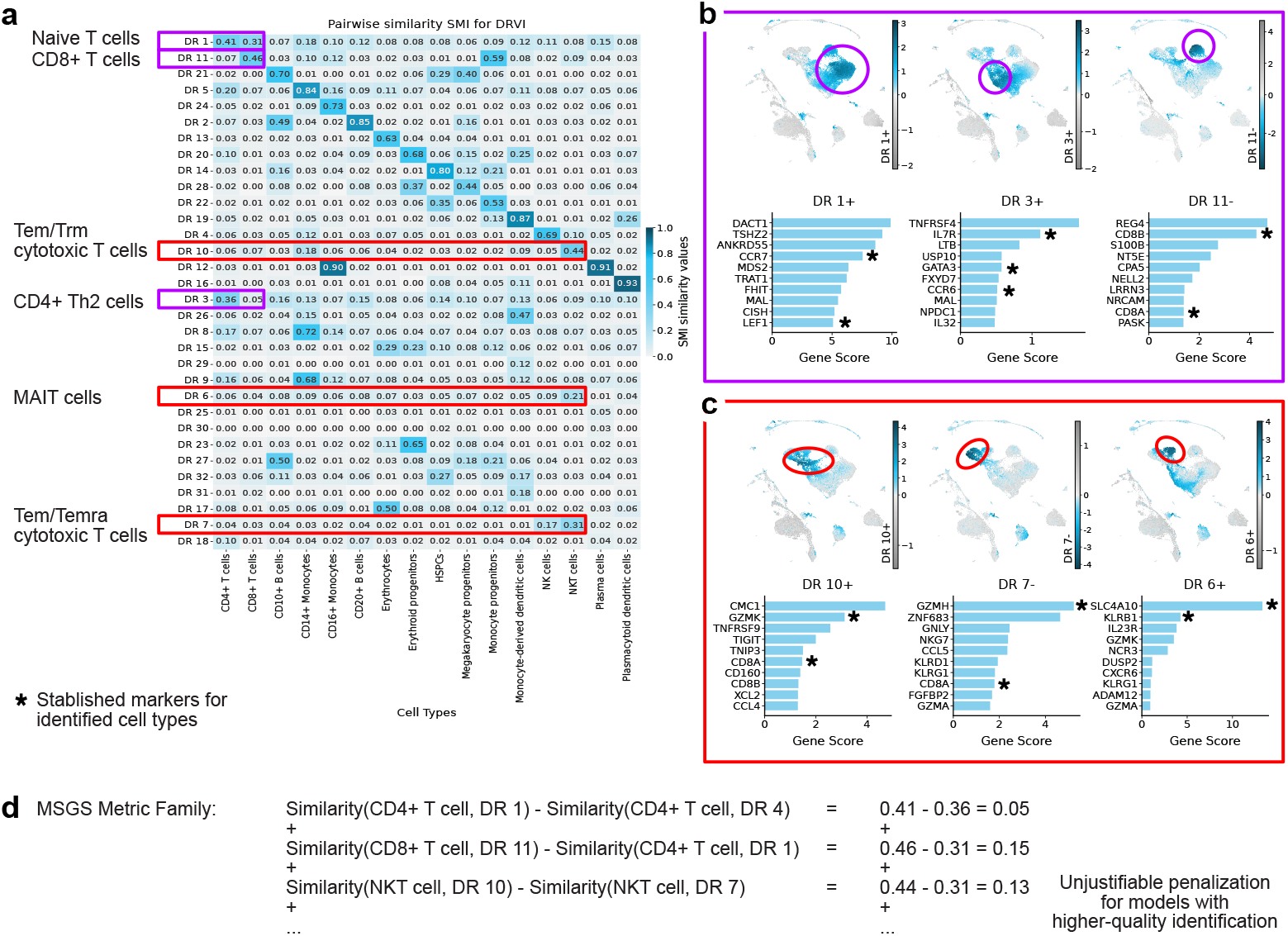
DRVI identifies granular immune cell subtypes that are penalized by MSGS metrics. **a**, Pairwise Similarity Heatmap. This heatmap compares the similarity of DRVI latent factors against broad, “ground truth” cell type annotations. Highlighted rows show factors of DRVI capturing more granular subtypes. **b, c**, Granular Subtype Identification. These panels show the specific gene markers for DRVI’s clusters, confirming they represent distinct, biologically relevant cell states that were grouped together in the original annotations. Panel (b) shows that DRVI correctly separates Naive T cells (DR 1), marked by CCR7 (a classic marker for naive T cells) and LEF1 (a transcription factor that is highly expressed in naive T cells) from CD4+ Th2 cells (DR 4), marked by GATA3 (transcription factor for Th2 cells) and IL7R (a cytokine critical for the survival of memory T cells). The ground truth annotation simply labels both as CD4+ T cells.. Panel (c) shows that DRVI also distinguishes between highly specific cytotoxic and innate-like T cells. It identifies Tem/Trm cytotoxic T cells (GZMK+ inflammatory CD8+ T cells, in DR 10), Tem/Temra cytotoxic T cells marked by ZNF683 (Tissue-Resident Memory CD8+ T cells, in DR 8), and MAIT cells marked by SLC4A10 and KLRB1 (DR 7). The ground truth annotations grouped these distinct types under a broad label of NKT cells. **d** problem of MSGS Metrics when ground-truth labels are imperfect. This metric family unjustifiably penalizes the DRVI model for its superior performance. By subtracting the similarity scores of valid subtypes (e.g., Similarity(CD4+ T cell, DR 1) - Similarity(CD4+ T cell, DR 4)), the metric punishes the model precisely because it was able to distinguish between Naive T cells and Th2 cells — a higher-quality identification that was more accurate than the provided ground truth.

**Supplemental Figure 18:**
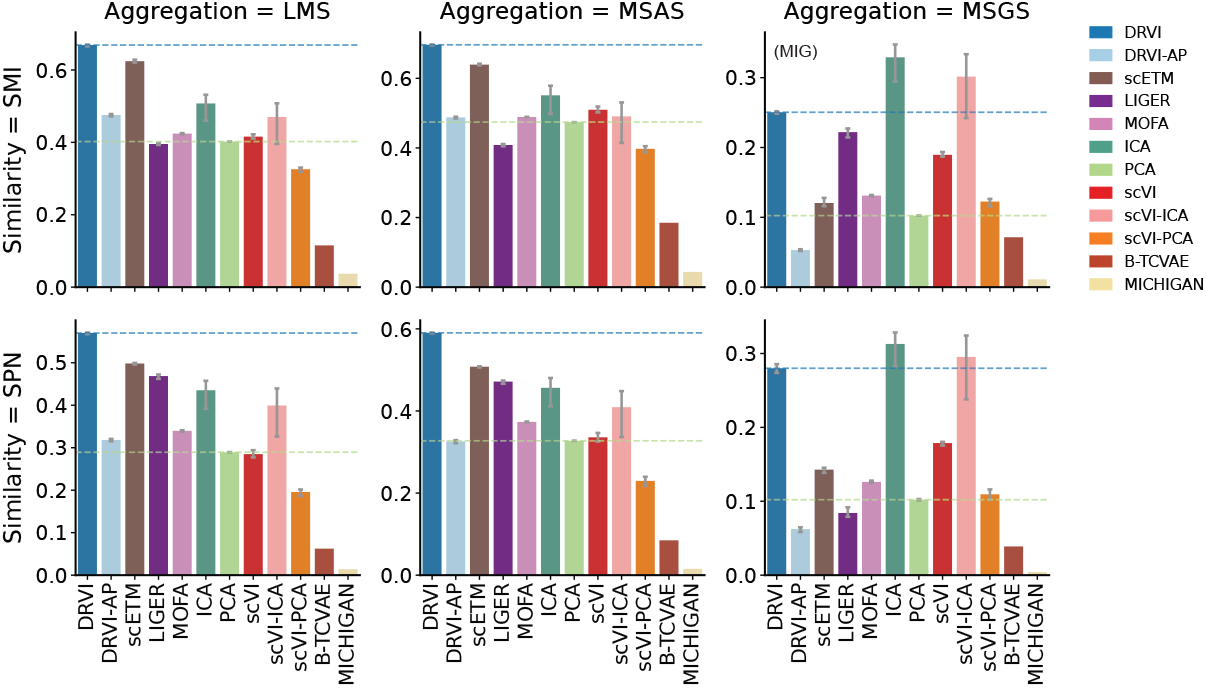
Summary of all metrics on the twelve CellHint organ datasets. This plot displays the average performance of the benchmarked methods across the twelve CellHint multi-batch datasets. Means are shown for every combination of the SMI and SPN similarity functions with the LMS, MSAS, and MSGS aggregation functions. LMS, latent matching score; MSAS, Most Similar Averaging Score; MSGS, Most Similar Gap Score; SMI, scaled mutual information; SPN, same-process neighbors.

**Supplemental Figure 19:**
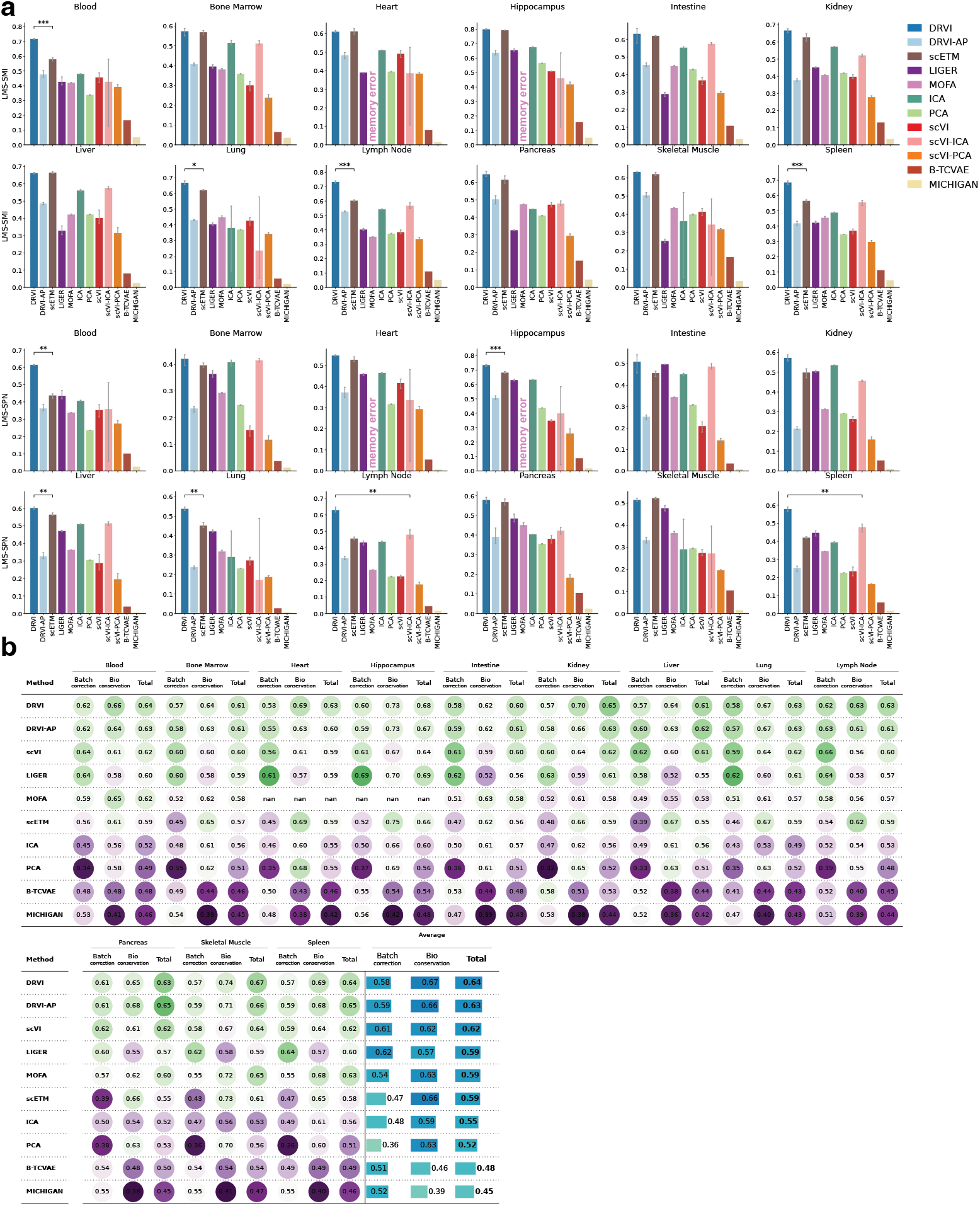
Primary disentanglement and integration results on the twelve CellHint organ datasets. **a**, LMS-SMI and LMS-SPN disentanglement metrics for each organ dataset. **b**, scIB batch correction, biological conservation, and total integration scores for each organ dataset. Asterisks indicate statistical significance of DRVI versus the next-best method per dataset. LMS, latent matching score; SMI, scaled mutual information; SPN, same-process neighbors.

**Supplemental Figure 20:**
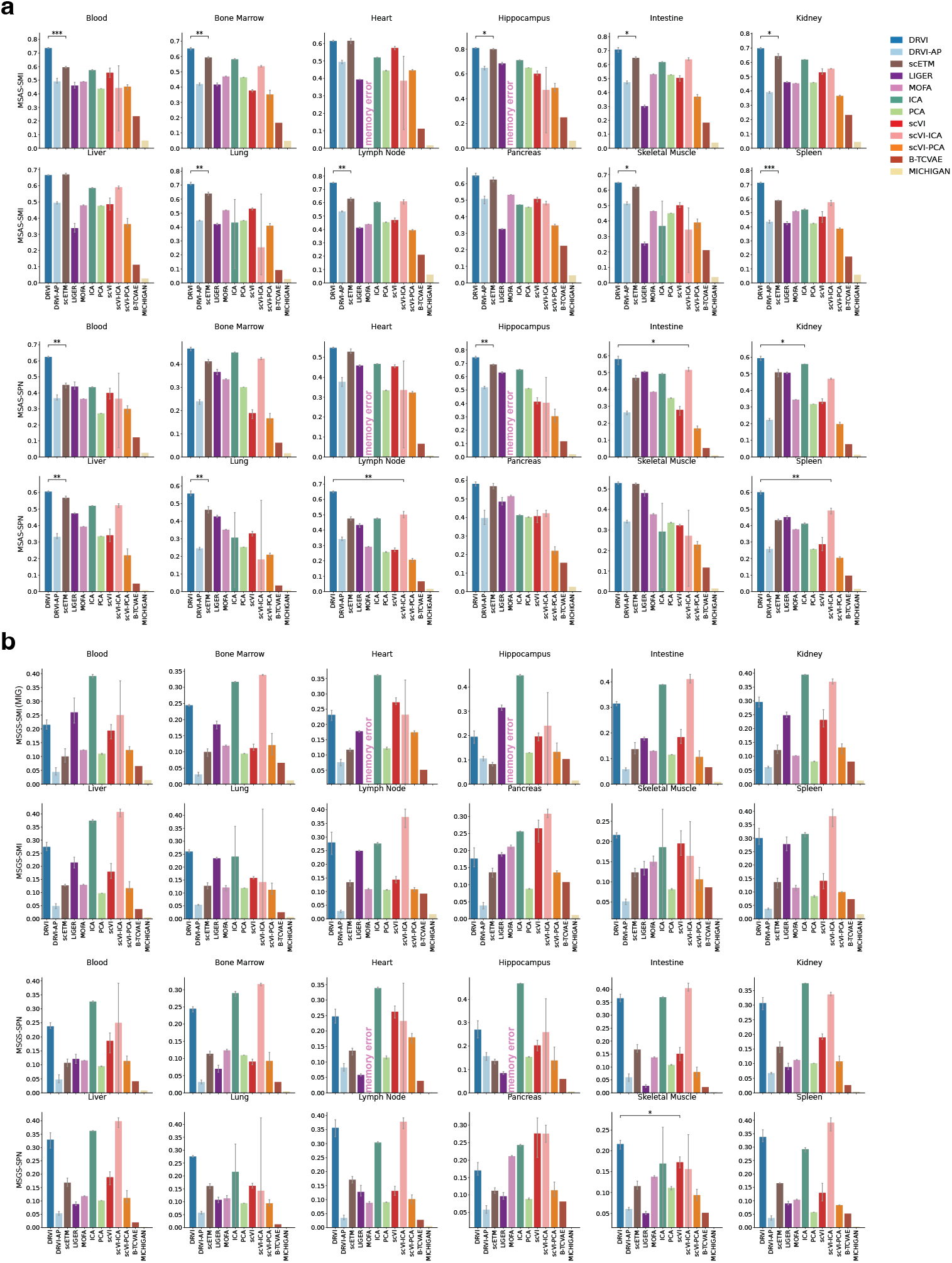
Results of the secondary disentanglement metrics on the twelve CellHint organ datasets. **a**,**b**, The results of the MSAS (a) and MSGS (b) as secondary metrics over the twelve CellHint organ datasets are provided. LMS, latent matching score; MSAS, Most Similar Averaging Score; MSGS, Most Similar Gap Score; SMI, scaled mutual information; SPN, same-process neighbors; MIG, mutual information gap.

**Supplemental Figure 21:**
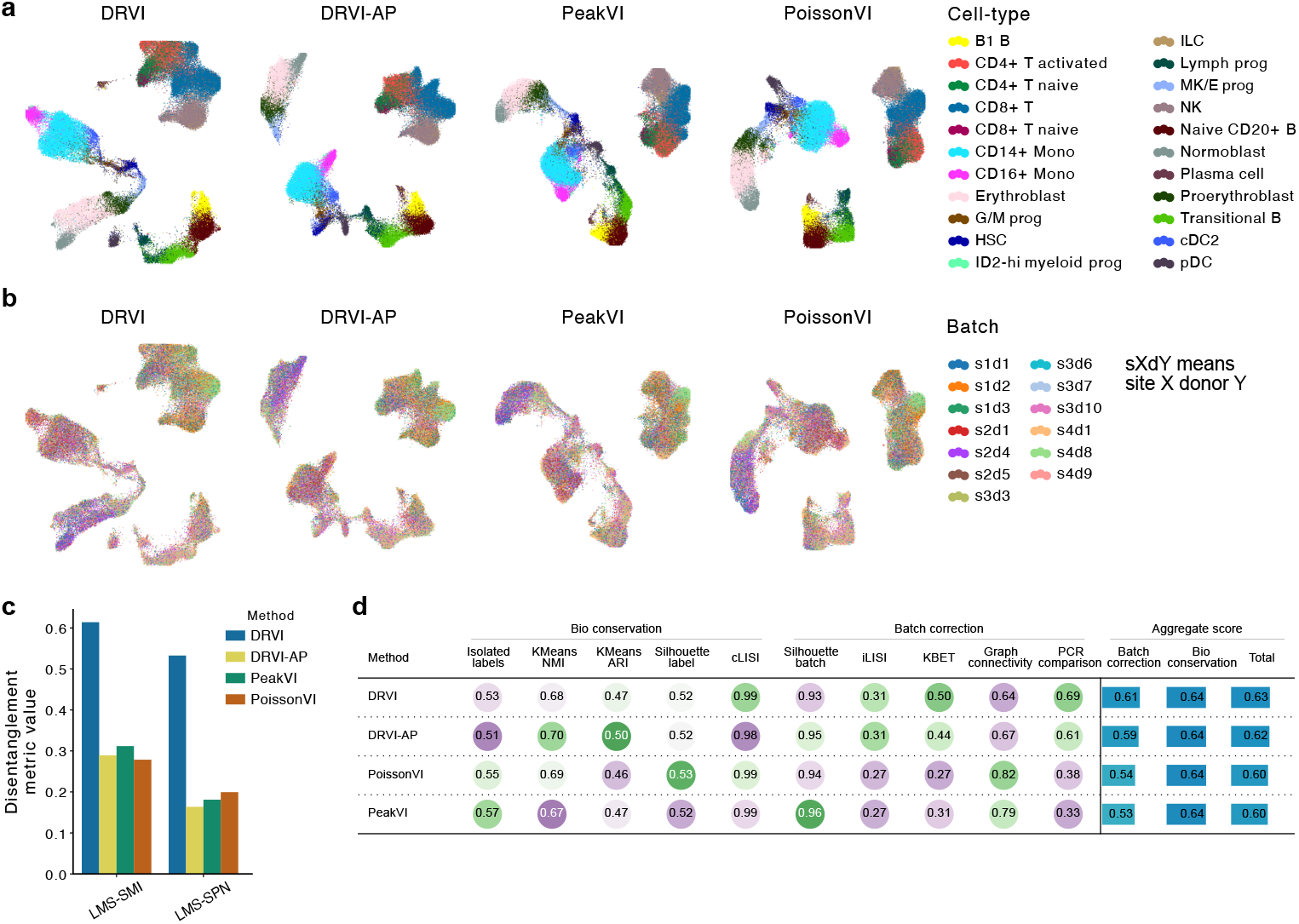
DRVI is applicable to scATAC-seq data. **a**,**b**, UMAPs of the ATAC modality of the NeurIPS 2021 data integrated by DRVI, DRVI-AP, PeakVI, and PoissonVI colored by cell-type annotations (a) and batch identity (b). **c**, Disentanglement performance of the benchmarked methods for scATAC-seq data. DRVI performs significantly better in this task. **d**, Integration performance of the benchmarked methods based on the scIB metrics. DRVI performs better compared to the three other methods. LMS, latent matching score; SMI, scaled mutual information; SPN, same-process neighbors.

**Supplemental Figure 22:**
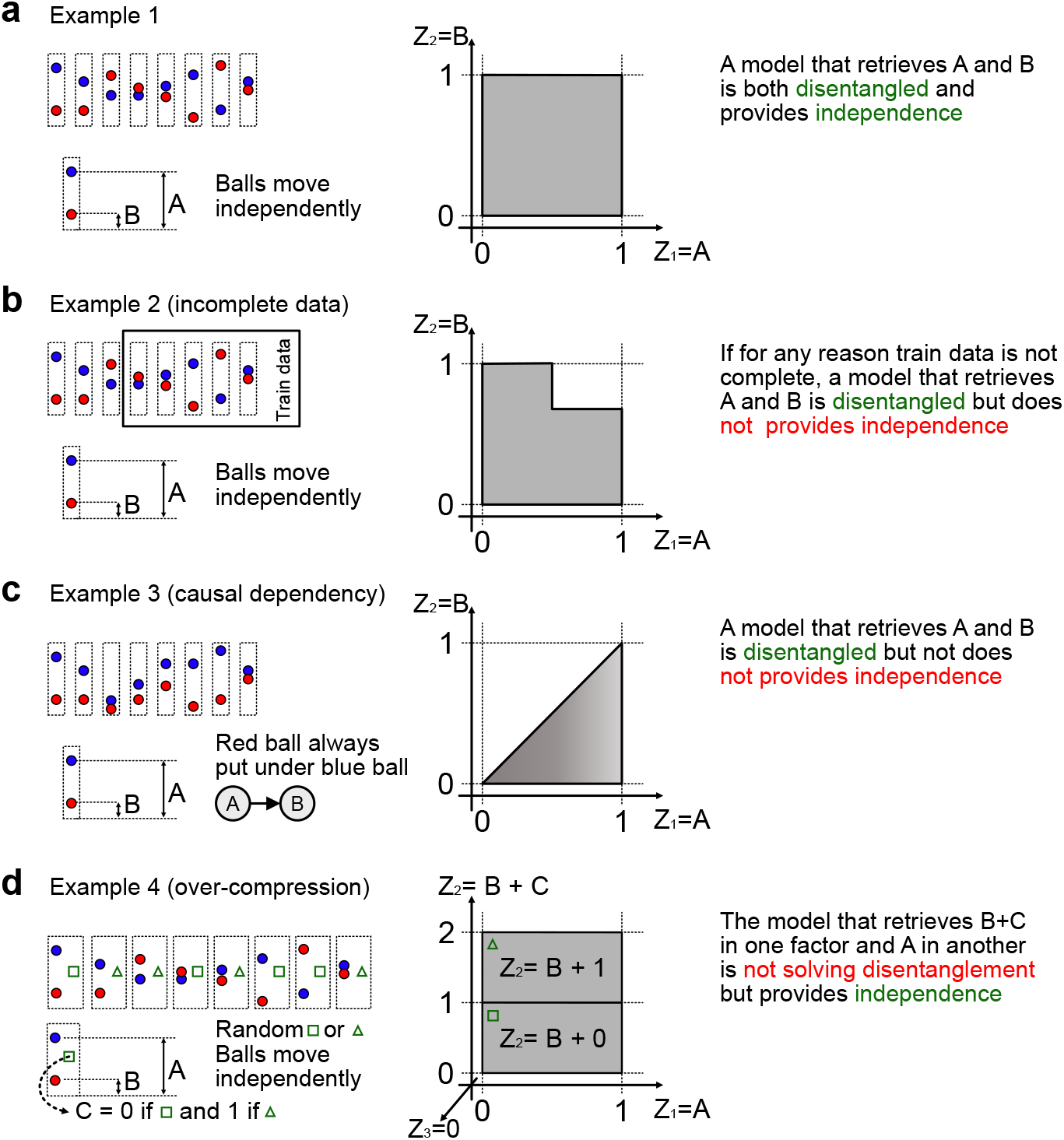
Disentanglement versus Statistical Independence. **a**, Alignment of Independence and Disentanglement. In the ideal scenario with independent underlying factors (e.g., *A* and *B*) and complete data coverage (square support), a disentangled model recovers factors that are also statistically independent. **b**, Disentanglement without Independence (Incomplete Data): When the training data is incomplete or biased (L-shaped support), correctly disentangled factors *A* and *B* will exhibit statistical dependence reflecting the observed distribution, despite being mechanically independent. **c**, Causal Dependency: A causal constraint, such as the red ball (*A*) always being positioned lower than the blue ball (*B*), restricts the data manifold (triangular support). A perfect disentanglement model recovers the distinct concepts *A* and *B* but yields correlated latent dimensions. **d**, Independence without Disentanglement: An over-compressed model that mixes factors (e.g., learning *A* and *B* +*C*) can achieve statistical independence in the latent space (rectangular support) but fails to disentangle the true underlying generative processes.

**Supplemental Figure 23:**
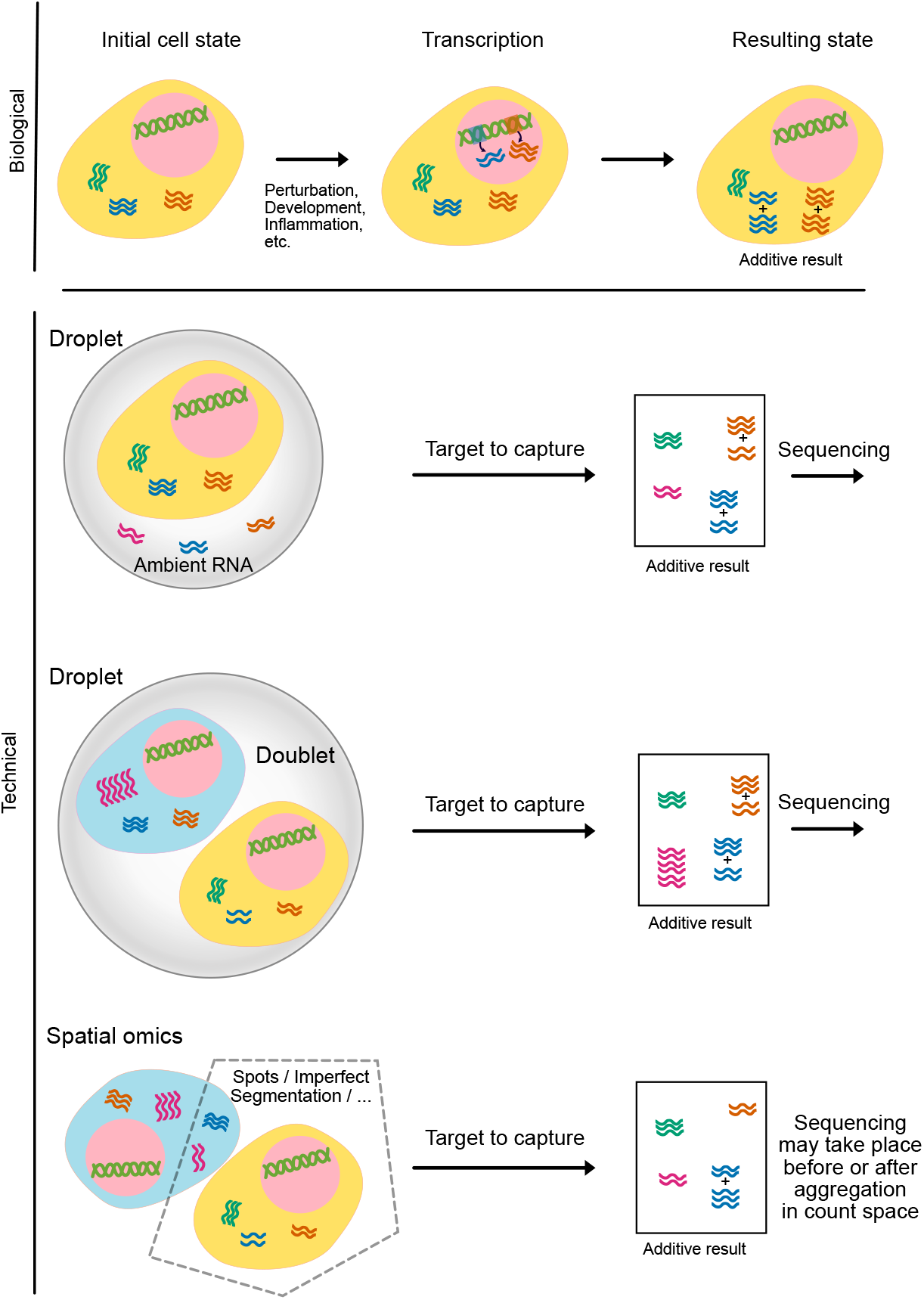
Plausibility of DRVI’s additive assumption in scRNA-seq data. The additivity of independent biological processes makes sense from both biological and technical perspectives. Newly transcribed genes (for example, during developmental bifurcations, induced perturbations, stress, etc.) add up within the cell and therefore add up in count space when sequenced. In addition, technical effects such as doublet cells, ambient background counts, and residual counts from neighboring cells due to imperfect segmentation in spatial omics show additive behaviors.

**Supplemental Figure 24:**
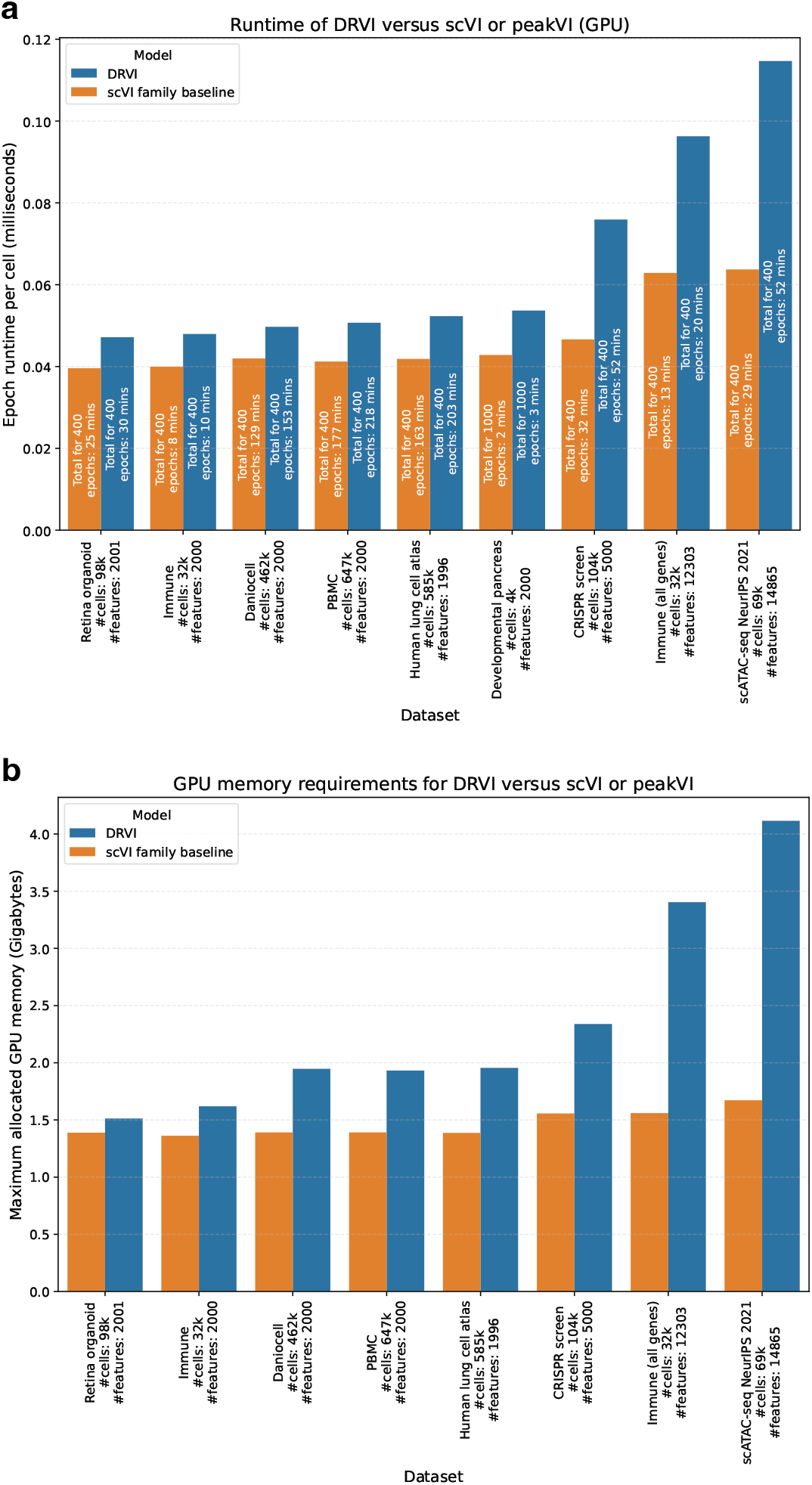
Runtime analysis of DRVI. **a**,**b**, The runtime (a) and maximum allocated GPU memory (b) for DRVI, scVI, and peakVI are shown for each benchmarked dataset. To normalize for varying dataset sizes and training lengths, the y-axis in (a) represents the runtime divided by the number of cells and the number of epochs (runtime per sample per epoch). The total runtime for each experiment is also displayed on the respective bars. DRVI’s training runtime is on the same order as scVI or peakVI up to a small scaling factor, with a maximum twofold increase observed in our experimental setups. In addition, training DRVI remains manageable when applied to the full gene set.

**Supplemental Figure 25:**
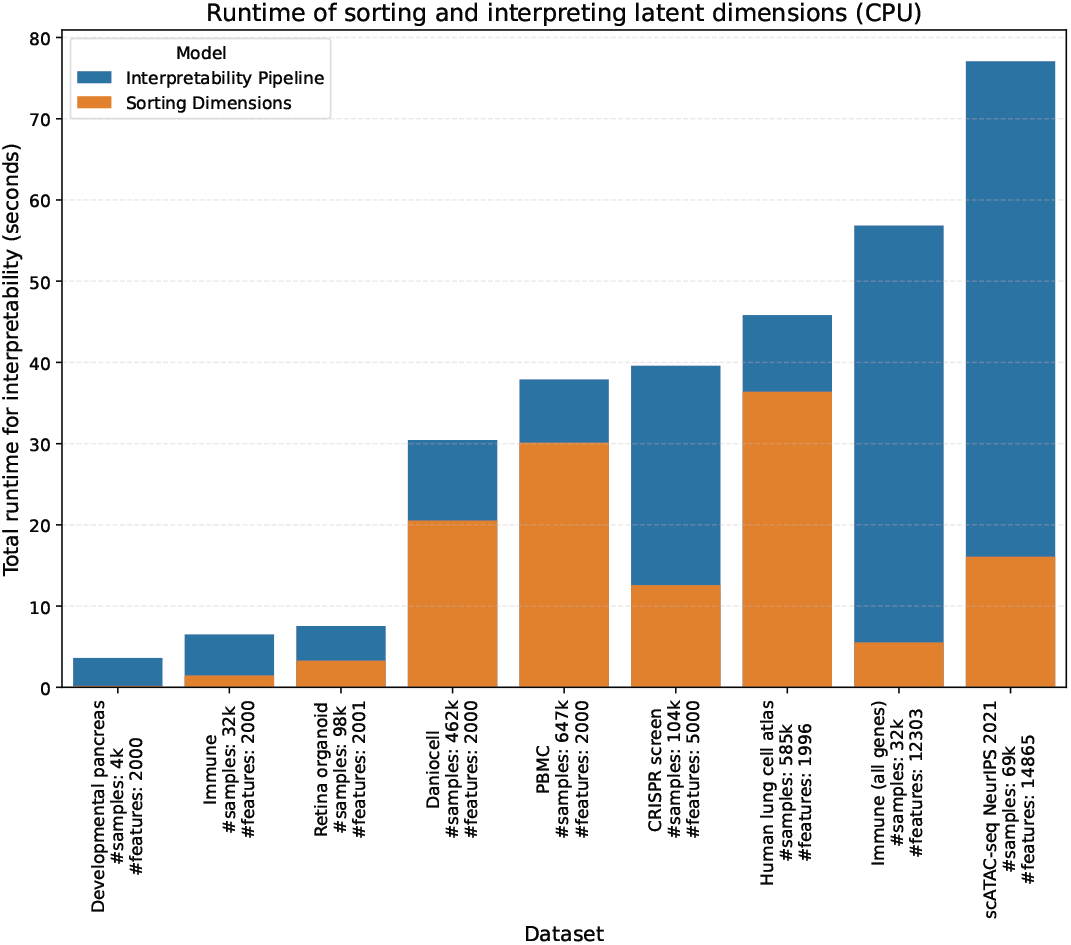
Runtime for interpretability and sorting latent factors. The time required to sort the latent factors and interpret gene programs for all latent factors is calculated. This demonstrates that the interpretability runtime is negligible.

**Supplemental Figure 26:**
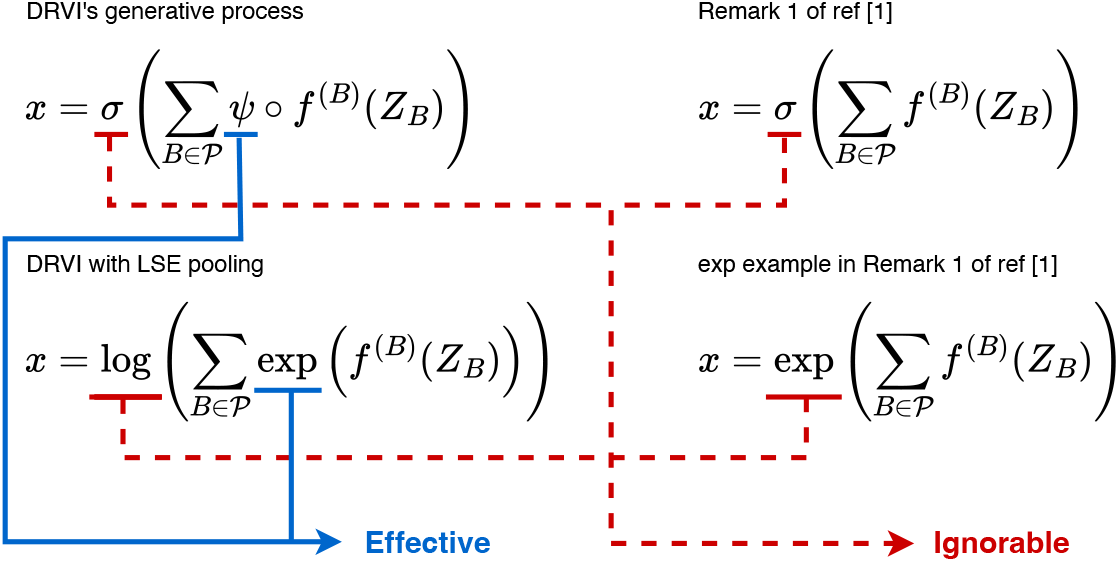
The effective and ignorable parts of the pooling function. Remark 1 from [34] suggests that activation functions applied after summing additive decoders can be disregarded. In contrast, here, we make use of a shared activation function (*ψ*) before the sum operator, which is not ignorable. We practically observe that this novel constraint plays a crucial role in enforcing disentanglement.

**Supplemental Figure 27:**
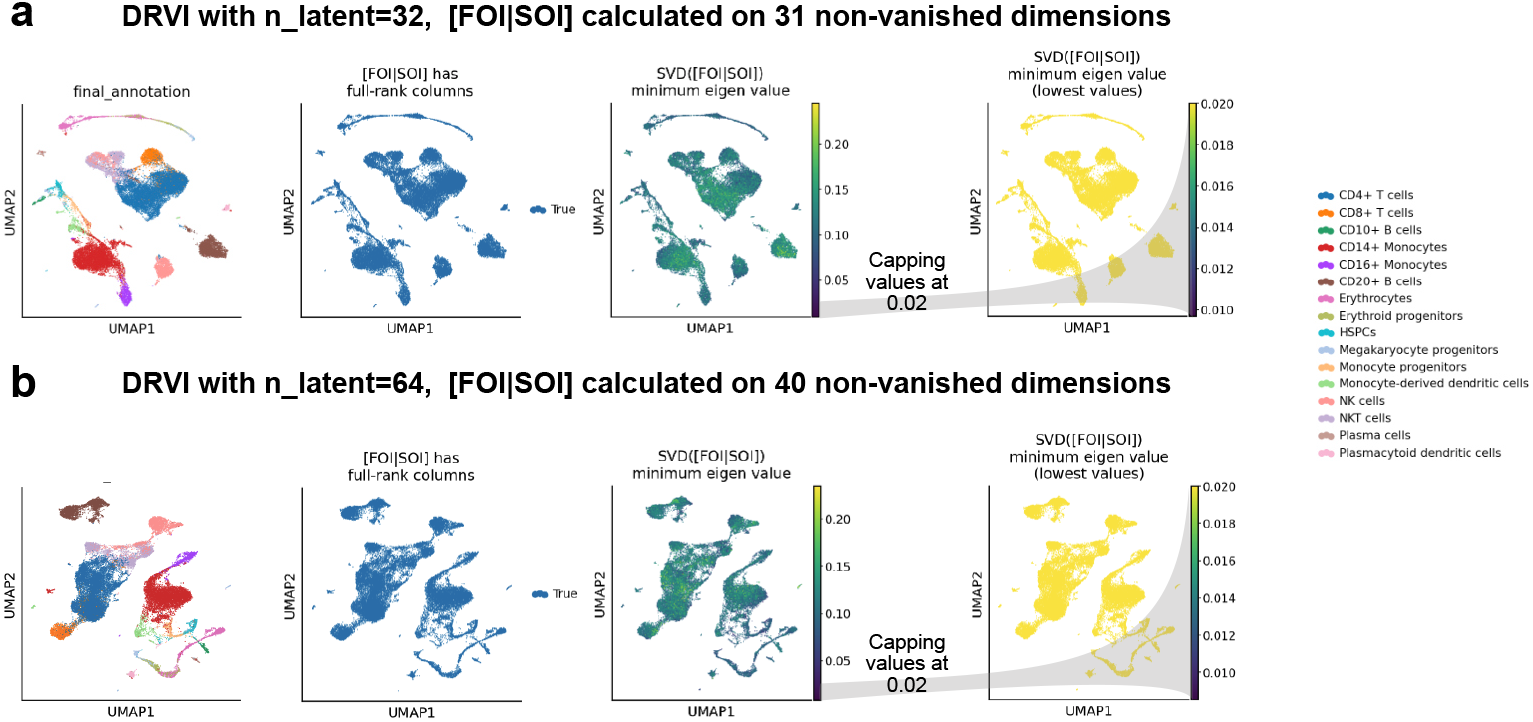
Numerical validation of sufficient independence in Immune data. **a**,**b**, The calculation of the rank of the sufficient independence matrix ([*FOI*|*SOI*]) on training data for DRVI models with (a) 32 and (b) 64 latent dimensions. The plot shows UMAP colored by cell types. The plot illustrates the status of the sufficient independence matrix being of full rank, the minimum eigenvalue of the SVD of this matrix after column-wise normalization, and the lower end of the minimum eigenvalues, demonstrating that the minimum values in both cases are more than zero (approximately 0.01) which indicates the matrices are numerically full-rank.

**Supplemental Figure 28:**
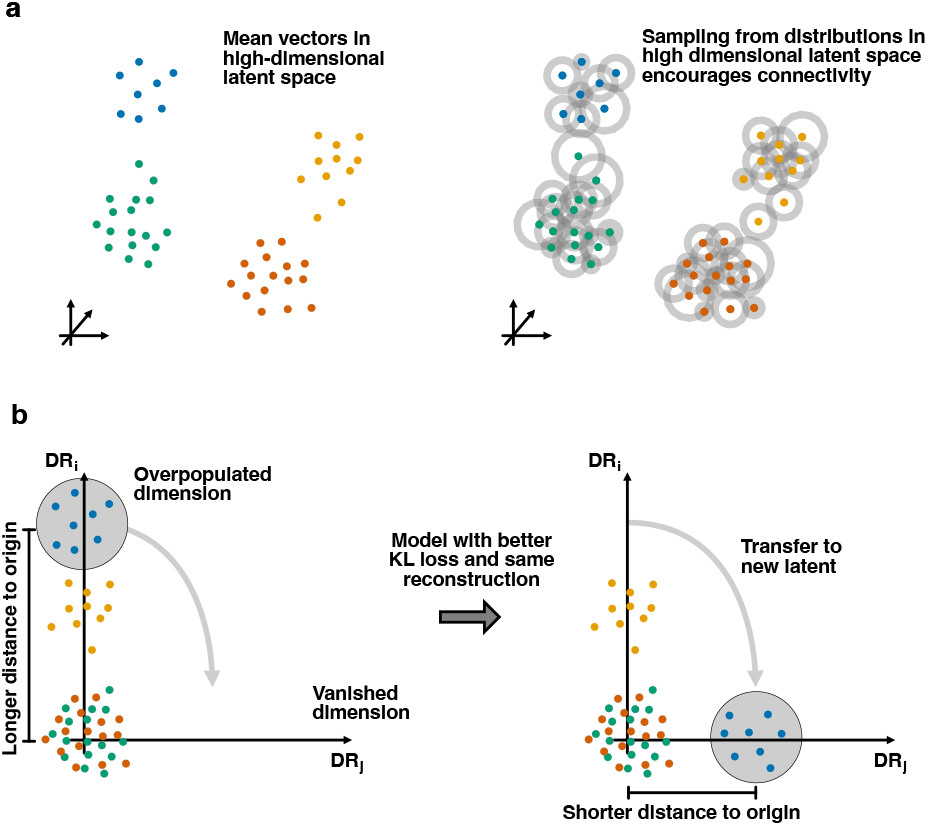
How latent space sampling and KL loss promote global disentanglement. **a**, In DRVI, like other VAE-based methods, the latent representation of each cell isn’t a vector, but rather a distribution, typically represented by a multivariate Gaussian with mean *µ* and a diagonal covariance matrix derived from a *σ* vector. The KL loss term encourages *µ* values to be near zero and *σ* values to be near one. As the model converges, these non-negligible *σ* values cause the distributions of neighboring cells to overlap. This prevents the model from indexing individual cells and promotes a continuous manifold of nearby cells. Thus, the KL loss term encourages path connectivity. **b**, Assuming the model has achieved local disentanglement and a sufficient number of latent dimensions are available, the KL loss term indirectly encourages irrelevant processes to be represented in individual latent factors. If two irrelevant processes are present within a single latent factor and an additional dimension is available, further optimization of the model is possible. By relocating the process further from the center to an available dimension, one can achieve the same reconstruction quality while simultaneously improving the KL loss.

**Supplemental Figure 29:**
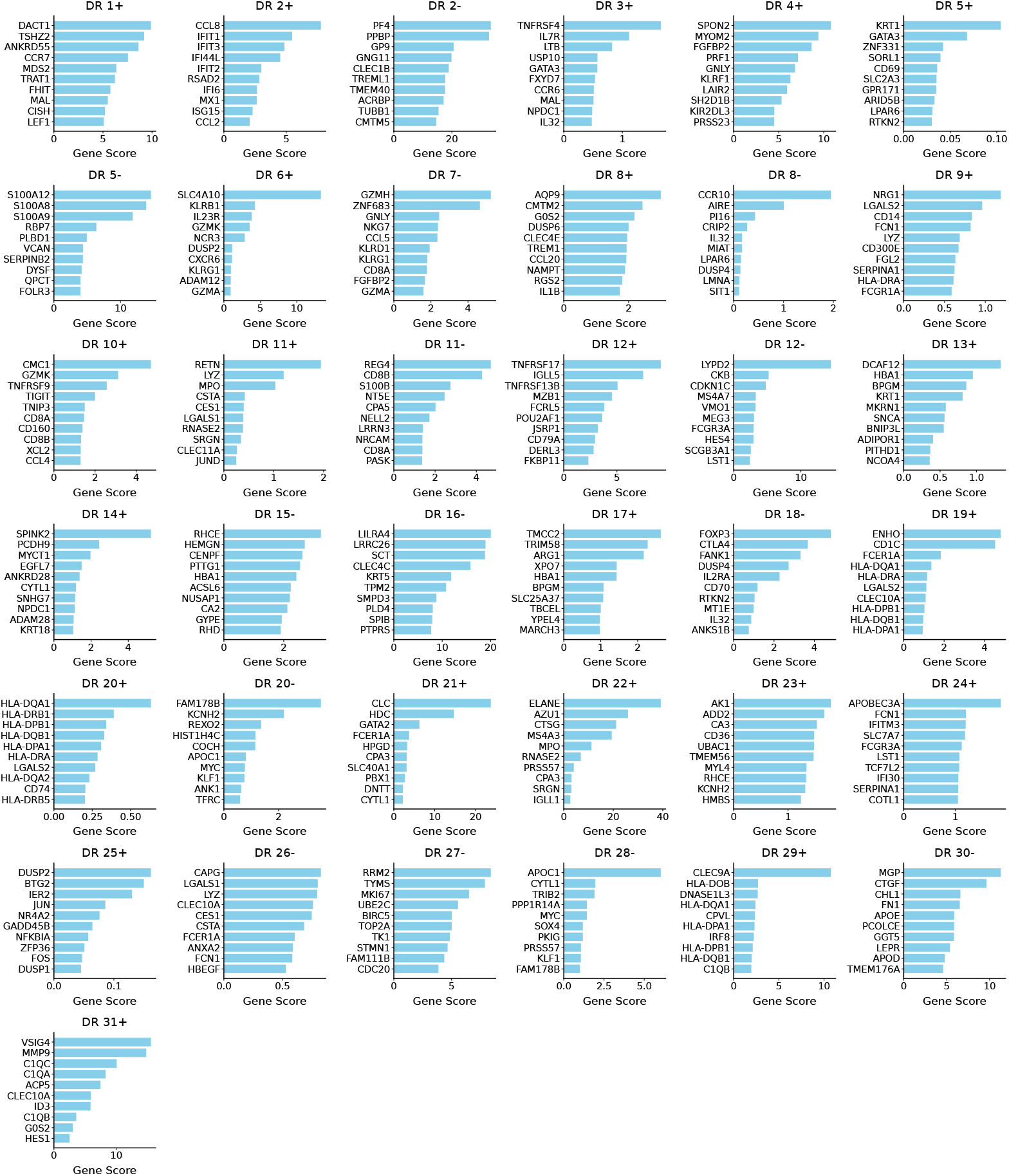
Interpretability of DRVI latent factors in immune dataset. The top relevant genes and their scores are plotted for each latent factor of DRVI trained on the immune dataset.

**Supplemental Figure 30:**
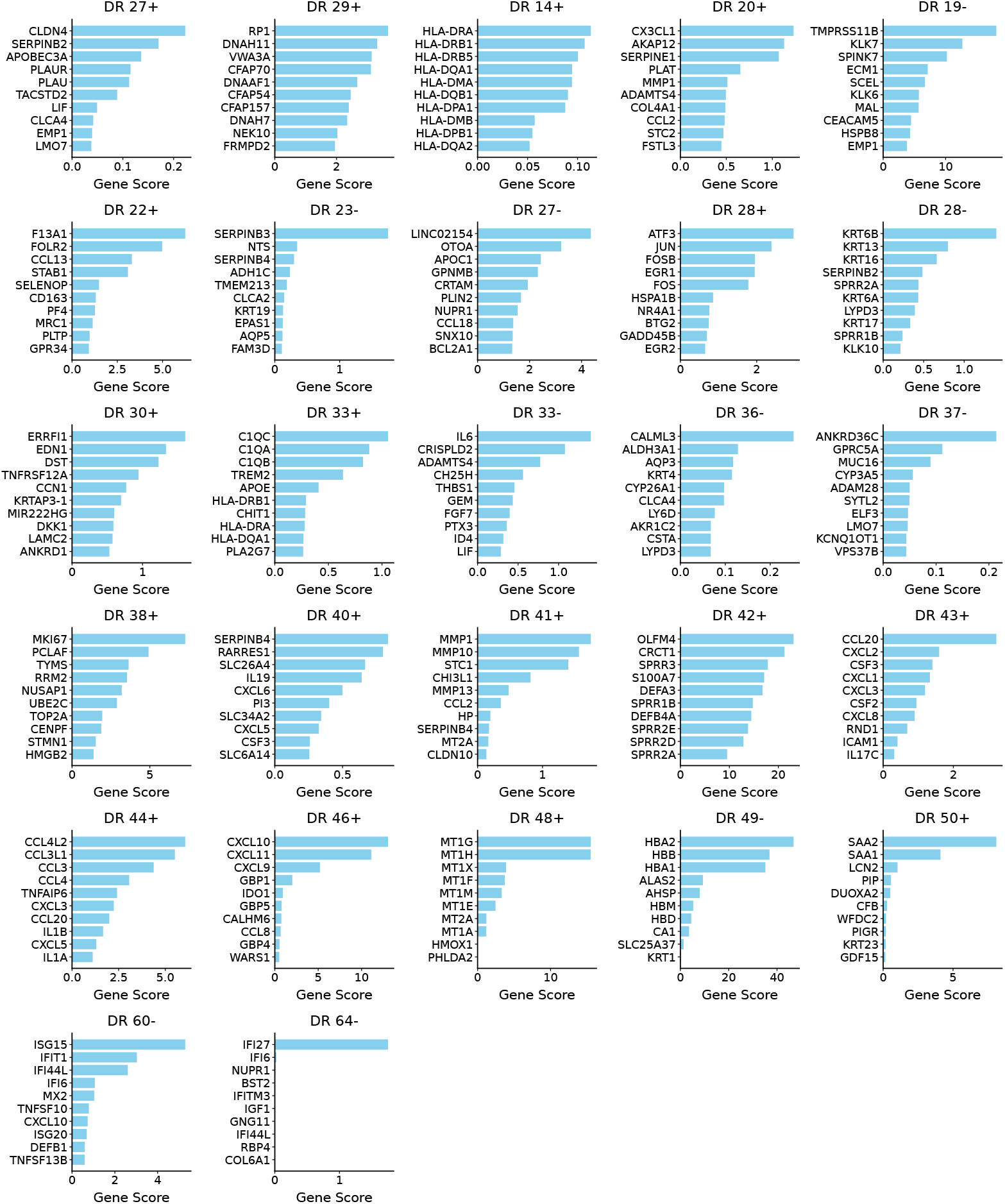
Interpretability of DRVI latent factors corresponding to non-cell-type processes in the HLCA. The top relevant genes and their scores are plotted for each non-cell-type latent factor of DRVI trained on the HLCA.

**Supplemental Figure 31:**
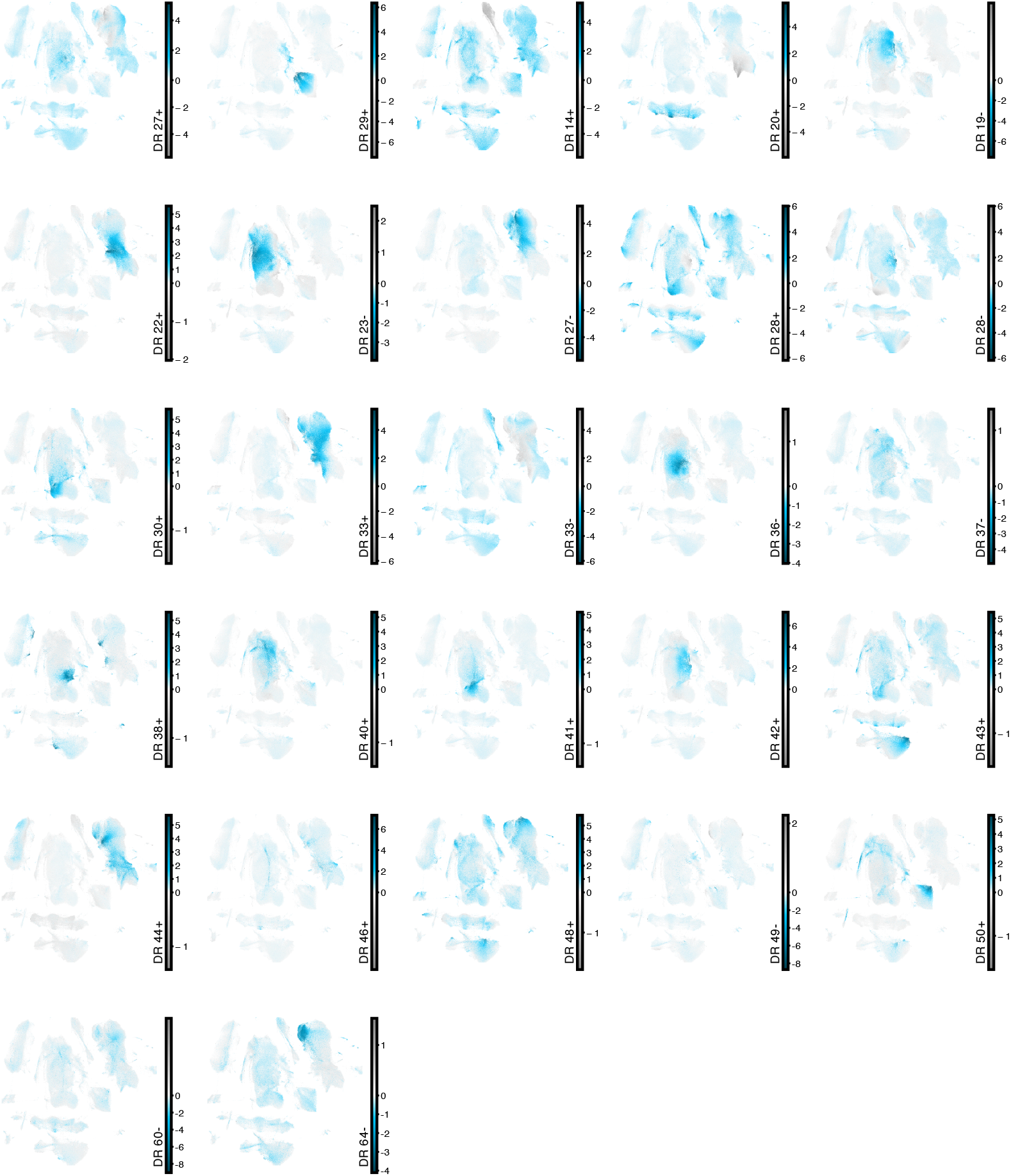
Activity of DRVI factors representing non-cell-type processes in the HLCA on UMAP. For factors representing non-cell-type processes, the activity on the UMAP is provided.

